# Dismantling Chromosomal Stasis Across the Eukaryotic Tree of Life

**DOI:** 10.64898/2026.04.14.718287

**Authors:** Megan Copeland, Meghann McConnell, Andres Barboza, Hannah M. Abraham, James Alfieri, Steven Arackal, Carrie E. Bernard, Kiedon Bryant, Shelbie Cast, Sean Chien, Emily Clark, Cassandra E. Cruz, Aileen Y. Diaz, Olivia Deiterman, Riya Girish, Kaya Harper, Carl E. Hjelmen, Michelle J. Thompson, Rachel Koehl, Tanvi Koneru, Kenzie Laird, Yoonseo Lee, Virginia R. Lopez, Mallory Murphy, Nayeli Perez, Gideon H. Schlab, Sarah Schmalz, Terrence Sylvester, Heath Blackmon

## Abstract

Chromosome number shapes genome organization, recombination, and speciation, yet how fast it evolves across the tree of life has never been measured. We analyzed 63,682 karyotypes across 55 eukaryotic clades and found that dysploidy rates vary by 844-fold, from approximately 0.0008 to 0.7 events per million years. This variation does not follow kingdom boundaries or deep phylogeny; intraclade variance exceeds interclade differences by more than an order of magnitude. Even birds, the textbook example of chromosomal stasis, exceed the global median rate once microchromosome dynamics are resolved. Contrasting the stasis of Odonata with the volatility of Orchidaceae reveals that life history and population structure, rather than deep phylogenetic constraints, govern the tempo of karyotypic change.

---

How fast do chromosome numbers evolve? The question sounds simple but has never been answered on a global scale. Chromosome number is a defining feature of genome architecture, setting the physical boundaries within which heritable variation is organized and transmitted. Whether through the incremental shifts of dysploidy or the saltational leaps of polyploidy, karyotypic change reshapes genome organization, recombination landscapes, and gene regulation (*1*–*3*). Beyond gene expression, structural karyotypic divergence drives speciation. By inducing meiotic mismatches in hybrids, chromosomal rearrangements erect reproductive barriers that catalyze lineage divergence (*4, 5*). These barriers facilitate lineage separation and, particularly in plants, drive bursts of diversification (*6, 7*). The number of chromosomes also dictates the recombination landscape: many small chromosomes accelerate allelic shuffling, whereas fewer, larger chromosomes constrain haplotype diversity and, by extension, the efficiency of natural selection (*8*).

For nearly a century, the prevailing paradigm in cytogenetics has been one of evolutionary stasis, a conviction that chromosome numbers are inherently conservative, changing at a glacial pace across most lineages (*6, 7*). This view has long been bolstered by seemingly pervasive empirical support. In birds, the ancestral karyotype appears frozen: over 60% of extant species cluster within 2n = 74–86, a pattern long interpreted as deep-time conservation (*9, 10*). Similarly, *Drosophila* has maintained six fundamental linkage groups for over 60 million years despite pervasive internal rearrangement, reinforcing the impression that chromosomal organization resists change (*11*).

While these flagship clades have shaped our theoretical foundations, the absence of a global, cross-kingdom framework has obscured the universal tempo of chromosomal evolution. Here, we close this gap by analyzing the broadest cytogenetic dataset ever assembled: 63,682 karyotypes spanning 56 major eukaryotic clades (Fig. 1). A course-based undergraduate research experience (CURE) coupled with AI-augmented literature discovery and expert validation enabled us to sample the full phylogenetic breadth of animals, plants, and fungi. With a unified Bayesian framework and consistent time-calibration across all phylogenies, we provide the first global metric of the rate of chromosomal evolution. Our results reveal that karyotypic change is not a conservative background process, but a highly volatile feature of the genome: dysploidy rates vary by 844-fold across the Tree of Life, with a global median rate of approximately one event every five million years.

**Figure 1.**
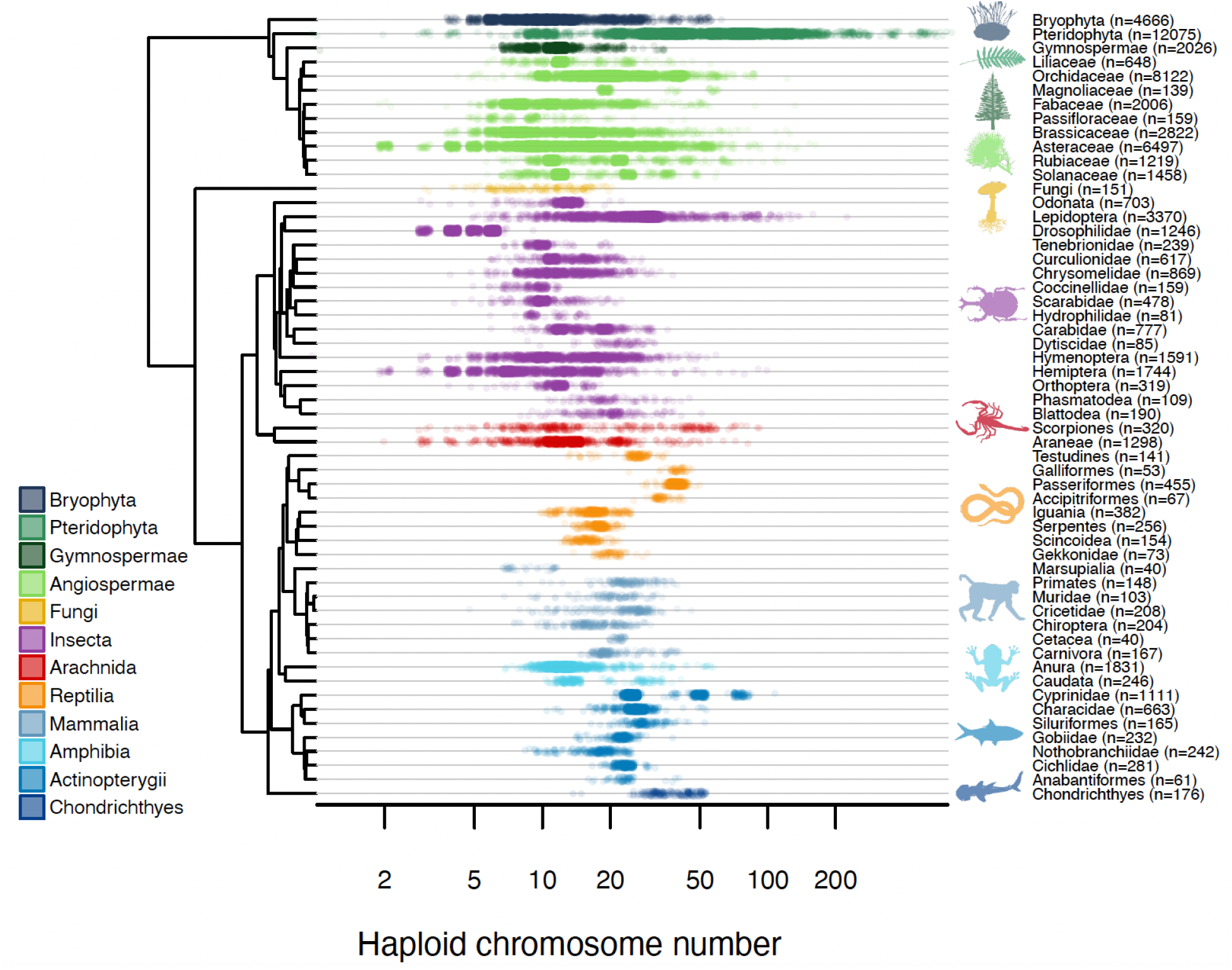
Chromosome number variation across the eukaryotic tree of life. Distribution of haploid chromosome counts compiled from 63,682 karyotypes spanning 56 monophyletic clades across plants, fungi, and animals. Each point represents a single species, plotted within its focal clade and colored by major taxonomic group. Chromosome number varies widely both within and among clades, spanning more than two orders of magnitude, and this variation is present across all major eukaryotic lineages rather than being restricted to any single kingdom.

## The Global Tempo of Chromosomal Change

Our analysis integrates 63,682 karyotypes across 55 monophyletic eukaryotic clades (excluding Magnoliaceae due to poor phylogenetic scaffolding) to provide a unified metric of chromosomal evolution across the tree of life. We find that dysploidy rates are remarkably heterogeneous (Fig. 2). This variation does not track major phylogenetic divisions or kingdom-level boundaries. Instead, the highest angiosperm rates overlap with those of the most volatile mammalian and insect lineages, while the slowest representatives of each kingdom are statistically indistinguishable from one another, all experiencing minimal chromosomal turnover. A formal variance components analysis confirmed that kingdom-level identity (Animal, Plant, Fungi) explains none of the observed variation in dysploidy rates (ANOVA F_2,52_ = 0.48, p = 0.62; REML ICC = 0.000). Even at the finer level of higher taxonomic classification (e.g., Mammalia, Insecta, Angiospermae), group identity accounts for only 6.2% of the variance; the remaining 93.8% falls among clades within groups (Fig. S69). The geometric mean dysploidy rates for animals and plants are nearly identical (both ∼0.023 events/Myr), yet rates within each kingdom span more than two orders of magnitude. These results indicate that the tempo of karyotype evolution is governed primarily by lineage-specific biological traits and ecological context rather than deep phylogenetic history.

**Figure 2.**
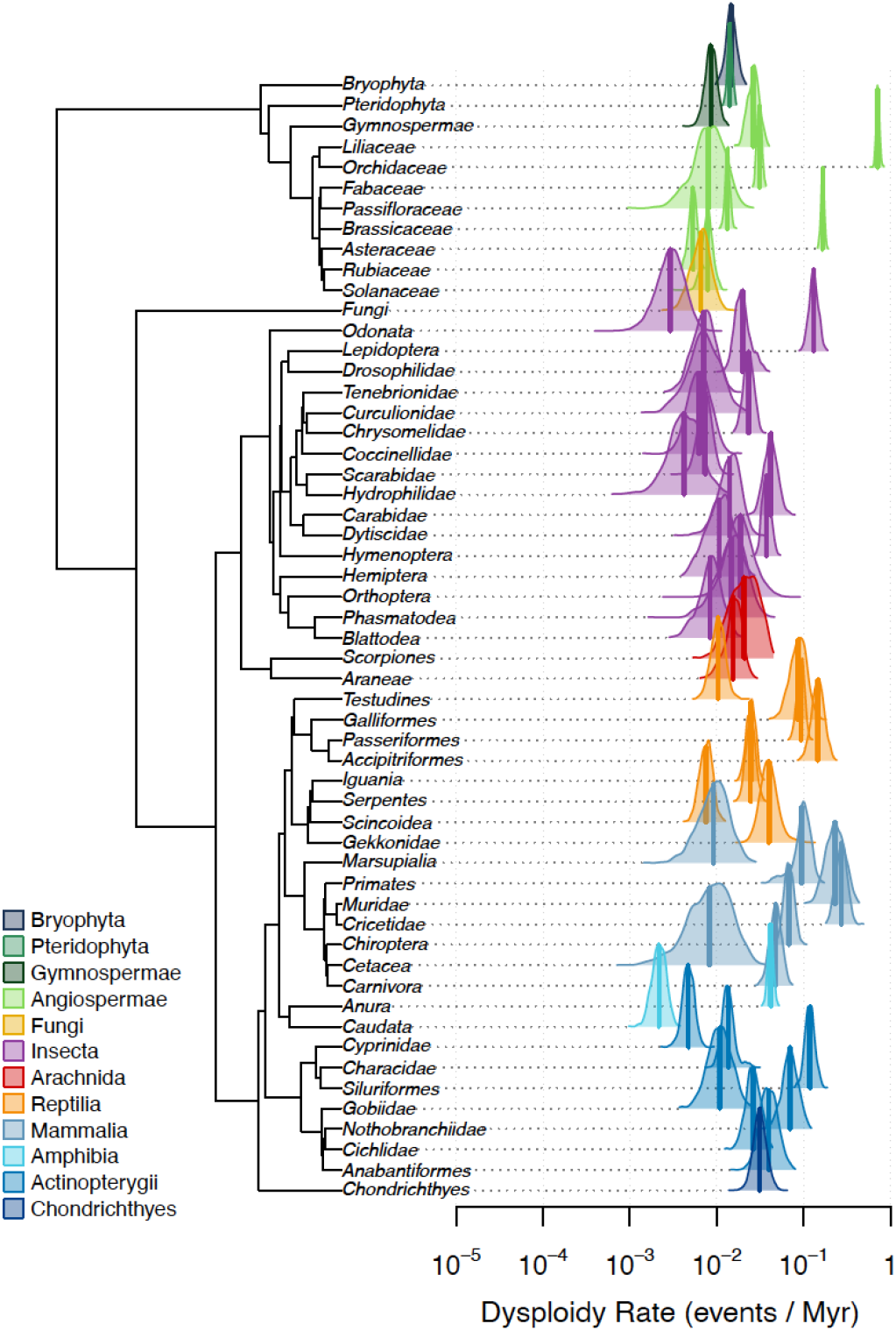
Probability distributions of dysploidy rates across the eukaryotic tree of life. Posterior probability densities of dysploidy rates (events per million years) estimated for 55 monophyletic clades spanning plants, fungi, and animals. Each distribution summarizes uncertainty in the inferred rate for a focal clade, with vertical lines indicating posterior medians. Dysploidy rates vary by 844-fold, yet substantial overlap is observed among major taxonomic groups. Notably, clades traditionally considered to exhibit rapid chromosomal evolution, such as mammals, show posterior density peaks that overlap extensively with those of angiosperms, indicating that similar evolutionary tempos arise independently across kingdoms.

To distinguish biological signals from methodological artifacts, we conducted extensive sensitivity analyses. We first evaluated the influence of prior specification and found that median posterior rate estimates were robust, with strong concordance between exponential and uniform priors. We then assessed how tree size and taxonomic sampling affected inference, observing that reduced sampling increased posterior uncertainty but did not bias estimated rate magnitudes. Finally, posterior predictive simulations of chromosome number variance and entropy confirmed that our models captured the key features of the observed data, validating the framework for global comparison.

## Dismantling the Paradigm: The Resolution of Avian Stasis

This global framework directly challenges the long-standing paradigm of lineage-specific stasis, a central tenet of cytogenetics for nearly a century. Nowhere is this challenge more apparent than in birds, historically the exemplar of chromosomal conservatism owing to the prevalence of the 2n = 74–86 ancestral karyotype (*9, 10*). Our results reveal that this perceived stasis is largely an artifact of limited methodological resolution and the absence of cross-clade rate estimates. Historical surveys were frequently blind to the dynamics of microchromosomes (*12*–*14*), which constitute roughly half of the avian genome (*15, 16*). By incorporating all chromosome transitions, including microchromosomes, into a unified phylogenetic framework, we find that every avian order in our dataset exceeds the global median dysploidy rate.

Rather than being frozen in deep time, the avian karyotype undergoes measurable, ongoing turnover. Avian karyotypic stability, then, reflects not an intrinsically constrained chromosomal architecture but the inability of earlier methods to resolve the most dynamic components of the karyotype. When compared to truly static lineages like Odonata, which likely maintain stability through nearly panmictic populations, birds emerge as active participants in chromosomal evolution. Karyotypic change is a volatile feature of the genome, driven by local ecology, not phylogenetic fate.

## Ecology and Demography, Not Architecture, Sets the Tempo

The vast heterogeneity in dysploidy rates suggests that the pace of karyotypic change is not stochastic but instead reflects the convergence of specific biological traits. Examining lineages at the extremes of our distribution reveals how genomic architecture, reproductive biology, and ecological context converge to accelerate or arrest chromosomal evolution.

At the high-velocity extreme, Orchidaceae exhibits the fastest dysploidy rates in our dataset, a result that appears paradoxical given the family’s monocentric chromosomes. Classical theory predicts strong constraint in orchids because their monocentric chromosomes should impose fitness costs on fissions and fusions (*17*–*19*). Such architecture should favor karyotypic stability, not volatility. Yet orchids repeatedly violate this expectation. Rather than preventing rearrangements, several features of orchid reproductive biology appear to buffer the fitness costs of chromosomal change. Pollinium-based reproduction generates extreme reproductive skew: most individuals fail to reproduce, but a single successful pollination event fertilizes an entire seed complement, amplifying the transmission probability of rare chromosomal variants (*20*). Widespread self-compatibility and vegetative propagation further reduce fitness costs for rare chromosomal variants, allowing them to reproduce without outcrossing (*21*). Coupled with fragmented habitats and small local effective population sizes, these traits create conditions under which chromosomal variants can escape purifying selection and drift to fixation, despite intrinsic monocentric constraints (*22*).

In stark contrast, Odonata exhibit profound karyotypic stasis despite possessing holocentric (diffuse) chromosomes, the very architecture predicted to allow fission–fusion dynamics. Under the holocentric model, chromosome fragments can retain centromeric activity and should segregate properly, so stasis in odonates cannot be attributed to the classic monocentric hazards of acentric loss or dicentric breakage (*7, 18*). That odonates nevertheless show long-term chromosomal stability indicates that centromere structure alone cannot explain evolutionary tempo. Instead, their stasis is best explained by demographic and ecological constraints. As highly mobile, obligately outcrossing aerial predators, odonates maintain large effective population sizes and extensive gene flow (*23, 24*), conditions under which novel rearrangements are efficiently purged by selection before they can spread (*25*).

Together, the contrasting karyotypic patterns in Orchidaceae and Odonata challenge the idea that centromere type sets a “speed limit” on chromosome evolution (Fig. 3). This empirical decoupling of centromere structure from evolutionary rate echoes broader comparative evidence: a phylogenetic analysis of insects found no consistent difference in dysploidy or polyploidy rates between holocentric and monocentric orders (*26*). If centromere architecture does not govern the tempo, what does? Population genetic theory predicts that small effective population sizes should allow underdominant rearrangements to drift to fixation more readily, yet the proxies available to test this prediction yield an inconsistent signal. Genome size, often invoked as a correlate of effective population size (*27*), does not predict dysploidy rate across clades (PGLS, p = 0.41; Fig. S63). Genome-wide dN/dS, an independent proxy for the efficacy of selection, shows a marginally significant positive association with dysploidy rate in animals (slope = 0.871, p = 0.059; Figs. S65, S67), suggesting that drift may contribute to chromosomal change in some lineages. Geographic range, however, shows no relationship (Fig. S66). The mixed signal across these proxies reinforces a central conclusion: no single parameter governs the tempo of chromosomal evolution. Rather, the rate of karyotypic change emerges from the interaction of life history, demographic context, and genomic architecture, a combination that varies idiosyncratically across lineages.

**Figure 3.**
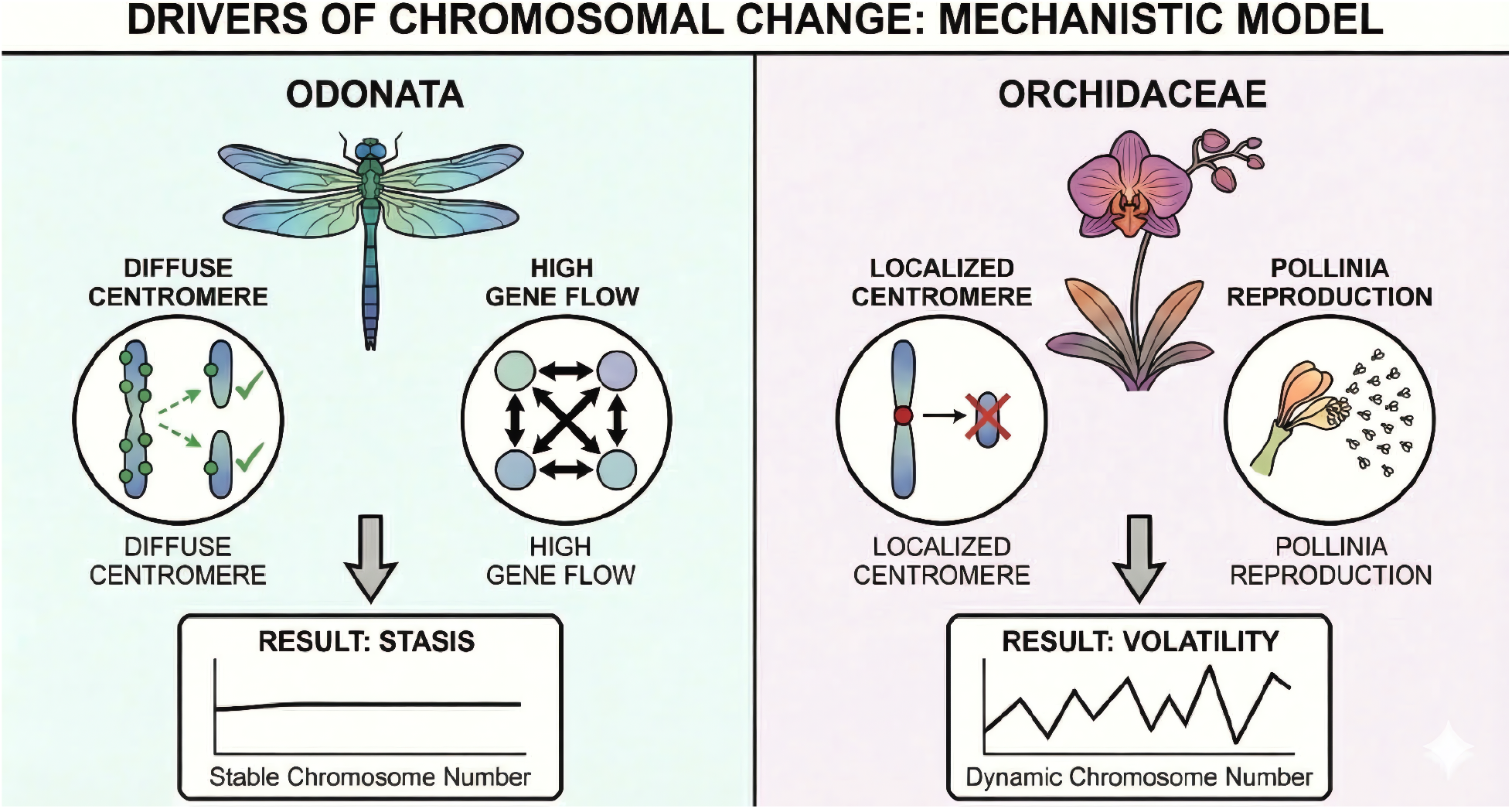
Drivers of chromosomal evolutionary tempo. Conceptual model showing how genomic architecture, reproductive biology, and demography interact to shape dysploidy rates. In Odonata (left), holocentric chromosomes permit rearrangements, but high gene flow and large effective population sizes lead to efficient purging of variants, resulting in karyotypic stasis. In Orchidaceae (right), despite monocentric constraints, pollinium-based reproduction, selfing, and small, fragmented populations reduce purifying selection, allowing chromosomal variants to persist and fix, producing high karyotypic volatility. These contrasts illustrate that centromere type alone does not determine evolutionary rate; instead, tempo is driven by life history and population genetic context.

## Modes of Evolution: The Mechanistic Divide

While our results demonstrate that the tempo of chromosomal evolution is kingdom-agnostic, the underlying mechanisms driving these changes reveal a clear divide between major lineages. AIC-based comparisons of nested models reveal that the relative contributions of dysploidy and polyploidy vary predictably across the tree of life. Across the 55 clades examined, the fully parameterized model (accounting for both incremental dysploidy and saltational polyploidy) provided the best fit for 24 lineages. Comparisons between full and reduced models indicated strong support for including polyploidy in 27 of 55 clades (ΔAIC > 5).

This support was strongest in plant clades, where removing polyploidy parameters substantially reduced model fit. All plant clades showed this pattern, whereas only 16 of 43 animal clades showed strong support for polyploidy. This confirms that whole-genome duplication remains a primary driver of karyotypic diversity in plants, often providing raw material for subsequent adaptive radiation (*6, 28, 29*). In contrast, for many animal lineages, including most insects, mammals, and ray-finned fishes, the removal of polyploidy parameters had a negligible effect on model performance. In these groups, chromosome number diversity is driven primarily by dysploidy (but see (*30*)). These findings underscore that while different lineages may arrive at similar evolutionary tempos, they do so through distinct genomic pathways: plants through a mix of duplication and rearrangement, and animals primarily through rearrangement.

## Conclusion: A New View of the Eukaryotic Karyotype

By integrating nearly 64,000 karyotypes into a unified, time-calibrated framework, we show that chromosome evolution proceeds on shorter timescales and with greater heterogeneity than traditionally assumed. Our findings dismantle the century-old paradigm of chromosomal stasis, demonstrating that even “frozen” lineages like birds are active participants in karyotypic turnover when viewed through a broad taxonomic lens.

The discovery that chromosomal volatility varies by 844-fold, and is largely independent of deep phylogenetic history, suggests that chromosome number is a highly responsive trait. Rather than a conserved structural constant, the karyotype is shaped by local biological conditions. Its pace of change is dictated by the interaction between genomic architecture, such as centromere organization, and the population genetic environment, including mating systems and gene flow. When these factors reduce the fitness costs of structural change, the genome reshapes itself with remarkable speed.

The physical organization of the genome is not a passive trait but a dynamic feature of a lineage’s natural history. By replacing range-based inferences with explicit rate estimation, we reveal the eukaryotic karyotype for what it is: a volatile and essential participant in evolutionary diversification.

## Methods Summary

This study was conducted within a Course-based Undergraduate Research Experience (CURE) with 15 undergraduates. Chromosome number and phylogenetic data were assembled using AI-assisted literature discovery followed by rigorous human validation. A large language model (ChatGPT-5) was used solely to identify candidate sources, after which all chromosome counts and phylogenies were independently verified by undergraduate curators and reviewed by the first and last authors. This workflow produced a curated dataset of 63,682 karyotypes spanning 56 monophyletic eukaryotic clades. Chromosome number evolution was analyzed within a unified Bayesian framework using the chromePlus and diversitree packages in R (*31, 32*). Haploid chromosome counts were encoded as discrete states and modeled on time-calibrated ultrametric phylogenies that were scaled in a unified framework and pruned to match the chromosomal datasets. We estimated independent rates of chromosome gain, loss, polyploidy, and demiploidy via MCMC (1,000 steps; all ESS > 200) with exponential priors; sensitivity analyses used uniform priors. Phylogenetic uncertainty was incorporated by repeating inference across posterior tree sets when available. Model adequacy was evaluated using posterior predictive simulations of chromosome variance and entropy and by assessing effective sample sizes. We tested whether dysploidy rates differ among major taxonomic groups using hierarchical variance decomposition: one-way ANOVA and REML variance components models (lme4) at both the kingdom and higher taxonomic levels, with significance assessed by permutation tests (9,999 permutations). Support for alternative evolutionary modes was assessed via AIC-based comparison of nested maximum-likelihood models. All data used in this study are publicly available through an online database (karyotype.org).

## Acknowledgements

Large language models (ChatGPT-5) were utilized for the data collection and image generation for figure 3 aspect of this work. All work was independently verified by the first and last authors.

## Funding

National Institute of General Medical Sciences, National Institutes of Health grant R35 GM138098 (HB)

## Author contributions

Conceptualization: HB

Data Collection: MC, HMA, JA, SA, AB, CEB, KB, S. Cast, S. Chien, EC, CEC, AYD, OD, KH, CH, MJ, RK, TK, KL, Yl, VRL, MM, MM, NP, SS, TS, HB

Analysis: MC, HMA, SA, AB, CEB, KB, SC, SC, EC, CEC, AYD, OD, KH, RK, TK, KL, Yl, VRL, MM, MM, NP, SS, HB

Visualization: MC, HMA, SA, AB, CEB, KB, SC, SC, EC, CEC, AYD, OD, KH, RK, TK, KL, Yl, VRL, MM, MM, NP, SS, HB

Supervision: MC, AB, S. Chien, HB

Writing – original draft: MC, HB

Writing – review & editing: MC, HB

## Competing interests

The authors declare that they have no competing interests.

## Data, code, and materials availability

All data and scripts used for this project are publicly available via GitHub (https://github.com/coleoguy/metazoa-chromosomes) and are permanently archived on Zenodo (https://doi.org/10.5281/zenodo.19559664).

## Supplementary Materials

The supplemental PDF contains:

Materials and Methods

Tables S1-S4

Figures S1-S63

References 33-44

## Notes

https://github.com/coleoguy/metazoa-chromosomes

https://doi.org/10.5281/zenodo.19559664

### Competing Interest Statement

The authors have declared no competing interest.

### Summary of Updates

The author list and affiliations have been updated.

