## Supplemental Info for "Dismantling Chromosomal Stasis Across the Eukaryotic Tree of Life"

### SUPPLEMENTARY MATERIALS

Table of Contents


1. Extended Methods
2. Data tracking and characteristics
3. MCMC Convergence
4. Mentor vs Mentee results
5. Prior Influence
    a. Exponential vs Uniform
    b. Interactions between tree size and prior
6. Model Comparison
7. Clade Audits
8. Tree Resolution Effects on Dysploidy Rates
9. Genome Size Effects on Dysploidy Rates
10. Variance of Haploid Chromosome Number (Plants vs. Animals)
11. Effective Population Size Effects on Dysploidy Rates
12. Variance Decomposition: Within- vs. Between-Group Variation in Dysploidy Rates
13. Online Database

### Extended Methods

*Study Design and Distributed Curation Network*

To achieve global sampling of eukaryotic chromosomal evolution, we compiled chromosome count and phylogenetic datasets using a structured data curation network. This network consisted of 20 undergraduate researchers, six graduate student mentors, and three faculty collaborators, enabling simultaneous sampling across animals, plants, and fungi. Each analyst was assigned two non-overlapping monophyletic clades selected to maximize phylogenetic breadth and avoid overlapping clades.

Preliminary literature discovery and data aggregation was accelerated using large language models (LLMs; specifically OpenAI's ChatGPT-5). Analysts used standardized prompts to generate candidate sources for phylogenies and chromosome data, including data repositories (e.g., Dryad, Figshare, GitHub) and primary publications. All LLM-derived outputs were treated strictly as leads and were subject to verification. Each chromosome count and phylogeny was manually validated against the published original source. Data acquisition proceeded under a two-step quality control procedure.

1. **Source Validation:** Analysts reviewed each primary source to verify the reported chromosome count, taxonomic validity, and the suitability of the phylogeny for comparative analysis.
2. **Expert Review:** Completed datasets were reviewed by the first author and the supervising faculty member. These reviews verified extraction accuracy and checked that phylogenies met criteria for completeness, time calibration, and clade structure.

*Data Collection and Standardization*

**Chromosomal Data**: For each target clade, we compiled a dataset of haploid chromosome numbers (n). To standardize reports containing ranges (e.g., "n=6–8") or polymorphisms, we implemented a uniform cleaning procedure: integer values were sampled uniformly from reported ranges, and single values were drawn at random from discrete polymorphic sets. Non-integer values (e.g., from averaging of counts from numerous cells) were stochastically rounded based on their fractional components to generate integer input for discrete state models.

**Phylogenetic Data:** We prioritized ultrametric, time-calibrated phylogenies with high taxon sampling. Trees were first pruned to match the species present in the chromosomal datasets. For plant lineages, trees were sourced from the Smith & Brown (2018) megaphylogeny (*33*). Although this source is time-calibrated, numerical precision issues occasionally result in non-ultrametric attributes; we therefore re-smoothed these trees using non-negative least squares (nnls.tree from the phangorn package) (*34*). Some of the methods require strictly bifurcating trees so any polytomies were resolved using the multi2di function in ape (*35*). Any negative branch lengths were replaced with a minimal positive constant ($1 \times 10^{-6}$). For any other clades with non-ultrametric trees, we transformed branch lengths using penalized likelihood via the chronos function in the R package ape (*35*). Trees were scaled to unit height prior to model fitting.

*Bayesian Rate Estimation*

We modeled the evolution of chromosome number as a continuous-time Markov process using the chromePlus package in R. Chromosome data was converted to a discrete-state matrix format required by diversitree using datatoMatrix (chromePlus), with a buffer of one state beyond the observed min/max (*31*). We constructed a multi-state Markov model likelihood function using make.mkn (diversitree) with ODE-based likelihood evaluation and the strict flag set to false (this allows for unsampled chromosome states to be present in the model) (*32*). We then constrained the likelihood to a biologically realistic model of chromosome evolution using constrainMkn (chromePlus), allowing for independent rates of chromosome gain (ascending dysploidy), loss (descending dysploidy), demiploidy (increases by 1.5x the chromosome number) and polyploidy (increase by 2x the chromosome number). For clades in which the observed chromosome state space did not support explicit estimation of polyploidy transitions, we increased the state buffer used during likelihood construction to ensure that the effective state space allowed for the estimation of polyploidy in all clades.

To quantify parameter uncertainty, we estimated rates using Bayesian Markov Chain Monte Carlo (MCMC) via the mcmc function in diversitree (*32*). Chains were initialized from random starting values between zero and one drawn independently for each free parameter and then run for 1,000 generations. In the primary analyses, we applied an exponential prior to all rate parameters to reflect the biological expectation that chromosomal changes are generally rare events and constrain the posterior distribution in instances of low information content. In sensitivity analyses, we replaced the exponential prior with a uniform prior on each rate parameter, assigning equal prior probability to all rates between 0 and 20 events per unit tree length. We held all preprocessing steps, likelihood construction, and MCMC settings the same as in the exponential runs. This allowed us to assess the extent to which posterior estimates were driven by the likelihood rather than being a function of an excessively informative exponential prior.

For clades with posterior distribution of phylogenetic trees, MCMC inference was repeated independently across trees. Posterior samples from individual tree-specific runs were combined into a single posterior distribution, with an explicit tree identifier retained for each sample, thereby accounting for phylogenetic uncertainty rather than making inferences based on a single topology and set of branch lengths.

MCMC inference was performed on unit-length trees to facilitate interpretability of initial estimates, However, reported rates have been transformed into units of millions of years by dividing the posterior rate estimate by the tree depth. Median divergence estimates were obtained from the Time Tree of Life for taxa that subtended the root of the tree. This step allows us to use one global time calibration dataset across all clades allowing for directly comparable rate estimates across diverse lineages.

*Computational Validation and Model Adequacy*

To ensure strict computational reproducibility and rule out operator error during the analysis phase, the first and final authors developed a parallel validation pipeline. This automated script accepted the curated raw inputs (chromosome count .csv files and corresponding phylogeny .tre or .nex files) and independently re-executed the full analytical workflow for every clade. This included duplicate-tip removal from trees, ultrametric smoothing, pruning, likelihood construction, model constraint specification, and Bayesian MCMC inference using identical settings to those applied in the primary analyses. Posterior samples generated by the validation pipeline were compared directly to student-generated results by comparing the 95% HPD intervals. Discrepancies were flagged and resolved manually (Solanaceae and Anura) (Fig S1).

We employed a three-tiered validation framework to assess the reliability of dysploidy rate estimates for each of the 56 monophyletic clades:

1. **Prior–Posterior Overlap:** We calculated the overlap between the prior and posterior distributions of the dysploidy rate parameter. For each clade, we compared the posterior distribution of the total dysploidy rate (the sum of ascending and descending dysploidy rates) to the corresponding prior distribution. Overlap between the prior and posterior density curves was calculated, and clades exhibiting greater than 70% overlap were flagged as cases in which posterior estimates were weakly informed by phylogenetic signals (Table S4; Figs S5-S60).
2. **Posterior Predictive Simulation (PPS):** We evaluated model adequacy using posterior predictive simulations implemented in chromePlus (*31*). We sampled 1,000 parameter draws from the post–burn-in posterior distribution of the fitted model. For each draw, we simulated chromosome number evolution on the empirical phylogeny using simChrom, using the fitted dysploidy, polyploidy, demiploidy rates when present in the posterior output (otherwise setting them to zero). Simulations were bounded to a plausible state range defined as 1 (biological constraint) to the maximum observed haploid count + 5. For each simulated dataset, we computed two summary statistics across tips: the variance of haploid chromosome numbers and Shannon's entropy of the chromosome number distribution. We compared the empirical variance and entropy to the posterior predictive distributions of these statistics. Clades were flagged for model inadequacy when the empirical statistic fell in the outer 5% of the simulated distribution.
3. **Effective Sample Sizes:** We further evaluated the robustness of inference by assessing MCMC mixing and reproducibility across independent runs. Effective sample sizes (ESS) were calculated for all estimated parameters (after removal of the first 100 steps as burn-in) using the coda package (*36*). Chains with low ESS values or parameters exhibiting near-zero variance were flagged for additional inspection.

*Sensitivity Analysis of Taxon Sampling*

To verify that rate estimates were comparable across clades with vastly different sampling densities, we conducted both a simulation-based and an empirical subsampling analysis. For the simulation-based analysis, we simulated chromosome evolution on a large phylogeny and then pruned the tree to retain a small fraction of tips, mimicking the sparse sampling of our empirical datasets. We then inferred rates on the pruned trees using the same Bayesian framework and priors described above. Posterior distributions inferred from full and pruned trees were compared using highest posterior density (HPD) interval overlap to assess the effect of reduced sampling. This process was repeated using our empirical Scarabaeidae dataset. These analyses demonstrated that our rate estimates remain robust to incomplete taxon sampling, validating comparisons between large, well-sampled clades and smaller or sparsely sampled clades.

*Model Comparison*

To evaluate support for alternative chromosome models, we performed a complementary model-selection analysis using maximum-likelihood estimation and Akaike Information Criterion (AIC). For each clade, we applied the same data preprocessing, phylogenetic preparation, likelihood construction steps as used in the Bayesian analyses. Four nested models were evaluated: a full model allowing ascending and descending dysploidy, polyploidy, and demi-polyploidy transitions; models excluding either polyploidy or demi-polyploidy; and a dysploidy-only model excluding both. Models were fit using maximum-likelihood optimization (find.mle) (*32*), with starting values initialized from posterior means of the Bayesian analyses, and compared using AIC. For clades represented by multiple phylogenetic trees, AIC values were calculated for each tree and averaged across trees. In some clades, one or more reduced model variants was not evaluated because likelihood optimization failed to converge.

#### Data tracking and characteristics

**Table S1. Clade Data Tracking Sheet.** Each row is one of the focal taxon that is analyzed in this manuscript. The citation column contains all papers and databases that were accessed to retrieve data (karyotypes or phylogenies).

| Clade Name | Student Leader | Karyotype records | Phylogeny tips | Ultrametric/ Bifurcating | Overlap | Karyotype and phylogeny sources |
|---|---|---|---|---|---|---|
| Accipitriformes | Aileen Diaz | 67 | 237 | T/T | 63 | Alfieri, James M., Kevin Bolwerk, Zhaobo Hu, and Heath Blackmon. 2024. “From Micro to Macro: Avian Chromosome Evolution Is Dominated by Natural Selection.” bioRxivorg. https://doi.org/10.1101/2024.11.29.626112.<br><br>Catanach, Therese A., Matthew R. Halley, and Stacy Pirro. 2025. “Enigmas No Longer: Using Ultraconserved Elements to Place Several Unusual Hawk Taxa and Address the Non-Monophyly of the Genus Accipiter (Accipitriformes: Accipitridae).” Biological Journal of the Linnean Society. Linnean Society of London 144 (2): blae028. |
| Passeriformes | Aileen Diaz | 455 | 9189 | T/T | 449 | Alfieri, James M., Kevin Bolwerk, Zhaobo Hu, and Heath Blackmon. 2024. “From Micro to Macro: Avian Chromosome Evolution Is Dominated by Natural Selection.” bioRxivorg. https://doi.org/10.1101/2024.11.29.626112.<br><br>Vertlife.org Database https://data.vertlife.org/ |
| Chondrichthyes | Andres Barboza | 176 | 1192 | T/T | 77 | Nagpure, Naresh Sahebrao, Ajey Kumar Pathak, Rameshwar Pati, Iliyas Rashid, Jyoti Sharma, Shri Prakash Singh, Mahender Singh, et al. 2016. “Fish Karyome Version 2.1: A Chromosome Database of Fishes and Other Aquatic Organisms.” Database: The Journal of Biological Databases and Curation 2016 (March): baw012.<br><br>Arai, Ryoichi. 2011. Fish Karyotypes: A Check List. 2011. Tokyo, Japan: Springer. |

| | | | | | | |
|---|---|---|---|---|---|---|
| | | | | | | Stein, R. William, Christopher G. Mull, Tyler S. Kuhn, Neil C. Aschliman, Lindsay N. K. Davidson, Jeffrey B. Joy, Gordon J. Smith, Nicholas K. Dulvy, and Arne O. Mooers. 2018. “Global Priorities for Conserving the Evolutionary History of Sharks, Rays and Chimaeras.” Nature Ecology & Evolution 2 (2): 288–98. https://vertlife.org/data/sharks/ |
| Pteridophytes | Andres Barboza | 12075 | 5868 | T/T | 1465 | Rice, Anna, Lior Glick, Shiran Abadi, Moshe Einhorn, Naama M. Kopelman, Ayelet Salman-Minkov, Jonathan Mayzel, Ofer Chay, and Itay Mayrose. 2015. “The Chromosome Counts Database (CCDB) - a Community Resource of Plant Chromosome Numbers.” The New Phytologist 206 (1): 19–26.<br><br>Nitta, Joel H., Eric Schuettpelz, Santiago Ramírez-Barahona, and Wataru Iwasaki. 2022. “An Open and Continuously Updated Fern Tree of Life.” Frontiers in Plant Science 13 (909768): 909768. |
| Carnivora | Carrie Bernard | 167 | 294 | T/F | 116 | Jonika, Michelle M., Kayla T. Wilhoit, Maximos Chin, Abhimanyu Arekere, and Heath Blackmon. 2024. “Drift Drives the Evolution of Chromosome Number II: The Impact of Range Size on Genome Evolution in Carnivora.” The Journal of Heredity 115 (5): 524–31.<br><br>Nyakatura, Katrin, and Olaf R. P. Bininda-Emonds. 2012. “Updating the Evolutionary History of Carnivora (Mammalia): A New Species-Level Supertree Complete with Divergence Time Estimates.” BMC Biology 10 (1): 12. |
| Caudata | Carrie Bernard | 246 | 796 | T/T | 204 | Perkins, Riddhi D., Julio Rincones Gamboa, Michelle M. Jonika, Johnathan Lo, Amy Shum, Richard H. Adams, and Heath Blackmon. 2019. “A Database of Amphibian Karyotypes.” Chromosome Research: An International Journal on the Molecular, Supramolecular and Evolutionary Aspects of Chromosome Biology 27 (4): 313–19.<br><br>Stewart, Alexander A., and John J. Wiens. 2025. “A Time-Calibrated Salamander Phylogeny Including 765 Species and 503 Genes.” Molecular Phylogenetics and |

| | | | | | | |
|---|---|---|---|---|---|---|
| | | | | | | Evolution 204 (108272): 108272. |
| Blattodea | Cassandra Cruz | 190 | 153 | T/T | 30 | Sylvester, Terrence, Carl E. Hjelmen, Shawn J. Hanrahan, Paul A. Lenhart, J. Spencer Johnston, and Heath Blackmon. 2020. “Lineage-Specific Patterns of Chromosome Evolution Are the Rule Not the Exception in Polyneoptera Insects.” Proceedings. Biological Sciences 287 (1935): 20201388.<br><br>Bourguignon, Thomas, Qian Tang, Simon Y. W. Ho, Frantisek Juna, Zongqing Wang, Daej A. Arab, Stephen L. Cameron, et al. 2018. “Transoceanic Dispersal and Plate Tectonics Shaped Global Cockroach Distributions: Evidence from Mitochondrial Phylogenomics.” Molecular Biology and Evolution 35 (4): 970–83. |
| Primates | Cassandra Cruz | 148 | 455 | T/T | 90 | Jonika, Michelle M., Kayla T. Wilhoit, Maximos Chin, Abhimanyu Arekere, and Heath Blackmon. 2024. “Drift Drives the Evolution of Chromosome Number II: The Impact of Range Size on Genome Evolution in Carnivora.” The Journal of Heredity 115 (5): 524–31.<br><br>Craig, Jack M., S. Blair Hedges, and Sudhir Kumar. 2024. “Completing a Molecular Timetree of Primates.” Frontiers in Bioinformatics 4 (December): 1495417. |
| Chrysomelidae | Emily Clark | 869 | 5870 | F/T | 182 | Blackmon, Heath, and Jeffery P. Demuth. 2015. “Coleoptera Karyotype Database.” The Coleopterists’ Bulletin 69 (1): 174–75. |
| Coccinellidae | Emily Clark | 159 | 222 | F/T | 24 | Blackmon, Heath, and Jeffery P. Demuth. 2015. “Coleoptera Karyotype Database.” The Coleopterists’ Bulletin 69 (1): 174–75.<br><br>Che, Liheng, Peng Zhang, Shaohong Deng, Hermes E. Escalona, Xingmin Wang, Yun Li, Hong Pang, et al. 2021. “New Insights into the Phylogeny and Evolution of Lady Beetles (Coleoptera: Coccinellidae) by Extensive Sampling of Genes and Species.” Molecular Phylogenetics and Evolution 156 (107045): 107045. |
| Lepidoptera | Eunice Lee | 3370 | 2258 | T/T | 322 | Chen, Xi, Zuoqi Wang, Chaowei Zhang, Jingheng Hu, Yueqi |

| | | | | | | |
|---|---|---|---|---|---|---|
| | | | | | | Lu, Hang Zhou, Yang Mei, et al. 2023. “Unraveling the Complex Evolutionary History of Lepidopteran Chromosomes through Ancestral Chromosome Reconstruction and Novel Chromosome Nomenclature.” *BMC Biology* 21 (1): 265.<br><br>Kawahara, Akito Y., Caroline Storer, Ana Paula S. Carvalho, David M. Plotkin, Fabien L. Condamine, Mariana P. Braga, Emily A. Ellis, et al. 2023. “A Global Phylogeny of Butterflies Reveals Their Evolutionary History, Ancestral Hosts and Biogeographic Origins.” Nature Ecology & Evolution 7 (6): 903–13. |
| Orchidaceae | Eunice Lee | 8122 | 41474 | T/F | 1904 | Rice, Anna, Lior Glick, Shiran Abadi, Moshe Einhorn, Naama M. Kopelman, Ayelet Salman-Minkov, Jonathan Mayzel, Ofer Chay, and Itay Mayrose. 2015. “The Chromosome Counts Database (CCDB) - a Community Resource of Plant Chromosome Numbers.” The New Phytologist 206 (1): 19–26.<br><br>Smith, Stephen A., and Joseph W. Brown. 2018. “Constructing a Broadly Inclusive Seed Plant Phylogeny.” American Journal of Botany 105 (3): 302–14. |
| Hymenoptera | Gideon Schlab | 1591 | 602 | T/T | 345 | Cardoso DC, Santos HG, Cristiano MP. The Ant Chromosome database – ACdb: an online resource for ant (Hymenoptera: Formicidae) chromosome researchers. Myrmecol News. 2018;27:87-91.<br><br>Tree of Sex Consortium, 2014. Tree of sex: a database of sexual systems. Scientific Data, 1, p.140015.<br><br>Blaimer, Bonnie B., Bernardo F. Santos, Astrid Cruaud, Michael W. Gates, Robert R. Kula, István Mikó, Jean-Yves Rasplus, et al. 2023. “Key Innovations and the Diversification of Hymenoptera.” Nature Communications 14 (1): 1212. |
| Muridae | Gideon Schalb | 103 | 913 | T/F | 57 | Jonika, Michelle M., Kayla T. Wilhoit, Maximos Chin, Abhimanyu Arekere, and Heath Blackmon. 2024. “Drift Drives the Evolution of Chromosome Number II: The Impact of Range Size on Genome Evolution in Carnivora.” The Journal of Heredity 115 (5): 524–31. |

| | | | | | | |
|---|---|---|---|---|---|---|
| | | | | | | Steppan, Scott J., and John J. Schenk. 2017. “Muroid Rodent Phylogenetics: 900-Species Tree Reveals Increasing Diversification Rates.” PloS One 12 (8): e0183070. |
| Odonata | Hannah Abraham | 703 | 669 | T/T | 84 | Kuznetsova, Valentina G., and Natalia V. Golub. 2020. “A Checklist of Chromosome Numbers and a Review of Karyotype Variation in Odonata of the World.” Comparative Cytogenetics 14 (4): 501–40.<br><br>Willink, Beatriz, Jessica L. Ware, and Erik I. Svensson. 2024. “Tropical Origin, Global Diversification, and Dispersal in the Pond Damselflies (Coenagrionoidea) Revealed by a New Molecular Phylogeny.” Systematic Biology 73 (2): 290–307. |
| Testudines | Hannah Abraham | 141 | 593 | F/F | 122 | Román-Palacios, Cristian, Cesar A. Medina, Shing H. Zhan, and Michael S. Barker. 2021. “Animal Chromosome Counts Reveal a Similar Range of Chromosome Numbers but with Less Polyploidy in Animals Compared to Flowering Plants.” Journal of Evolutionary Biology 34 (8): 1333–39.<br><br>Thomson, Robert C., Phillip Q. Spinks, and H. Bradley Shaffer. 2021. “A Global Phylogeny of Turtles Reveals a Burst of Climate-Associated Diversification on Continental Margins.” Proceedings of the National Academy of Sciences of the United States of America 118 (7): e2012215118. |
| Curculionidae | Heath Blackmon | 617 | 1492 | T/T | 33 | Blackmon, Heath, and Jeffery P. Demuth. 2015. “Coleoptera Karyotype Database.” The Coleopterists’ Bulletin 69 (1): 174–75.<br><br>Haran, J., X. Li, R. Allio, S. Shin, L. Benoit, R. G. Oberprieler, B. D. Farrell, et al. 2023. “Phylogenomics Illuminates the Phylogeny of Flower Weevils (Curculioninae) and Reveals Ten Independent Origins of Brood-Site Pollination Mutualism in True Weevils.” Proceedings. Biological Sciences 290 (2008): 20230889. [Data set]. Zenodo. https://doi.org/10.5281/zenodo.7849369 |
| Fungi | Heath Blackmon | 151 | 5418 | T/T | 40 | Zolan, M. E. 1995. “Chromosome-Length Polymorphism in Fungi.” Microbiological Reviews 59 (4): 686–98. |

| | | | | | | |
|---|---|---|---|---|---|---|
| | | | | | | Kumar, Sudhir, Michael Suleski, Jack M. Craig, Adrienne E. Kasprowicz, Maxwell Sanderford, Michael Li, Glen Stecher, and S. Blair Hedges. 2022. “TimeTree 5: An Expanded Resource for Species Divergence Times.” Molecular Biology and Evolution 39 (8). https://doi.org/10.1093/molbev/msac174. |
| Cyprinidae | Kaya Harper | 1111 | 1368 | T/T | 398 | Nagpure, Naresh Sahebrao, Ajey Kumar Pathak, Rameshwar Pati, Iliyas Rashid, Jyoti Sharma, Shri Prakash Singh, Mahender Singh, et al. 2016. “Fish Karyome Version 2.1: A Chromosome Database of Fishes and Other Aquatic Organisms.” Database: The Journal of Biological Databases and Curation 2016 (March): baw012.<br><br>Arai, Ryoichi. 2011. Fish Karyotypes: A Check List. PDF. 2011th ed. Tokyo, Japan: Springer.<br><br>Chang, Jonathan, Daniel L. Rabosky, Stephen A. Smith, and Michael E. Alfaro. 2019. “An r Package and Online Resource for Macroevolutionary Studies Using the Ray‐finned Fish Tree of Life.” Methods in Ecology and Evolution 10 (7): 1118–24. |
| Gekkonidae | Kaya Harper | 73 | 1331 | T/T | 59 | Román-Palacios, Cristian, Cesar A. Medina, Shing H. Zhan, and Michael S. Barker. 2021. “Animal Chromosome Counts Reveal a Similar Range of Chromosome Numbers but with Less Polyploidy in Animals Compared to Flowering Plants.” Journal of Evolutionary Biology 34 (8): 1333–39.<br><br>Tonini, João Filipe Riva, Karen H. Beard, Rodrigo Barbosa Ferreira, Walter Jetz, and R. Alexander Pyron. 2016. “Fully-Sampled Phylogenies of Squamates Reveal Evolutionary Patterns in Threat Status.” Biological Conservation 204 (December): 23–31. |
| Scincoidea | Kenzie Laird | 154 | 1519 | T/T | 134 | Román-Palacios, Cristian, Cesar A. Medina, Shing H. Zhan, and Michael S. Barker. 2021. “Animal Chromosome Counts Reveal a Similar Range of Chromosome Numbers but with Less Polyploidy in Animals Compared to Flowering Plants.” Journal of Evolutionary Biology 34 (8): 1333–39. |

| | | | | | | |
|---|---|---|---|---|---|---|
| | | | | | | Tonini, João Filipe Riva, Karen H. Beard, Rodrigo Barbosa Ferreira, Walter Jetz, and R. Alexander Pyron. 2016. “Fully-Sampled Phylogenies of Squamates Reveal Evolutionary Patterns in Threat Status.” Biological Conservation 204 (December): 23–31. |
| Serpentes | Kenzie Laird | 256 | 1877 | T/T | 213 | Román-Palacios, Cristian, Cesar A. Medina, Shing H. Zhan, and Michael S. Barker. 2021. “Animal Chromosome Counts Reveal a Similar Range of Chromosome Numbers but with Less Polyploidy in Animals Compared to Flowering Plants.” Journal of Evolutionary Biology 34 (8): 1333–39.<br><br>Kumar, Sudhir, Michael Suleski, Jack M. Craig, Adrienne E. Kasprowicz, Maxwell Sanderford, Michael Li, Glen Stecher, and S. Blair Hedges. 2022. “TimeTree 5: An Expanded Resource for Species Divergence Times.” Molecular Biology and Evolution 39 (8). https://doi.org/10.1093/molbev/msac174. |
| Characidae | Kiedon Bryant | 663 | 1019 | T/T | 238 | Nagpure, Naresh Sahebrao, Ajey Kumar Pathak, Rameshwar Pati, Iliyas Rashid, Jyoti Sharma, Shri Prakash Singh, Mahender Singh, et al. 2016. “Fish Karyome Version 2.1: A Chromosome Database of Fishes and Other Aquatic Organisms.” Database: The Journal of Biological Databases and Curation 2016 (March): baw012.<br><br>Arai, Ryoichi. 2011. Fish Karyotypes: A Check List. 2011. Tokyo, Japan: Springer.<br><br>Melo, Bruno F., Rafaela P. Ota, Ricardo C. Benine, Fernando R. Carvalho, Flavio C. T. Lima, George M. T. Mattox, Camila S. Souza, et al. 2024. “Phylogenomics of Characidae, a Hyper-Diverse Neotropical Freshwater Fish Lineage, with a Phylogenetic Classification Including Four Families (Teleostei: Characiformes).” Zoological Journal of the Linnean Society 202 (1). https://doi.org/10.1093/zoolinnean/zlae101. |
| Cichlidae | Kiedon Bryant | 281 | 750 | T/T | 88 | Nagpure, Naresh Sahebrao, Ajey Kumar Pathak, Rameshwar Pati, Iliyas Rashid, Jyoti Sharma, Shri Prakash Singh, Mahender Singh, et al. 2016. “Fish Karyome Version 2.1: A Chromosome Database of Fishes and Other Aquatic |

| | | | | | | |
|---|---|---|---|---|---|---|
| | | | | | | Organisms.” Database: The Journal of Biological Databases and Curation 2016 (March): baw012.<br><br>Arai, Ryoichi. 2011. Fish Karyotypes: A Check List. 2011. Tokyo, Japan: Springer.<br><br>Rabosky, Daniel L., Jonathan Chang, Pascal O. Title, Peter F. Cowman, Lauren Sallan, Matt Friedman, Kristin Kaschner, et al. 2018. “An Inverse Latitudinal Gradient in Speciation Rate for Marine Fishes.” Nature 559 (7714): 392–95. |
| Nothobranchiidae | Mallory Murphy | 242 | 11638 | T/T | 79 | Nagpure, Naresh Sahebrao, Ajey Kumar Pathak, Rameshwar Pati, Iliyas Rashid, Jyoti Sharma, Shri Prakash Singh, Mahender Singh, et al. 2016. “Fish Karyome Version 2.1: A Chromosome Database of Fishes and Other Aquatic Organisms.” Database: The Journal of Biological Databases and Curation 2016 (March): baw012.<br><br>Arai, Ryoichi. 2011. Fish Karyotypes: A Check List. 2011. Tokyo, Japan: Springer.<br><br>Merwe, P. De Wet van der, Fenton P. D. Cotterill, Martha Kandziora, Brian R. Watters, Béla Nagy, Tyrone Genade, Tyrel J. Flügel, David S. Svendsen, and Dirk U. Bellstedt. 2021. “Genomic Fingerprints of Palaeogeographic History: The Tempo and Mode of Rift Tectonics across Tropical Africa Has Shaped the Diversification of the Killifish Genus Nothobranchius (Teleostei: Cyprinodontiformes).” Molecular Phylogenetics and Evolution 158 (106988): 106988. |
| Orthoptera | Mallory Murphy | 319 | 232 | T/T | 36 | Sylvester, Terrence and Heath Blackmon. 2019. Idiosycratic patterns of chromosome evolution are the rule not the exception.<br><br>Song, Hojun, Ricardo Mariño-Pérez, Derek A. Woller, and Maria Marta Cigliano. 2018. “Evolution, Diversification, and Biogeography of Grasshoppers (Orthoptera: Acrididae).” Insect Systematics and Diversity 2 (4): 3. |
| Brassicaceae | Megan | 2822 | 1557 | T/F | 1557 | Rice, Anna, Lior Glick, Shiran Abadi, Moshe Einhorn, Naama |

| | | | | | | |
|---|---|---|---|---|---|---|
| | Copeland | | | | | M. Kopelman, Ayelet Salman-Minkov, Jonathan Mayzel, Ofer Chay, and Itay Mayrose. 2015. “The Chromosome Counts Database (CCDB) - a Community Resource of Plant Chromosome Numbers.” The New Phytologist 206 (1): 19–26.<br><br>Smith, Stephen A., and Joseph W. Brown. 2018. “Constructing a Broadly Inclusive Seed Plant Phylogeny.” American Journal of Botany 105 (3): 302–14. |
| Solanaceae | Megan Copeland | 1458 | 930 | T/F | 930 | Rice, Anna, Lior Glick, Shiran Abadi, Moshe Einhorn, Naama M. Kopelman, Ayelet Salman-Minkov, Jonathan Mayzel, Ofer Chay, and Itay Mayrose. 2015. “The Chromosome Counts Database (CCDB) - a Community Resource of Plant Chromosome Numbers.” The New Phytologist 206 (1): 19–26.<br><br>Smith, Stephen A., and Joseph W. Brown. 2018. “Constructing a Broadly Inclusive Seed Plant Phylogeny.” American Journal of Botany 105 (3): 302–14. |
| Iguania | Meghann McConnell | 382 | 1416 | F/T | 353 | Román-Palacios, Cristian, Cesar A. Medina, Shing H. Zhan, and Michael S. Barker. 2021. “Animal Chromosome Counts Reveal a Similar Range of Chromosome Numbers but with Less Polyploidy in Animals Compared to Flowering Plants.” Journal of Evolutionary Biology 34 (8): 1333–39.<br><br>Kumar, Sudhir, Michael Suleski, Jack M. Craig, Adrienne E. Kasprowicz, Maxwell Sanderford, Michael Li, Glen Stecher, and S. Blair Hedges. 2022. “TimeTree 5: An Expanded Resource for Species Divergence Times.” Molecular Biology and Evolution 39 (8). https://doi.org/10.1093/molbev/msac174. |
| Tenebrionidae | Meghann McConnell | 239 | 318 | T/T | 40 | Blackmon, Heath, and Jeffery P. Demuth. 2015. “Coleoptera Karyotype Database.” The Coleopterists’ Bulletin 69 (1): 174–75.<br><br>Li, Yun, Craig Moritz, Ian G. Brennan, Andreas Zwick, James Nicholls, Alicia Grealy, and Adam Slipinski. 2024. “Evolution across the Adaptive Landscape in a Hyperdiverse Beetle Radiation.” Current Biology 34 (16): 3685-3697.e6. |
| Cetacea | Nayeli | 40 | 90 | T/T | 34 | Blackmon, Heath, Joshua Justison, Itay Mayrose, and Emma |

| | | | | | | |
|---|---|---|---|---|---|---|
| | Perez | | | | | E. Goldberg. 2019. “Meiotic Drive Shapes Rates of Karyotype Evolution in Mammals.” Evolution; International Journal of Organic Evolution 73 (3): 511–23.<br><br>Kumar, Sudhir, Michael Suleski, Jack M. Craig, Adrienne E. Kasprowicz, Maxwell Sanderford, Michael Li, Glen Stecher, and S. Blair Hedges. 2022. “TimeTree 5: An Expanded Resource for Species Divergence Times.” Molecular Biology and Evolution 39 (8). https://doi.org/10.1093/molbev/msac174. |
| Siluriformes | Nayeli Perez | 165 | 4205 | F/T | 131 | Nagpure, Naresh Sahebrao, Ajey Kumar Pathak, Rameshwar Pati, Iliyas Rashid, Jyoti Sharma, Shri Prakash Singh, Mahender Singh, et al. 2016. “Fish Karyome Version 2.1: A Chromosome Database of Fishes and Other Aquatic Organisms.” Database: The Journal of Biological Databases and Curation 2016 (March): baw012.<br><br>Arai, Ryoichi. 2011. Fish Karyotypes: A Check List. 2011. Tokyo, Japan: Springer.<br><br>Pinna, Mário de, Luiz Peixoto, Victor Tagliacollo, and Marcelo Britto. 2025. “Phylogenetic Relationships and Evolution of the Major Groups of Siluriformes.” In Catfishes, a Highly Diversified Group, 97–127. Boca Raton: CRC Press. |
| Carabidae | Rachel Koehl | 777 | 104 | T/T | 95 | Blackmon, Heath, and Jeffery P. Demuth. 2015. “Coleoptera Karyotype Database.” The Coleopterists’ Bulletin 69 (1): 174–75.<br><br>Kavanaugh, David H., David R. Maddison, W. Brian Simison, Sean D. Schoville, Joachim Schmidt, Arnaud Faille, Wendy Moore, et al. 2021. “Phylogeny of the Supertribe Nebriitae (Coleoptera, Carabidae) Based on Analyses of DNA Sequence Data.” ZooKeys 1044 (June): 41–152. |
| Liliaceae | Riya Girish | 648 | 79340 | T/F | 430 | Rice, Anna, Lior Glick, Shiran Abadi, Moshe Einhorn, Naama M. Kopelman, Ayelet Salman-Minkov, Jonathan Mayzel, Ofer Chay, and Itay Mayrose. 2015. “The Chromosome Counts Database (CCDB) - a Community Resource of Plant Chromosome Numbers.” The New Phytologist 206 (1): 19–26. |

| | | | | | | |
|---|---|---|---|---|---|---|
| | | | | | | Smith, Stephen A., and Joseph W. Brown. 2018. “Constructing a Broadly Inclusive Seed Plant Phylogeny.” American Journal of Botany 105 (3): 302–14. |
| Rubiaceae | Riya Girish | 1219 | 794 | T/F | 794 | Rice, Anna, Lior Glick, Shiran Abadi, Moshe Einhorn, Naama M. Kopelman, Ayelet Salman-Minkov, Jonathan Mayzel, Ofer Chay, and Itay Mayrose. 2015. “The Chromosome Counts Database (CCDB) - a Community Resource of Plant Chromosome Numbers.” The New Phytologist 206 (1): 19–26.<br><br>Smith, Stephen A., and Joseph W. Brown. 2018. “Constructing a Broadly Inclusive Seed Plant Phylogeny.” American Journal of Botany 105 (3): 302–14. |
| Anabantiformes | Megan Copeland | 61 | 196 | T/T | 41 | Nagpure, Naresh Sahebrao, Ajey Kumar Pathak, Rameshwar Pati, Iliyas Rashid, Jyoti Sharma, Shri Prakash Singh, Mahender Singh, et al. 2016. “Fish Karyome Version 2.1: A Chromosome Database of Fishes and Other Aquatic Organisms.” Database: The Journal of Biological Databases and Curation 2016 (March): baw012.<br><br>Arai, Ryoichi. 2011. Fish Karyotypes: A Check List. 2011. Tokyo, Japan: Springer.<br><br>S. Kumar, M. Suleski, J.E. Craig, A.E. Kasprowicz, M. Sanderford, M. Li, G. Stecher, and S.B. Hedges, 2022. TimeTree 5: An Expanded Resource for Species Divergence Times. Molecular Biology and Evolution, DOI: 10.1093/molbev/msac174. |
| Gobiidae | Megan Copeland | 232 | 827 | F/T | 65 | Nagpure, Naresh Sahebrao, Ajey Kumar Pathak, Rameshwar Pati, Iliyas Rashid, Jyoti Sharma, Shri Prakash Singh, Mahender Singh, et al. 2016. “Fish Karyome Version 2.1: A Chromosome Database of Fishes and Other Aquatic Organisms.” Database: The Journal of Biological Databases and Curation 2016 (March): baw012.<br><br>Arai, Ryoichi. 2011. Fish Karyotypes: A Check List. 2011. Tokyo, Japan: Springer. |

| | | | | | | |
|---|---|---|---|---|---|---|
| | | | | | | McCraney, W. Tyler, Christine E. Thacker, and Michael E. Alfaro. 2020. “Supermatrix Phylogeny Resolves Goby Lineages and Reveals Unstable Root of Gobiaria.” Molecular Phylogenetics and Evolution 151 (106862): 106862. |
| Bryophyta | Sarah Schmalz | 4666 | 533 | F/T | 239 | Rice, Anna, Lior Glick, Shiran Abadi, Moshe Einhorn, Naama M. Kopelman, Ayelet Salman-Minkov, Jonathan Mayzel, Ofer Chay, and Itay Mayrose. 2015. “The Chromosome Counts Database (CCDB) - a Community Resource of Plant Chromosome Numbers.” *The New Phytologist* 206 (1): 19–26.<br><br>Kumar, Sudhir, Michael Suleski, Jack M. Craig, Adrienne E. Kasprowicz, Maxwell Sanderford, Michael Li, Glen Stecher, and S. Blair Hedges. 2022. “TimeTree 5: An Expanded Resource for Species Divergence Times.” Molecular Biology and Evolution 39 (8). https://doi.org/10.1093/molbev/msac174. |
| Passifloraceae | Sarah Schmalz | 159 | 134 | T/F | 134 | Rice, Anna, Lior Glick, Shiran Abadi, Moshe Einhorn, Naama M. Kopelman, Ayelet Salman-Minkov, Jonathan Mayzel, Ofer Chay, and Itay Mayrose. 2015. “The Chromosome Counts Database (CCDB) - a Community Resource of Plant Chromosome Numbers.” The New Phytologist 206 (1): 19–26.<br><br>Smith, Stephen A., and Joseph W. Brown. 2018. “Constructing a Broadly Inclusive Seed Plant Phylogeny.” American Journal of Botany 105 (3): 302–14. |
| Hemiptera | Sean Chien | 1744 | 1967 | T/T | 46 | Tree of Sex Consortium, 2014. Tree of sex: a database of sexual systems. Scientific Data, 1, p.140015.<br><br>Kumar, Sudhir, Michael Suleski, Jack M. Craig, Adrienne E. Kasprowicz, Maxwell Sanderford, Michael Li, Glen Stecher, and S. Blair Hedges. 2022. “TimeTree 5: An Expanded Resource for Species Divergence Times.” Molecular Biology and Evolution 39 (8). https://doi.org/10.1093/molbev/msac174. |
| Phasmatodea | Sean Chien | 109 | 148 | T/T | 11 | Sylvester, Terrence, Carl E. Hjelmen, Shawn J. Hanrahan, Paul A. Lenhart, J. Spencer Johnston, and Heath Blackmon. 2020. “Lineage-Specific Patterns of Chromosome Evolution Are the Rule Not the Exception in Polyneoptera Insects.” |

| | | | | | | |
|---|---|---|---|---|---|---|
| | | | | | | Proceedings. Biological Sciences 287 (1935): 20201388.<br><br>Kumar, Sudhir, Michael Suleski, Jack M. Craig, Adrienne E. Kasprowicz, Maxwell Sanderford, Michael Li, Glen Stecher, and S. Blair Hedges. 2022. “TimeTree 5: An Expanded Resource for Species Divergence Times.” Molecular Biology and Evolution 39 (8). https://doi.org/10.1093/molbev/msac174. |
| Dytiscidae | Shelbie Cast | 85 | 973 | T/T | 29 | Blackmon, Heath, and Jeffery P. Demuth. 2015. “Coleoptera Karyotype Database.” The Coleopterists’ Bulletin 69 (1): 174–75.<br><br>Ribera, Ignacio, Alfried P. Vogler, and Michael Balke. 2008. “Phylogeny and Diversification of Diving Beetles (Coleoptera: Dytiscidae).” Cladistics: The International Journal of the Willi Hennig Society 24 (4): 563–90. |
| Hydrophilidae | Shelbie Cast | 81 | 168 | T/T | 21 | Blackmon, Heath, and Jeffery P. Demuth. 2015. “Coleoptera Karyotype Database.” The Coleopterists’ Bulletin 69 (1): 174–75.<br><br>Kumar, Sudhir, Michael Suleski, Jack M. Craig, Adrienne E. Kasprowicz, Maxwell Sanderford, Michael Li, Glen Stecher, and S. Blair Hedges. 2022. “TimeTree 5: An Expanded Resource for Species Divergence Times.” Molecular Biology and Evolution 39 (8). https://doi.org/10.1093/molbev/msac174. |
| Anura | Steven Arackal | 1831 | 5326 | T/T | 1207 | Perkins, Riddhi D., Julio Rincones Gamboa, Michelle M. Jonika, Johnathan Lo, Amy Shum, Richard H. Adams, and Heath Blackmon. 2019. “A Database of Amphibian Karyotypes.” Chromosome Research: An International Journal on the Molecular, Supramolecular and Evolutionary Aspects of Chromosome Biology 27 (4): 313–19.<br><br>Portik, Daniel M., Jeffrey W. Streicher, and John J. Wiens. 2023. “Frog Phylogeny: A Time-Calibrated, Species-Level Tree Based on Hundreds of Loci and 5,242 Species.” Molecular Phylogenetics and Evolution 188 (107907): 107907. |
| Cricetidae | Steven Arackal | 208 | 913 | T/T | 103 | Jonika, Michelle M., Kayla T. Wilhoit, Maximos Chin, Abhimanyu Arekere, and Heath Blackmon. 2024. “Drift Drives |

| | | | | | | |
|---|---|---|---|---|---|---|
| | | | | | | the Evolution of Chromosome Number II: The Impact of Range Size on Genome Evolution in Carnivora.” The Journal of Heredity 115 (5): 524–31.<br><br>Bangs, Max R., Alexandre R. Percequillo, Víctor Pacheco, and Scott J. Steppan. 2024. “Phylogenomics of the Sigmodontine Rodents: Cloud Forests and Pliocene Extinction Explain Timing and Spread of the Radiation of South American Mice and Rats.” bioRxiv.https://doi.org/10.1101/2024.12.25.630327. |
| Chiroptera | Tanvi Koneru | 204 | 815 | T/T | 154 | Jonika, Michelle M., Kayla T. Wilhoit, Maximos Chin, Abhimanyu Arekere, and Heath Blackmon. 2024. “Drift Drives the Evolution of Chromosome Number II: The Impact of Range Size on Genome Evolution in Carnivora.” The Journal of Heredity 115 (5): 524–31.<br><br>Agnarsson, Ingi, Carlos M. Zambrana-Torrelio, Nadia Paola Flores-Saldana, and Laura J. May-Collado. 2011. “A Time-Calibrated Species-Level Phylogeny of Bats (Chiroptera, Mammalia).” PLoS Currents 3 (February): RRN1212. |
| Drosophilidae | Tanvi Koneru | 1246 | 685 | T/T | 352 | Morelli, Magnolia W., Heath Blackmon, and Carl E. Hjelmen. 2022. “Diptera and Drosophila Karyotype Databases: A Useful Dataset to Guide Evolutionary and Genomic Studies.” Frontiers in Ecology and Evolution 10 (March). https://doi.org/10.3389/fevo.2022.832378.<br><br>Kumar, Sudhir, Michael Suleski, Jack M. Craig, Adrienne E. Kasprowicz, Maxwell Sanderford, Michael Li, Glen Stecher, and S. Blair Hedges. 2022. “TimeTree 5: An Expanded Resource for Species Divergence Times.” Molecular Biology and Evolution 39 (8). https://doi.org/10.1093/molbev/msac174. |
| Asteraceae | Virginia Lopez | 6497 | 3336 | T/F | 3336 | Rice, Anna, Lior Glick, Shiran Abadi, Moshe Einhorn, Naama M. Kopelman, Ayelet Salman-Minkov, Jonathan Mayzel, Ofer Chay, and Itay Mayrose. 2015. “The Chromosome Counts Database (CCDB) - a Community Resource of Plant Chromosome Numbers.” The New Phytologist 206 (1): 19–26.<br><br>Smith, Stephen A., and Joseph W. Brown. 2018. “Constructing |

| | | | | | | |
|---|---|---|---|---|---|---|
| | | | | | | a Broadly Inclusive Seed Plant Phylogeny." American Journal of Botany 105 (3): 302–14. |
| Fabaceae | Virginia Lopez | 2006 | 1692 | T/F | 1692 | Rice, Anna, Lior Glick, Shiran Abadi, Moshe Einhorn, Naama M. Kopelman, Ayelet Salman-Minkov, Jonathan Mayzel, Ofer Chay, and Itay Mayrose. 2015. "The Chromosome Counts Database (CCDB) - a Community Resource of Plant Chromosome Numbers." The New Phytologist 206 (1): 19–26.<br><br>Smith, Stephen A., and Joseph W. Brown. 2018. "Constructing a Broadly Inclusive Seed Plant Phylogeny." American Journal of Botany 105 (3): 302–14. |
| Araneae | Various authors | 1298 | 969 | F/T | 87 | Araujo, D.; Schneider, M.C.; Paula-Neto, E.; Cella, D.M. 2025. The spider cytogenetic database. Available in www.arthropodacytogenetics.bio.br/spiderdatabase<br><br>Wheeler, Ward C., Jonathan A. Coddington, Louise M. Crowley, Dimitar Dimitrov, Pablo A. Goloboff, Charles E. Griswold, Gustavo Hormiga, et al. 2017. "The Spider Tree of Life: Phylogeny of Araneae Based on Target-Gene Analyses from an Extensive Taxon Sampling." Cladistics: The International Journal of the Willi Hennig Society 33 (6): 574–616. |
| Galliformes | Various authors | 53 | 52 | T/T | 52 | Alfieri, James M., Reina Hingoranee, Giridhar N. Athrey, and Heath Blackmon. 2024. "Domestication Is Associated with Increased Interspecific Hybrid Compatibility in Landfowl (Order: Galliformes)." The Journal of Heredity 115 (1): 1–10.<br><br>Alfieri, James M., Kevin Bolwerk, Zhaobo Hu, and Heath Blackmon. 2024. "From Micro to Macro: Avian Chromosome Evolution Is Dominated by Natural Selection." bioRxivorg. https://doi.org/10.1101/2024.11.29.626112. |
| Gymnospermae | Various authors | 2026 | 31749 | T/T | 375 | Rice, Anna, Lior Glick, Shiran Abadi, Moshe Einhorn, Naama M. Kopelman, Ayelet Salman-Minkov, Jonathan Mayzel, Ofer Chay, and Itay Mayrose. 2015. "The Chromosome Counts Database (CCDB) - a Community Resource of Plant Chromosome Numbers." The New Phytologist 206 (1): 19–26. |

| | | | | | | |
|---|---|---|---|---|---|---|
| | | | | | | Smith, Stephen A., and Joseph W. Brown. 2018. “Constructing a Broadly Inclusive Seed Plant Phylogeny.” American Journal of Botany 105 (3): 302–14. |
| Magnoliaceae | Various authors | 139 | 304 | T/F | 77 | Rice, Anna, Lior Glick, Shiran Abadi, Moshe Einhorn, Naama M. Kopelman, Ayelet Salman-Minkov, Jonathan Mayzel, Ofer Chay, and Itay Mayrose. 2015. “The Chromosome Counts Database (CCDB) - a Community Resource of Plant Chromosome Numbers.” The New Phytologist 206 (1): 19–26.<br><br>Smith, Stephen A., and Joseph W. Brown. 2018. “Constructing a Broadly Inclusive Seed Plant Phylogeny.” American Journal of Botany 105 (3): 302–14. |
| Marsupialia | Various authors | 40 | 40 | T/T | 40 | Blackmon, Heath, Joshua Justison, Itay Mayrose, and Emma E. Goldberg. 2019. “Meiotic Drive Shapes Rates of Karyotype Evolution in Mammals.” Evolution; International Journal of Organic Evolution 73 (3): 511–23.<br><br>Kumar, Sudhir, Michael Suleski, Jack M. Craig, Adrienne E. Kasprowicz, Maxwell Sanderford, Michael Li, Glen Stecher, and S. Blair Hedges. 2022. “TimeTree 5: An Expanded Resource for Species Divergence Times.” Molecular Biology and Evolution 39 (8). https://doi.org/10.1093/molbev/msac174. |
| Scarabidae | Various authors | 478 | 211 | T/T | 174 | Chien and Blackmon. Chromosomal Rearrangements: Tempo and Mode of Karyotype Evolution in Scarabaeoidea (submitted Journal of Evolutionary Biology)<br><br>Blackmon, Heath, and Jeffery P. Demuth. 2015. “Coleoptera Karyotype Database.” The Coleopterists’ Bulletin 69 (1): 174–75. |
| Scorpiones | Various authors | 320 | 190 | T/T | 36 | Schneider, M.C.; Mattos, V.F.; Cella, D.M. 2025. The scorpion cytogenetic database. Available in www.arthropodacytogenetics.bio.br/scorpiondatabase<br><br>Kumar, Sudhir, Michael Suleski, Jack M. Craig, Adrienne E. Kasprowicz, Maxwell Sanderford, Michael Li, Glen Stecher, and S. Blair Hedges. 2022. “TimeTree 5: An Expanded Resource for Species Divergence Times.” Molecular Biology |

| | | | | | | |
|---|---|---|---|---|---|---|
| | | | | | | and Evolution 39 (8). https://doi.org/10.1093/molbev/msac174. |

#### MCMC Convergence

Effective sample size (ESS) was used to evaluate chain mixing and sampling efficiency and to identify parameters with elevated autocorrelation. ESS values were calculated from post–burn-in MCMC samples using the coda package and provide a measure of chain mixing and sampling efficiency.

**Table S2**. **Effective sample size (ESS) diagnostics for Bayesian chromosome evolution models.** ESS values for post–burn-in MCMC samples are shown for all estimated parameters across clades.

| Clade | Asc1 | Desc1 | Demi | Poly |
|---|---|---|---|---|
| Accipitriformes | 1900 | 1585.62873 | 715.365591 | 323.312868 |
| Anabantiformes | 743.685372 | 525.640379 | 239.370123 | 681.546713 |
| Anura | 1790.46339 | 1633.86593 | 2156.11014 | 1900 |
| Araneae | 338.365048 | 323.05475 | 406.929341 | 286.242533 |
| Asteraceae | 773.985301 | 665.578676 | 569.935714 | 900 |
| Blattodea | 687.327276 | 239.710209 | 504.719774 | 446.788115 |
| Brassicaceae | 912.8312 | 900 | 762.773383 | 900 |
| Bryophyta | 502.114067 | 634.704999 | 464.146828 | 525.690945 |
| Carabidae | 567.728046 | 707.841545 | 755.326188 | 385.502121 |
| Carnivora | 475.45622 | 429.240176 | 662.655167 | 740.827092 |
| Caudata | 637.158759 | 470.812042 | 772.147261 | 386.902505 |
| Cetacea | 805.40829 | 428.226823 | 705.32682 | 576.32958 |
| Characidae | 3270.36055 | 267.02256 | 408.523608 | 1882.06601 |
| Chiroptera | 978.733497 | 869.834793 | 1653.25198 | 381.878853 |
| Chondrichthyes | 3389.18956 | 1503.40485 | 4310.8604 | 7001.57287 |
| Chrysomelidae | 616.690879 | 700.551859 | 732.395617 | 693.488516 |
| Cichlidae | 688.698503 | 900 | 794.380704 | 339.558534 |
| Coccinellidae | 401.272235 | 679.978151 | 472.223617 | 366.957934 |
| Cricetidae | 428.948629 | 510.257899 | 440.963211 | 634.671187 |
| Curculionidae | 376.674399 | 424.381751 | 644.468148 | 606.612127 |
| Cyprinidae | 900 | 900 | 900 | 721.512501 |

| Clade | Asc1 | Desc1 | Demi | Poly |
|---|---|---|---|---|
| Drosophilidae | 1249.33992 | 334.962655 | 482.864922 | 1247.45221 |
| Dytiscidae | 316.210839 | 432.389402 | 521.691789 | 1169.50905 |
| Fabaceae | 561.250066 | 900 | 668.57821 | 900 |
| Fungi | 522.309589 | 366.135279 | 576.900608 | 809.711298 |
| Galliformes | 1508.48606 | 1309.56294 | 1457.27006 | 504.720693 |
| Gekkonidae | 881.511751 | 2313.31018 | 5264.53465 | 32978.8397 |
| Gobiidae | 615.082729 | 657.90489 | 900 | 645.51057 |
| Gymnospermae | 645.453702 | 609.06176 | 689.352665 | 900 |
| Hemiptera | 232.282027 | 327.717257 | 267.089647 | 276.058867 |
| Hydrophilidae | 255.792315 | 723.270734 | 466.922954 | 254.429088 |
| Hymenoptera | 651.508211 | 635.467161 | 601.575492 | 777.326915 |
| Iguania | 777.207384 | 769.810088 | 690.21555 | 482.516329 |
| Lepidoptera | 542.283242 | 741.877087 | 346.130277 | 603.33512 |
| Liliaceae | 682.447371 | 789.460929 | 900 | 703.549837 |
| Magnoliaceae | 255.580941 | 256.181174 | 604.26875 | 846.899972 |
| Marsupialia | 587.967721 | 488.140531 | 623.90269 | 356.291296 |
| Muridae | 405.258962 | 540.152883 | 630.09907 | 622.528861 |
| Nothobranchiidae | 488.520628 | 426.822477 | 534.388996 | 222.605053 |
| Odonata | 46954.3002 | 70789.3307 | 50718.4256 | NA |
| Odonataforced_mcmc | 95891.6274 | 143821.342 | 95898.0813 | 66056.3898 |
| Orchidaceae | 442.286077 | 862.795435 | 1100.5405 | 900 |
| Orthoptera | 19613.3099 | 1829.59532 | 18000.6573 | 4789.36625 |
| Passeriformes | 726.201814 | 1055.57383 | 900.050666 | 508.044955 |
| Passifloraceae | 530.376118 | 430.325762 | 797.158303 | 901.707821 |
| Phasmatodea | 295.857286 | 454.776352 | 351.168774 | 517.88846 |
| Primates | 251.697218 | 244.416772 | 448.578993 | 326.147719 |
| Pteridophyta | 817.261577 | 900 | 886.736213 | 900 |
| Rubiaceae | 900 | 900 | 900 | 900 |

| Clade | Asc1 | Desc1 | Demi | Poly |
|---|---|---|---|---|
| Scarabidae | 3139.00635 | 1075.22303 | 3089.15348 | 6447.8689 |
| Scincoidea | 759.471745 | 541.362255 | 452.469754 | 296.328038 |
| Scorpiones | 256.67631 | 366.883647 | 656.668986 | 624.719726 |
| Serpentes | 1598.69586 | 1656.19612 | 723.994118 | 728.362645 |
| Siluriformes | 567.375043 | 566.713108 | 610.353034 | 562.831187 |
| Solanaceae | 495.226476 | 753.951939 | 900 | 900 |
| Tenebrionidae | 787.009159 | 574.75645 | 372.61789 | 568.20908 |
| Testudines | 620.140692 | 900 | 900 | 461.883148 |

### Mentor vs Mentee results

To assess the reproducibility of rate estimates generated, we compared dysploidy rate inferences produced independently by student analysts to those generated by mentor-led validation runs. For each clade, the mentor analysis re-executed the full analytical workflow starting from the curated chromosome count and phylogenetic inputs in parallel, using identical model specifications and inference settings. Comparisons between student and mentor estimates were evaluated by direct comparison of posterior medians and overlap of 95% highest posterior density (HPD) intervals for the total dysploidy rate (the sum of ascending and descending dysploidy). We found that the dysploidy estimates were highly concordant between student and mentor analyses (Fig. S1). The interaction of the HPD intervals intersect on or close to the one-to-one expectation line. These results demonstrate that chromosome evolution rate estimates are robust to analyst identity and independent implementation of the pipeline, supporting the reliability of the distributed analytical framework used in this study.

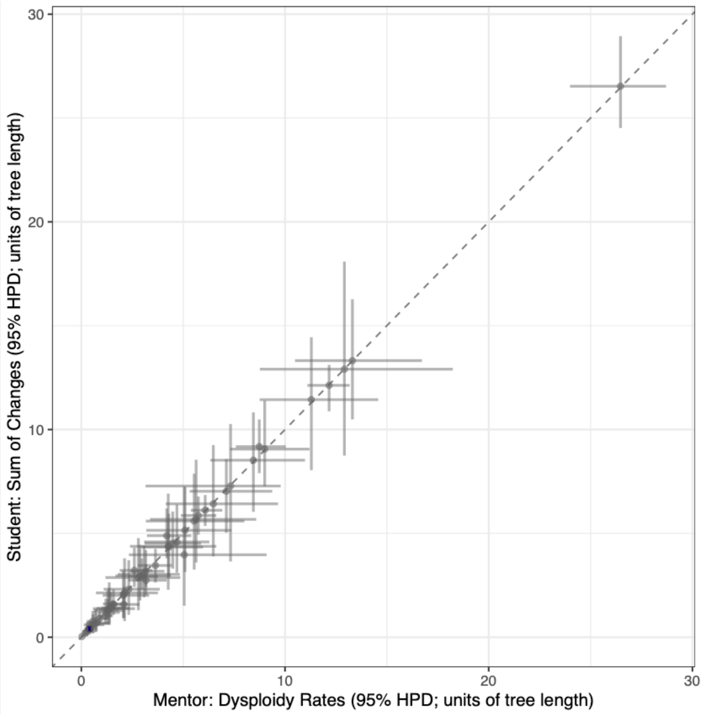


**Figure S1**. **Comparison of total dysploidy rate estimates (sum of ascending and descending dysploidy) obtained independently by student analysts and by mentor-led validation analyses for each clade.** Points represent posterior median estimates, with horizontal and vertical error bars indicating 95% highest posterior density (HPD) intervals expressed in units of tree length. The dashed line denotes the one-to-one expectation. This plot illustrates the results of final analyses. Earlier iterations were used to identify two clades where the student and mentor had different inferences. In both cases these were caused by errors in the analysis code and once fixed the above 1:1 ratios were observed.

### Prior Influence

#### Exponential vs Uniform

One potential concern is that by using an exponential prior that favors low rates we are artificially biasing our results towards low rates. This concern is reflected by the fact that for each dataset the points in Figure S2 fall above the diagonal. However, we note that many of the most extreme rate estimates are unchanged and the relative ranking of taxa is relatively stable to different priors. For instance the three fastest evolving clades change order but the same three clades are still ranked most highly. For low rate clades, the results are even more stable with the ordering of the six slowest evolving clades remaining nearly unchanged.

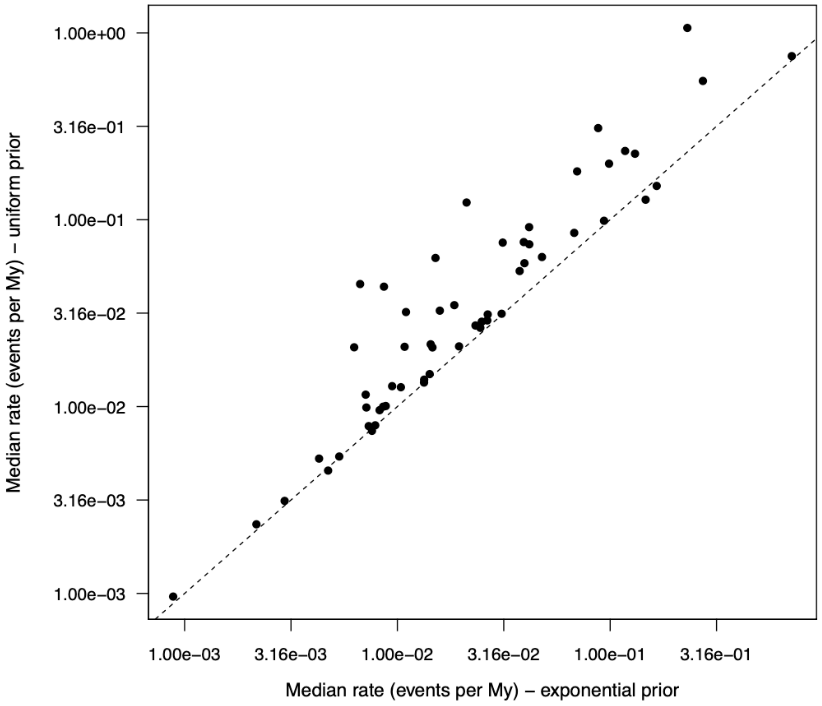


**Figure S2.** Robustness of chromosome evolution rate estimates to prior specification. Comparison of median posterior dysploidy rate estimates inferred under exponential and uniform priors across clades. Each point represents a single clade, plotted on logarithmic axes in units of events per million years. The dashed line indicates the one-to-one expectation.

#### Interactions between tree size and prior

To ensure that the global heterogeneity in chromosomal evolutionary rates reported in the main text reflects biological reality rather than statistical artifacts driven by clade size, we performed two validation analyses. In Bayesian frameworks, estimates from small datasets can be susceptible to the "pull of the prior," where the posterior distribution collapses toward the prior mean rather than reflecting the data1. Given that our study compares clades ranging from tens to thousands of species, we tested whether differences in sampling density introduce systematic bias using the R packages chromePlus (*31*), diversitree (*32*), TreeSim (*37*), and ape (*35*).

*Simulation-Based Validation*

We first conducted a simulation experiment to test the robustness of our model to extreme data loss. We simulated 100 phylogenies with a tree depth of 1 and with 500 taxa each using a birth-death process (lambda=3, mu=1) implemented in TreeSim (*37*). 100 Chromosome number datasets were evolved along these trees under a Markov model with equal rates (fusion = 1 and fission = 1) using simChrom in chromePlus (*31*). For each replicate, we estimated the rates of chromosome evolution on the full tree (n=500) using a likelihood function with the function make.mkn from the package diversitree (*3*2) constrained to exclude polyploidy and demi-polyploidy (constrainMkn from the package chromePlus) (*31*). We then randomly pruned 90% of the tips, retaining only 50 taxa, and re-estimated the rates using the same exponential prior lambda=2 used in the main text.

The results demonstrate that even after a 90% reduction in data, the posterior parameter estimates for the pruned datasets consistently recover the "true" rate derived from the full datasets (Fig. S3). We quantified the concordance between the two datasets by calculating the overlap of the 95% Highest Posterior Density (HPD) intervals. In the 100 simulation replicates, we observed only 5 instances (of the 200 possible) where the HPD intervals of the pruned and full datasets failed to overlap (a 2.5% false positive rate). This confirms that even with moderate taxonomic sampling (n=50), the biological signal retains sufficient strength to overcome the prior and recover the correct evolutionary tempo.

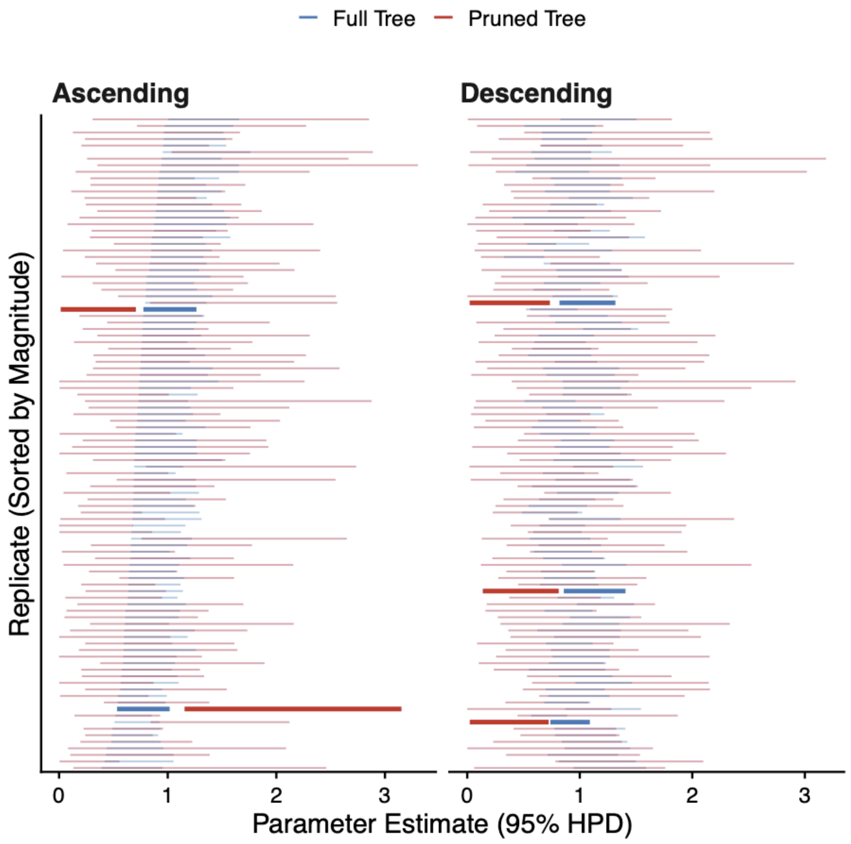


**Figure S3. Robustness of chromosome evolution rate estimates to extreme data loss.** Posterior parameter estimates (95% Highest Posterior Density, HPD) for chromosome evolution rates simulated on 100 phylogenetic trees. Each replicate compares estimates from the full dataset (500 taxa, blue) against the same dataset randomly pruned by 90% (50 taxa, red). Panels show rate estimates for ascending (fission) and descending (fusion) dysploidy. Replicates are sorted along the y-axis by the magnitude of the pruned tree estimate. Despite the 90% reduction in data, the 95% HPD intervals between full and pruned datasets overlap in 97.5% of comparisons, indicating that the biological signal for chromosomal evolution remains robust even with moderate taxonomic sampling (n=50)

*Empirical Validation (Scarabaeidae)*

To validate this pattern with empirical data, which contains inherent rate heterogeneity absent in simulations, we analyzed the family Scarabaeidae (scarab beetles). We utilized the phylogeny containing 174 species used in the primary analysis. We estimated the evolutionary rates for the full clade and compared them to 100 replicates in which the tree was randomly subsampled to 50% of its original richness (n=87).

Consistent with the simulation results, rate estimates from the subsampled empirical trees largely overlapped with the estimates from the full dataset (Fig. S4). We observed only 2 instances of non-overlapping HPD intervals across 100 replicates each of which has two opportunities (ascending and descending rate comparisons providing a study wide false positive rate of 1%). Furthermore, we detected no systematic shift in the posterior distribution toward the prior mean in the subsampled replicates. These combined analyses indicate that the order-of-magnitude differences in evolutionary rates observed across the Tree of Life cannot be attributed to the pull of the prior in smaller clades. While smaller clades yield broader posterior distributions reflecting greater uncertainty, the signal of rate magnitude is unbiased and robust.

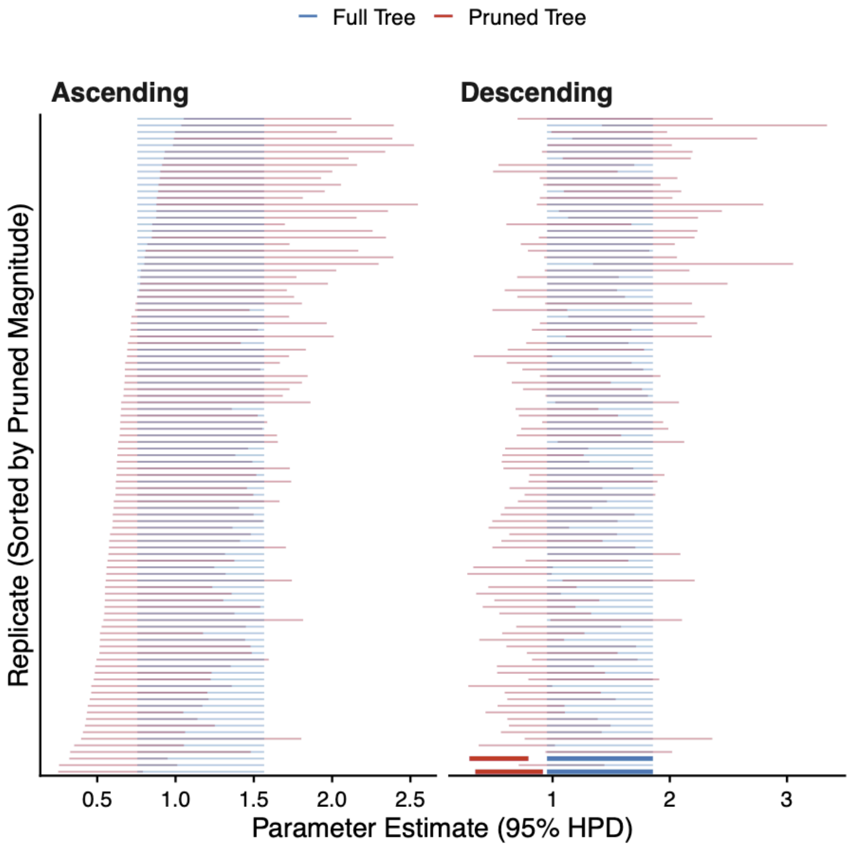


**Figure S4. Empirical validation of rate estimate robustness using *Scarabaeidae*.** Posterior parameter estimates (95% Highest Posterior Density, HPD) for ascending (fission) and descending (fusion) dysploidy rates derived from the full *Scarabaeidae* phylogeny (n=174, blue) compared against 100 replicates where the tree was randomly subsampled to 50% richness (n=87, red). The blue intervals represent the estimate from the single full dataset, repeated across rows for comparison, while red intervals represent independent subsampled replicates sorted by magnitude. Bold lines indicate the rare instances (2 out of 200 comparisons) where the HPD intervals of the subsampled and full datasets failed to overlap. The analysis demonstrates that reducing taxonomic sampling by half does not introduce a systematic bias toward the prior or alter the inferred order of magnitude for evolutionary rates.

#### Model Comparison

To evaluate support for alternative chromosome transition models, we compared the fit of models that differed in the inclusion of polyploidy and demiploidy using Akaike Information Criterion (AIC). For many plant clades, the model excluding both polyploidy and demiploidy could not be evaluated. The observed chromosome state space and transition structure did not support stable likelihood optimization after removing these parameters, resulting in non-convergent or undefined AIC values. Across the 56 clades examined (including Magnoliaceae), the fully parameterized model often produced the lowest AIC. We comparing the full model and a reduced model excluding polyploidy, we identified 28 clades showing strong support for polyploidy inclusion using a cutoff ΔAIC of 5. Of the 28 clades, 12 are plant groups with the other 16 being animal groups as the fungi clade did not show support (Table S3).

**Table S3**. **AIC-based comparisons of alternate chromosome evolution models across clades.** Lower AIC scores indicate better model support.

| Clade | Full | No Poly | No Demi | Drop Both |
|---|---|---|---|---|
| Accipitriformes | 240.990268 | 238.990218 | 238.990218 | 236.990218 |
| Anabantiformes | 182.42615 | 201.177782 | 180.42615 | 232.723532 |
| Anura | 2453.34223 | 2844.19021 | 2866.03997 | 5139.54417 |
| Araneae | 500.834802 | 498.443818 | 498.849983 | 496.103843 |
| Asteraceae | 16489.3789 | 20548.4273 | 18288.3553 | NA |
| Blattodea | 193.957026 | 197.665547 | 200.901203 | 206.69342 |
| Brassicaceae | 8143.27079 | 10213.3995 | 9474.93095 | 20022.2674 |
| Bryophyta | 1073.85571 | 1226.1518 | 1081.7789 | 1468.82726 |
| Carabidae | 363.199 | 361.422477 | 366.617168 | 369.419821 |
| Carnivora | 423.823875 | 452.035553 | 443.563109 | 556.685308 |
| Caudata | 373.336128 | 367.172745 | 472.267409 | 495.541538 |
| Cetacea | 24.9436224 | 22.9437909 | 22.967304 | 20.9436193 |
| Characidae | 652.455549 | 675.871001 | 669.148055 | 877.748981 |
| Chiroptera | 708.967884 | 706.967883 | 736.963722 | 732.422857 |
| Chondrichthyes | 480.198342 | 480.99967 | 500.367256 | 524.470286 |
| Chrysomelidae | 809.056143 | 817.844538 | 894.239153 | 957.531234 |
| Cichlidae | 226.910462 | 225.482361 | 257.680082 | 255.696872 |
| Coccinellidae | 83.5465258 | 81.5465258 | 85.7218207 | 85.7131134 |
| Cricetidae | 584.046617 | 591.630745 | 582.046617 | 615.963621 |

| Clade | Full | No Poly | No Demi | Drop Both |
|---|---|---|---|---|
| Curculionidae | 144.453868 | 151.150772 | 160.046493 | 166.572036 |
| Cyprinidae | 573.335634 | 780.805176 | 1305.81152 | 2205.97327 |
| Drosophilidae | 527.804576 | 525.804575 | 541.666558 | 539.666555 |
| Dytiscidae | 108.136464 | 114.779942 | 106.136464 | 132.285523 |
| Fabaceae | 5946.34932 | 7296.75743 | 6728.17395 | NA |
| Fungi | 215.321085 | 213.321083 | 213.321083 | 211.321083 |
| Galliformes | 278.191954 | 283.222739 | 288.291109 | 321.864897 |
| Gekkonidae | 226.826832 | 224.865492 | 236.345481 | 235.634577 |
| Gobiidae | 268.659255 | 269.606788 | 349.362989 | 684.430955 |
| Gymnospermae | 716.506394 | 896.307575 | 800.658902 | 1524.54242 |
| Hemiptera | 233.553768 | 231.553768 | 230.089642 | 238.397589 |
| Hydrophilidae | 56.4471426 | 54.4471423 | 59.5839253 | 61.5437628 |
| Hymenoptera | 1858.07331 | 1882.07226 | 1943.69196 | 2120.76204 |
| Iguania | 1307.78742 | 1315.43547 | 1435.07779 | 1440.48393 |
| Lepidoptera | 2182.47547 | 2212.33589 | 2203.55863 | 2358.89122 |
| Liliaceae | 1215.23181 | 1688.83684 | 1673.67289 | 2711.22588 |
| Magnoliaceae | 119.256637 | 307.970282 | 328.883394 | 391.631678 |
| Marsupialia | 106.445827 | 104.445827 | 120.274832 | 118.461327 |
| Muridae | 332.510354 | 332.762515 | 336.798327 | 334.798327 |
| Nothobranchiidae | 409.808079 | 408.911754 | 419.688164 | 425.93902 |
| Odonata | 78.9015608 | 76.9015536 | 101.700324 | 99.7003211 |
| Orchidaceae | 9421.09151 | 10361.7831 | 9878.1715 | NA |
| Orthoptera | 98.1345171 | 111.182437 | 104.163969 | 113.795355 |
| Passeriformes | 1519.60342 | 1519.60342 | 1538.78895 | 1538.78895 |
| Passifloraceae | 417.979747 | 529.769402 | 552.741578 | 739.634492 |
| Phasmatodea | 56.6784778 | 59.3813704 | 54.678476 | 65.4547842 |
| Primates | 444.105651 | 443.915105 | 450.092584 | 467.970856 |
| Pteridophyta | 9065.88588 | 17648.2763 | 15141.7301 | NA |
| Rubiaceae | 2720.64348 | 4402.09394 | 3882.00406 | NA |

| Clade | Full | No Poly | No Demi | Drop Both |
|---|---|---|---|---|
| Scarabidae | 462.304568 | 461.637963 | 501.31899 | 511.507442 |
| Scincoidea | 248.635381 | 246.641189 | 259.465538 | 257.465538 |
| Scorpiones | 233.274905 | 231.274905 | 254.032317 | 251.923215 |
| Serpentes | 530.564562 | 528.56456 | 528.56456 | 526.56456 |
| Siluriformes | 1385.58061 | 1422.60378 | 1783.20541 | 2063.93921 |
| Solanaceae | 1906.86888 | 3152.56387 | 2463.06238 | 5832.48306 |
| Tenebrionidae | 91.754721 | 94.9659345 | 89.7547209 | 109.437734 |
| Testudines | 316.727845 | 326.090109 | 377.085435 | 442.785353 |

#### Clade Audits

For each clade included in our analysis we performed a three step validation and adequacy procedure.

*Continuous Trait Mapping*

We first plotted ancestral state estimates of chromosome number, treated as a continuous character evolving via Brownian motion. Though this is a simplified model it is a fast and convenient approach to visualize the distribution of approximate chromosome numbers along the branches of a phylogeny. These plots were constructed using the ContMap function from phytools (*38*). This visualization allowed us check that the range and distribution of chromosome number was reasonable, and that the distribution of branch lengths in the tree did not appear concerning (e.g. a large number of taxa extending from a single node as is the case for trees with large number of taxonomically placed species).

*Prior Only Analysis*

Next we generated prior only runs for all MCMCs. Briefly using this approach we are able to compare the rate inference for our data to one where the prior used in our analysis is held constant but no data is available. By comparing rate estimates based only on the prior to those based on both we are able to determine whether the data has sufficient signal to define the rate estimate and allows us to exclude any taxa where the data were insufficient to shift the posterior away from the prior distribution.

*Posterior Predictive Simulations*

Finally, to evaluate whether the fitted chromosome evolution models adequately reproduce key features of the observed data, we conducted posterior predictive simulations for each of the 56 monophyletic clades analyzed in this study. For each clade, posterior predictive datasets were generated using the same empirical chromosome counts and phylogenies used for Bayesian rate inference.

Posterior parameter samples were obtained from the post–burn-in portion of the Bayesian MCMC chains, with the first 25% of iterations discarded as burn-in. To generate posterior predictive simulations, 1,000 parameter vectors were randomly drawn from the post–burn-in posterior distribution. For clades represented by multiple phylogenetic trees, a single tree was randomly selected for each simulation replicate, thereby propagating phylogenetic uncertainty into the posterior predictive distributions.

Chromosome number evolution was simulated on the empirical phylogeny using the function simChrom in the chromePlus package (*31*). Simulations incorporated ascending dysploidy, descending dysploidy, demiploidy, and polyploidy transitions when these parameters were included in the fitted model; when specific transition parameters were not estimated for a clade, the corresponding rates were set to zero. The root chromosome number for each simulation was set to the rounded mean of the empirical haploid counts for that clade. To prevent unbounded exploration of chromosome state space while retaining biological realism, simulated chromosome numbers were constrained to lie between one (i.e., a biological constraint) and five chromosomes beyond the maximum haploid count observed empirically for each clade.

For both the empirical data and each simulated dataset, we calculated two summary statistics across the tips of the phylogeny: the variance of haploid chromosome numbers and Shannon entropy of the chromosome number distribution. Variance captures the overall dispersion of chromosome counts, while entropy reflects the diversity and evenness of chromosome states and is sensitive to skewed frequency distributions. These complementary statistics were chosen to assess whether fitted models reproduce both the magnitude and structure of chromosome number variation.

Empirical values of variance and entropy were compared to their corresponding posterior predictive distributions. For each statistic, the empirical value was assigned a percentile based on the proportion of simulated values less than or equal to the empirical estimate. Clades were flagged as exhibiting potential inadequacy for a given statistic when the empirical value fell within the 5% tails of the posterior predictive distribution. Posterior predictive density plots for variance and entropy were generated for each clade, with empirical values shown as vertical reference lines (Fig. S5-S60), and a summary table recording adequacy flags across clades was compiled (Table S4).

*Interpretation of posterior predictive results*

Across most clades, empirical variance and entropy values fell within the central region of the posterior predictive distributions, indicating that the fitted models capture key features of observed chromosome number variation. However, departures from posterior predictive expectations were not uniform across statistics. In several clades, empirical entropy values fell toward the lower tail of the posterior predictive distribution despite variance being well reproduced, reflecting highly uneven chromosome state frequencies rather than a failure to capture overall dispersion. Conversely, a smaller number of clades showed mismatches in variance while entropy remained well predicted, indicating localized differences in the magnitude of chromosome number spread rather than in state diversity. Importantly, these deviations were clade-specific and statistic-specific rather than systematic across major taxonomic groups. No kingdom exhibited consistent failure across both summary statistics, and clades showing posterior predictive departures were interspersed among those showing strong adequacy.

Bryophyta | Higher Taxonomy: Bryophyta | Chromosomes sampled: 4,666 | Phylogenetic tips: 533 | Overlap with phylogeny: 239 species | Root age: 488 Ma | Taxonomic tips: FALSE | Unresolved: N/A

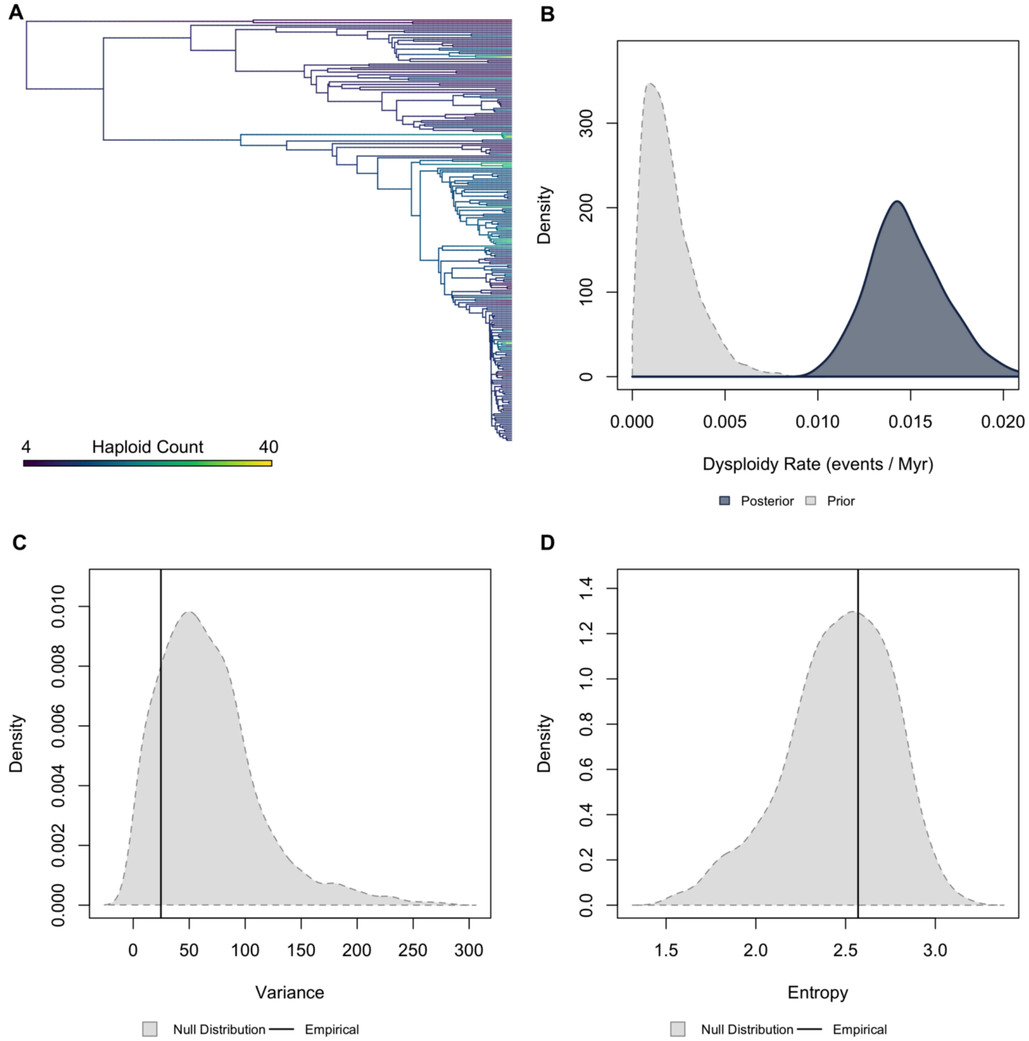


**Figure S5. Validation plots for Bryophyta.** (A) Continuous character map of haploid chromosome number across the phylogeny. (B) Prior versus posterior density distributions for dysploidy rate. (C) Posterior predictive simulation adequacy test for variance. (D) Posterior predictive simulation adequacy test for Shannon’s entropy.

Pteridophyta | Higher Taxonomy: Pteridophyta | Chromosomes sampled: 12,075 | Phylogenetic tips: 5,868 | Overlap with phylogeny: 1,465 species | Root age: 429 Ma | Taxonomic tips: FALSE | Unresolved: N/A

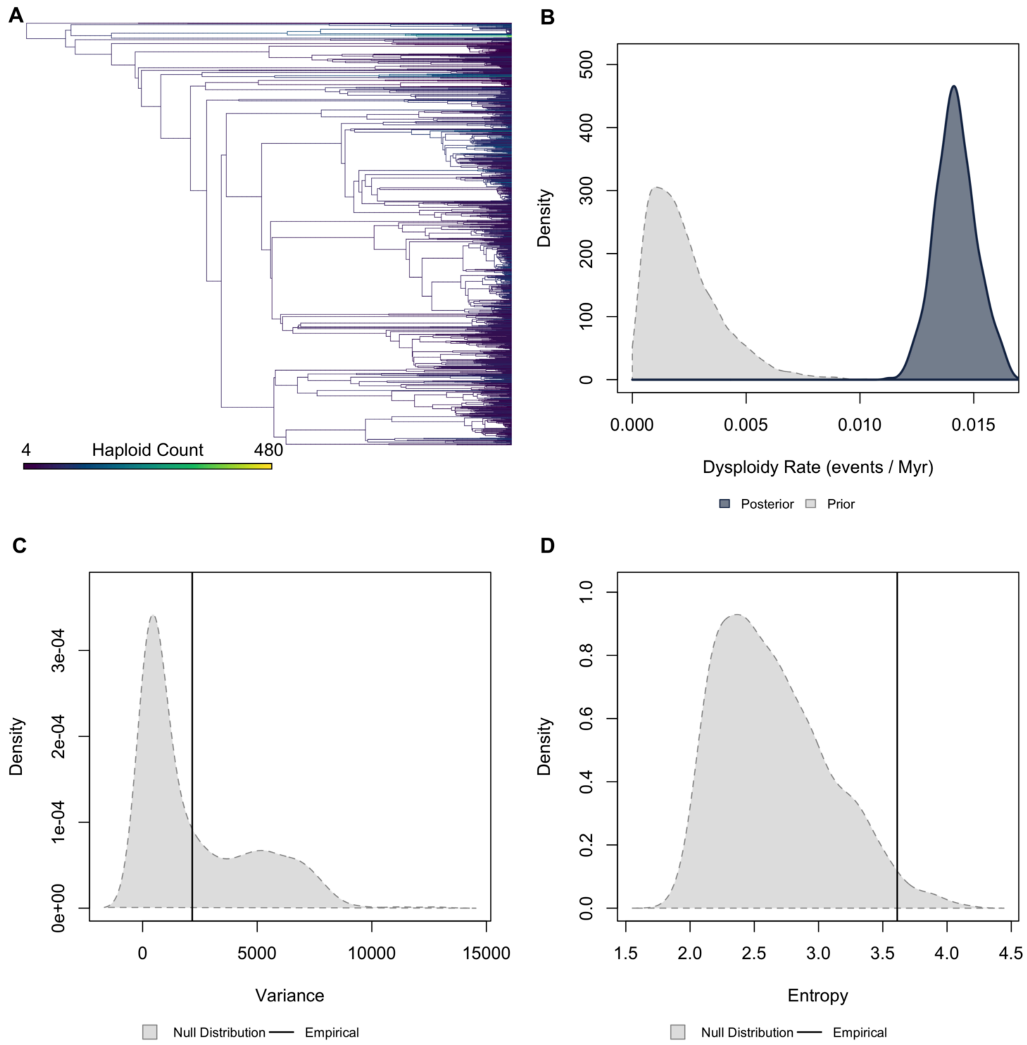


**Figure S6. *Validation plots for Pteridophyta.*** *(A) Continuous character map of haploid chromosome number across the phylogeny. (B) Prior versus posterior density distributions for dysploidy rate. (C) Posterior predictive simulation adequacy test for variance. (D) Posterior predictive simulation adequacy test for Shannon's entropy.*

Gymnospermae | Higher Taxonomy: Gymnospermae | Chromosomes sampled: 2,026 | Phylogenetic tips: 31,749 | Overlap with phylogeny: 375 species | Root age: 304 Ma | Taxonomic tips: FALSE | Unresolved: N/A

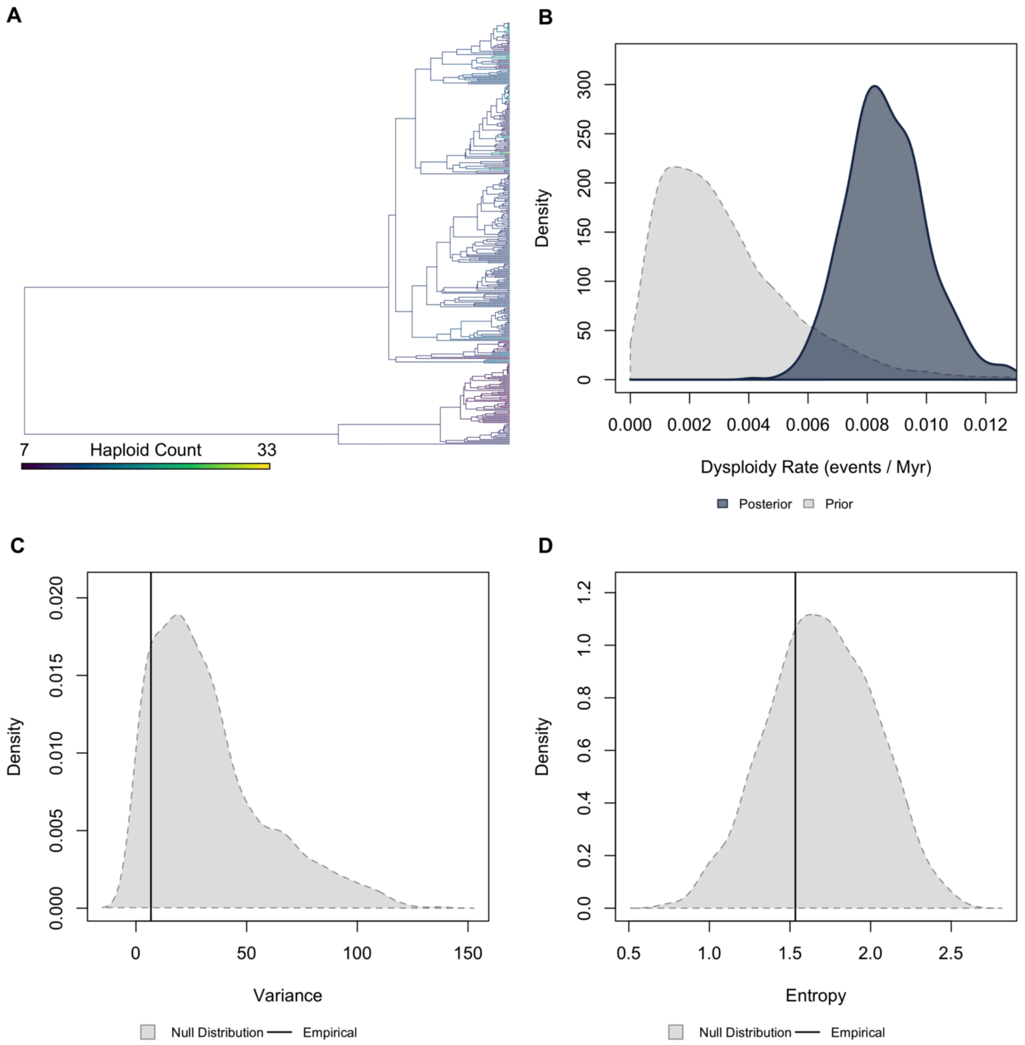


**Figure S7.** ***Validation plots for Gymnospermae.*** *(A) Continuous character map of haploid chromosome number across the phylogeny. (B) Prior versus posterior density distributions for dysploidy rate. (C) Posterior predictive simulation adequacy test for variance. (D) Posterior predictive simulation adequacy test for Shannon's entropy.*

Asteraceae | Higher Taxonomy: Angiospermae | Chromosomes sampled: 6,497 | Phylogenetic tips: 3,336 | Overlap with phylogeny: 3,336 species | Root age: 160 Ma | Taxonomic tips: TRUE | Unresolved: 21.52%

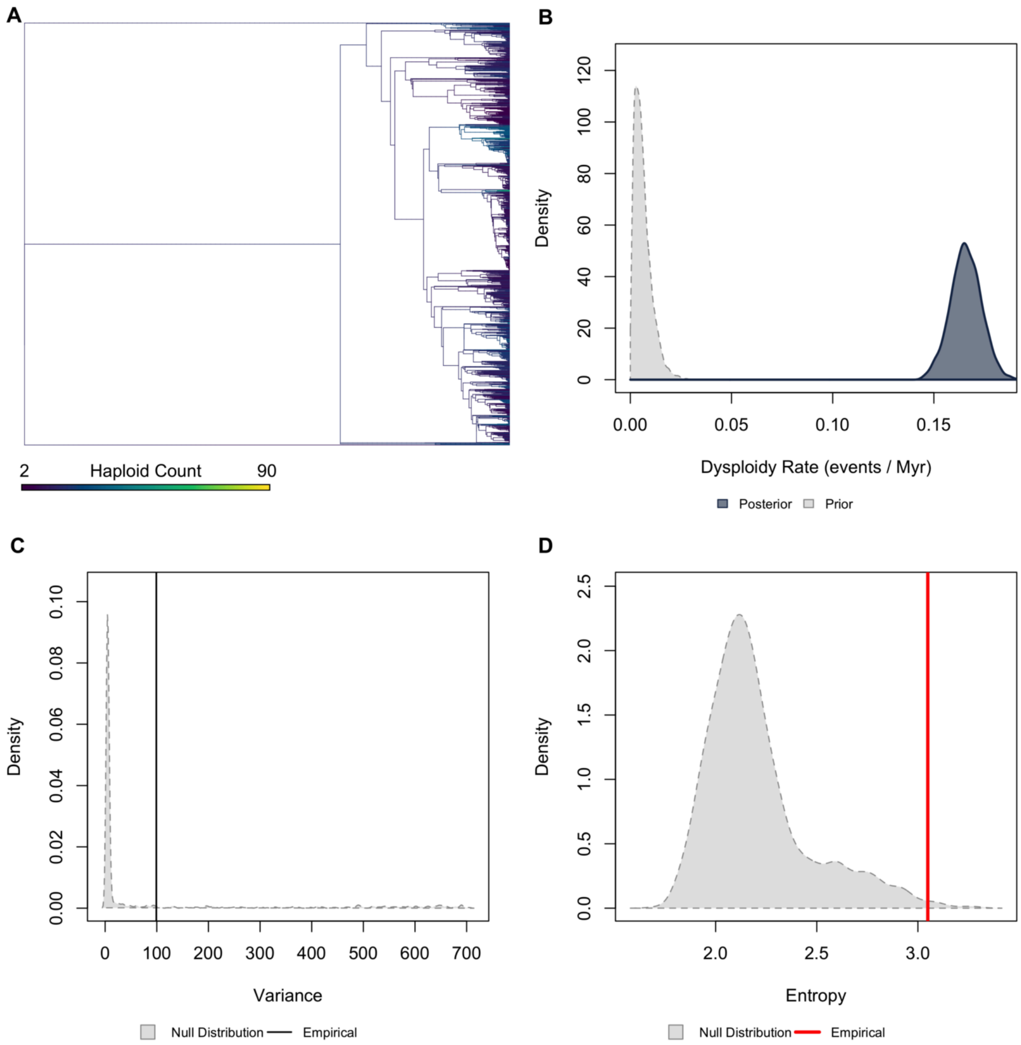


**Figure S8.** ***Validation plots for Asteraceae.*** *(A) Continuous character map of haploid chromosome number across the phylogeny. (B) Prior versus posterior density distributions for dysploidy rate. (C) Posterior predictive simulation adequacy test for variance. (D) Posterior predictive simulation adequacy test for Shannon's entropy.*

Brassicaceae | Higher Taxonomy: Angiospermae | Chromosomes sampled: 2,822 | Phylogenetic tips: 1,557 | Overlap with phylogeny: 1,557 species | Root age: 118 Ma | Taxonomic tips: TRUE | Unresolved: 45.02%

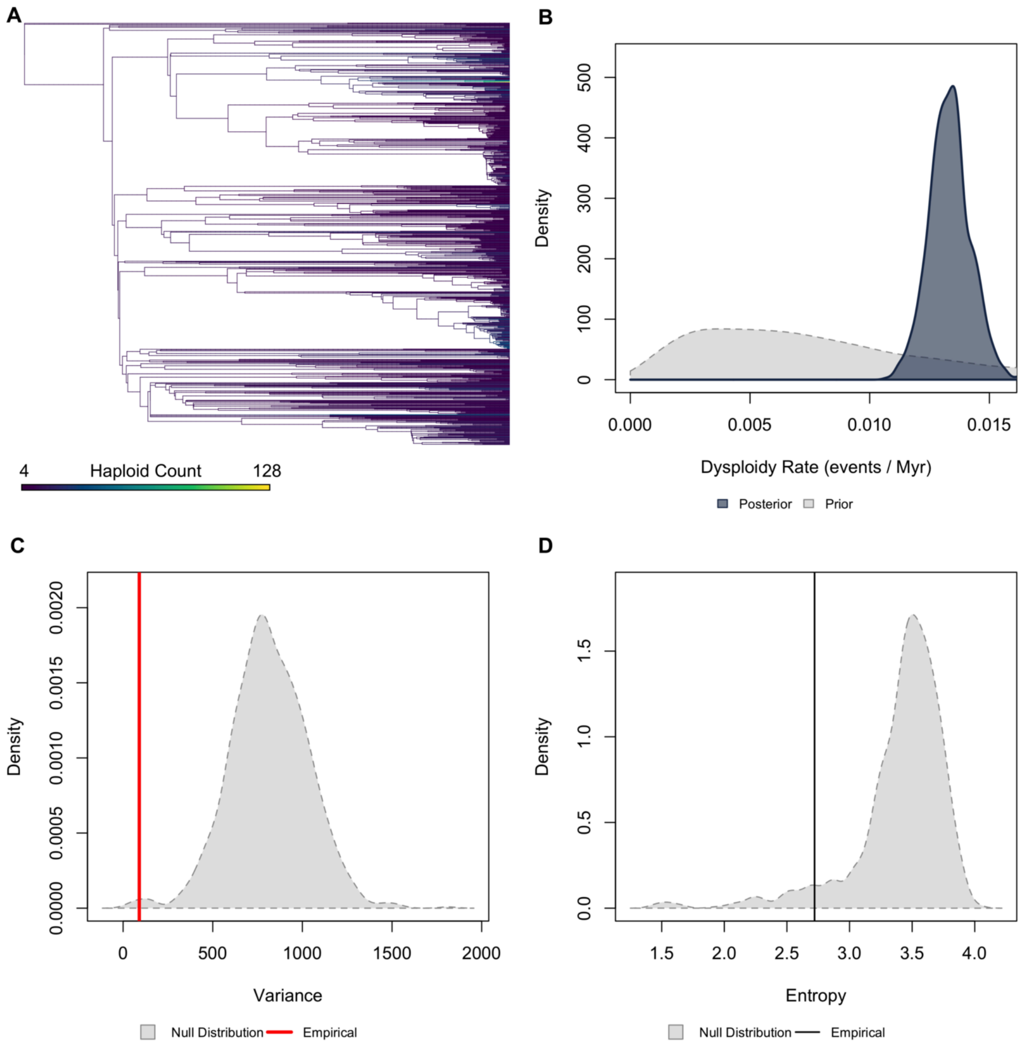


**Figure S9. *Validation plots for Brassicaceae.*** *(A) Continuous character map of haploid chromosome number across the phylogeny. (B) Prior versus posterior density distributions for dysploidy rate. (C) Posterior predictive simulation adequacy test for variance. (D) Posterior predictive simulation adequacy test for Shannon's entropy.*

Fabaceae | Higher Taxonomy: Angiospermae | Chromosomes sampled: 2,006 | Phylogenetic tips: 1,692 | Overlap with phylogeny: 1,692 species | Root age: 68 Ma | Taxonomic tips: TRUE | Unresolved: 55.56%

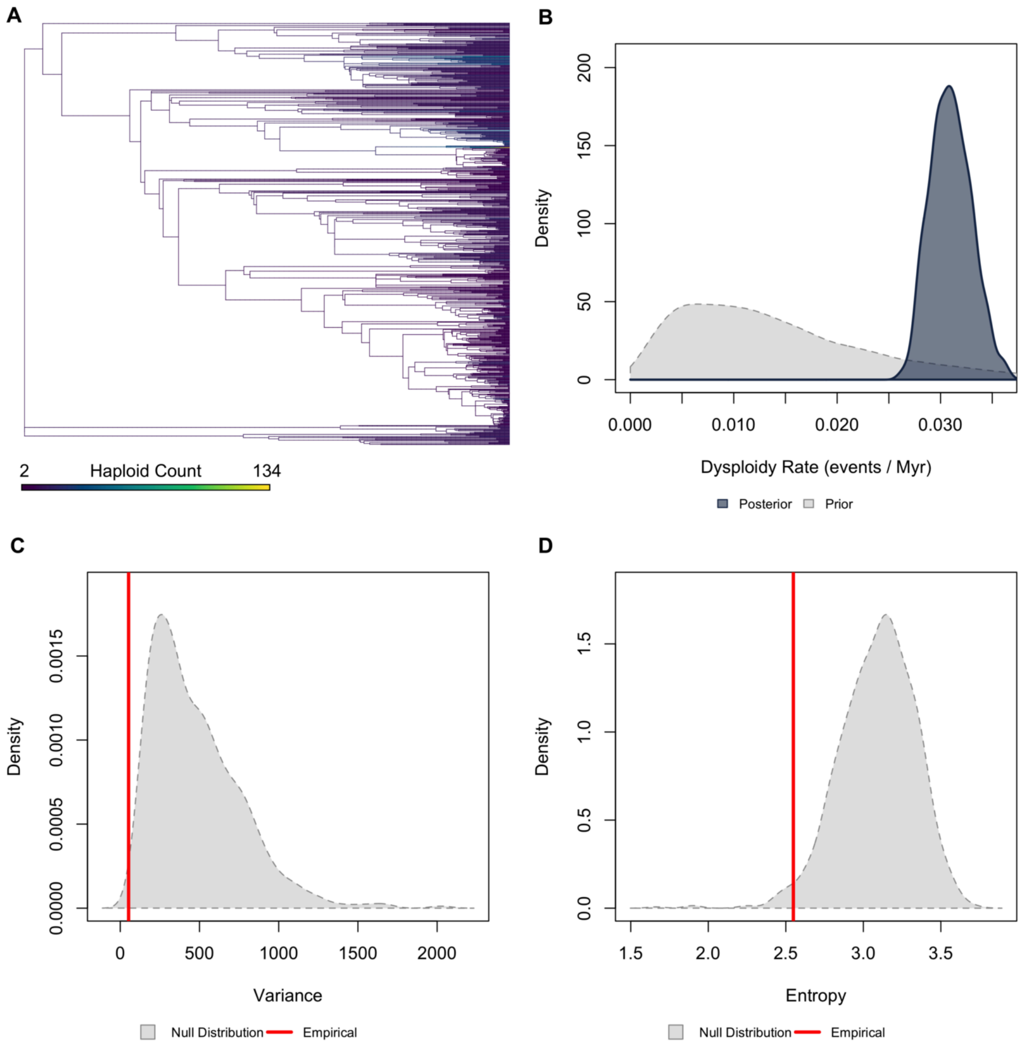


**Figure S10.** ***Validation plots for Fabaceae.*** *(A) Continuous character map of haploid chromosome number across the phylogeny. (B) Prior versus posterior density distributions for dysploidy rate. (C) Posterior predictive simulation adequacy test for variance. (D) Posterior predictive simulation adequacy test for Shannon's entropy.*

Liliaceae | Higher Taxonomy: Angiospermae | Chromosomes sampled: 648 | Phylogenetic tips: 79,340 | Overlap with phylogeny: 430 species | Root age: 119 Ma | Taxonomic tips: TRUE | Unresolved: 18.14%

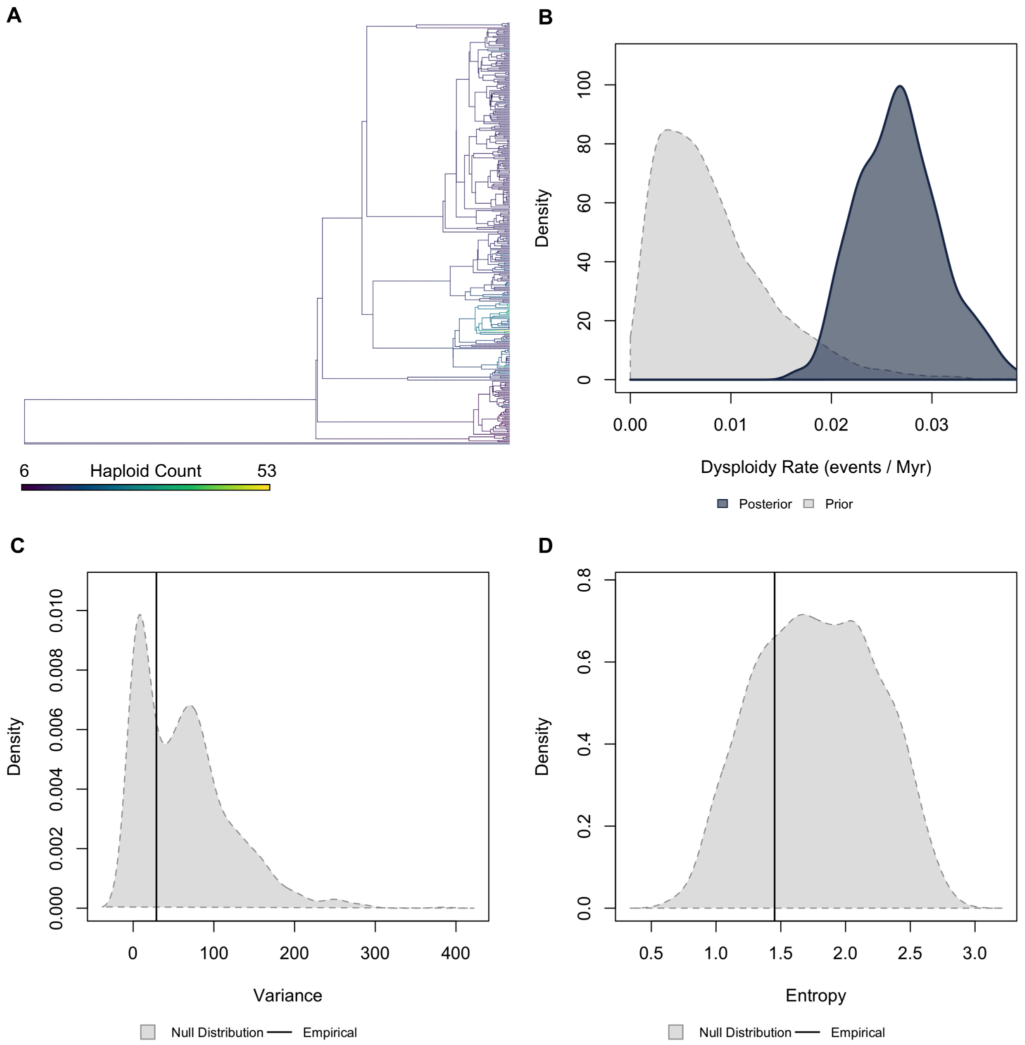


**Figure S11.** ***Validation plots for Liliaceae.*** *(A) Continuous character map of haploid chromosome number across the phylogeny. (B) Prior versus posterior density distributions for dysploidy rate. (C) Posterior predictive simulation adequacy test for variance. (D) Posterior predictive simulation adequacy test for Shannon's entropy.*

Magnoliaceae | Higher Taxonomy: Angiospermae | Chromosomes sampled: 139 | Phylogenetic tips: 304 | Overlap with phylogeny: 77 species | Root age: 42 Ma | Taxonomic tips: TRUE | Unresolved: 59.74%

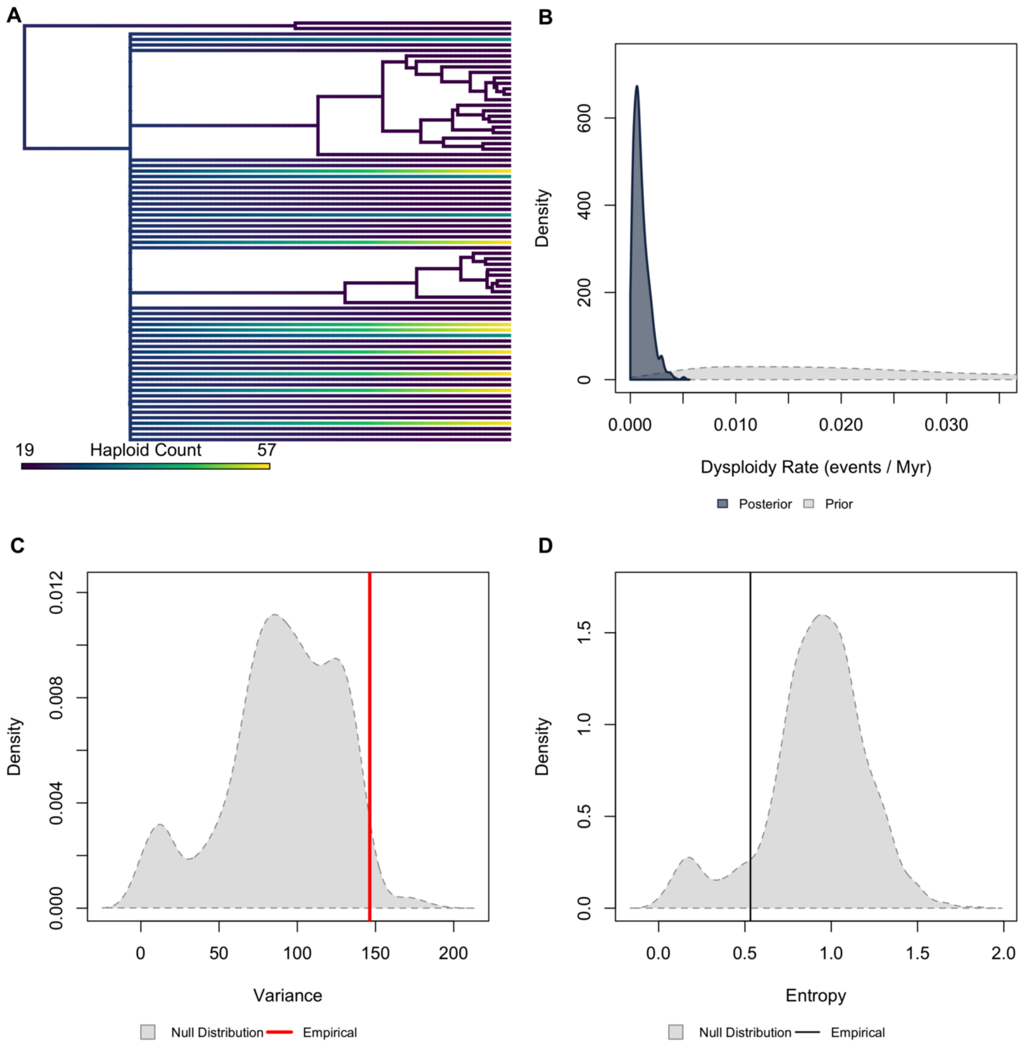


**Figure S12.** ***Validation plots for Magnoliaceae.*** *(A) Continuous character map of haploid chromosome number across the phylogeny. (B) Prior versus posterior density distributions for dysploidy rate. (C) Posterior predictive simulation adequacy test for variance. (D) Posterior predictive simulation adequacy test for Shannon's entropy.*

Note the tree topology in panel A is indicative of most species being placed on the tree via taxonomy which may lead to biases in rate estimates for that reason Magnoliaceae was dropped from the final results reported in the main text.

Orchidaceae | Higher Taxonomy: Angiospermae | Chromosomes sampled: 8,122 | Phylogenetic tips: 41,474 | Overlap with phylogeny: 1,904 species | Root age: 17 Ma | Taxonomic tips: TRUE | Unresolved: 43.22%

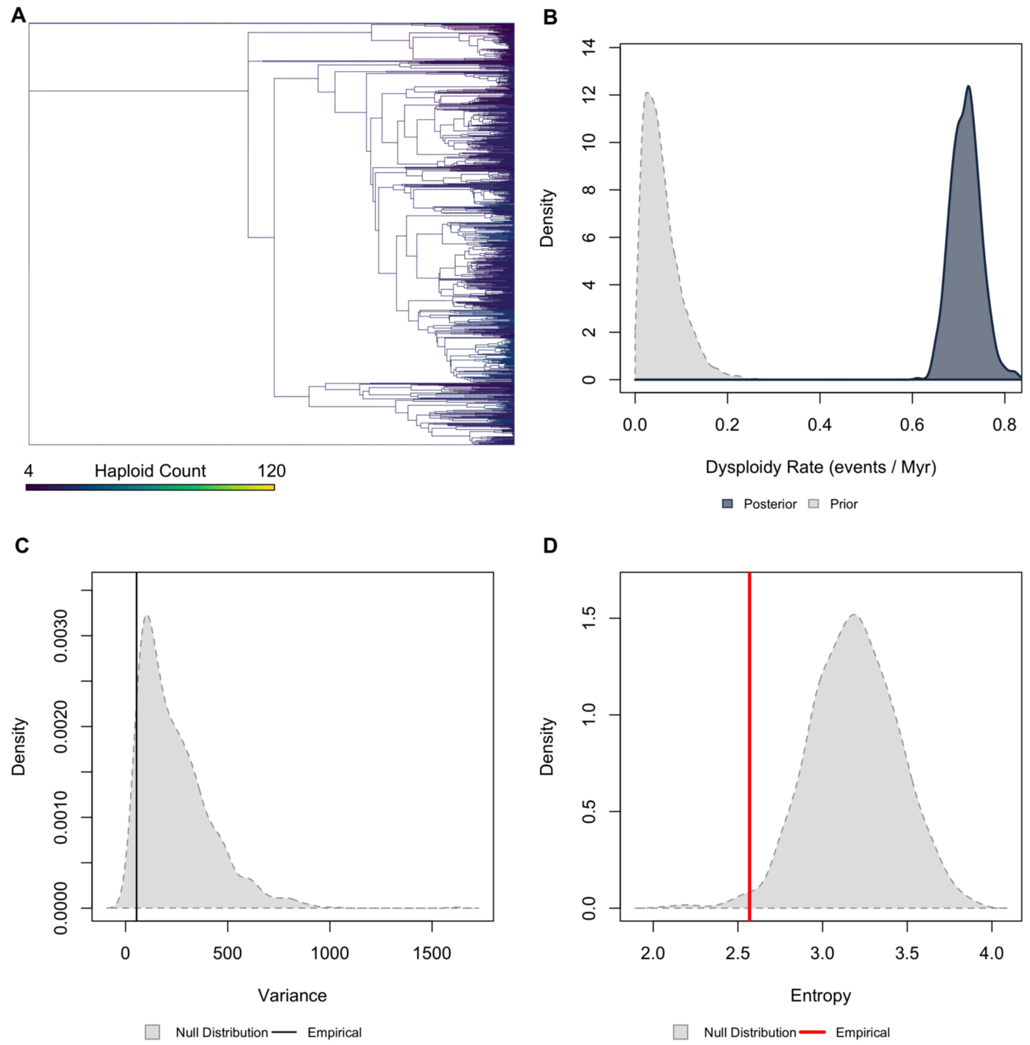


**Figure S13.** ***Validation plots for Orchidaceae.*** *(A) Continuous character map of haploid chromosome number across the phylogeny. (B) Prior versus posterior density distributions for dysploidy rate. (C) Posterior predictive simulation adequacy test for variance. (D) Posterior predictive simulation adequacy test for Shannon's entropy.*

Passifloraceae | Higher Taxonomy: Angiospermae | Chromosomes sampled: 159 | Phylogenetic tips: 134 | Overlap with phylogeny: 134 species | Root age: 50 Ma | Taxonomic tips: TRUE | Unresolved: 26.12%

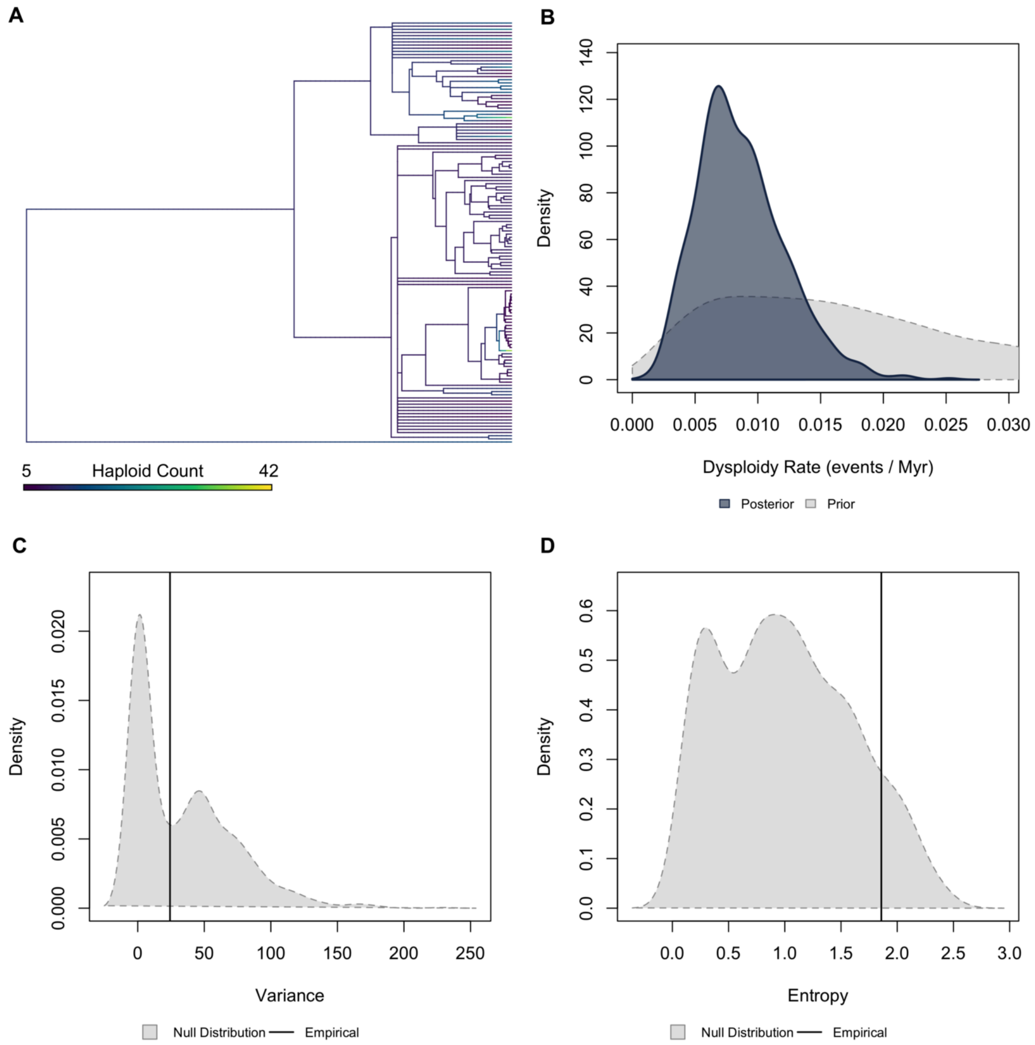


**Figure S14.** ***Validation plots for Passifloraceae.*** *(A) Continuous character map of haploid chromosome number across the phylogeny. (B) Prior versus posterior density distributions for dysploidy rate. (C) Posterior predictive simulation adequacy test for variance. (D) Posterior predictive simulation adequacy test for Shannon's entropy.*

Rubiaceae | Higher Taxonomy: Angiospermae | Chromosomes sampled: 1,219 | Phylogenetic tips: 794 | Overlap with phylogeny: 794 species | Root age: 72 Ma | Taxonomic tips: TRUE | Unresolved: 53.90%

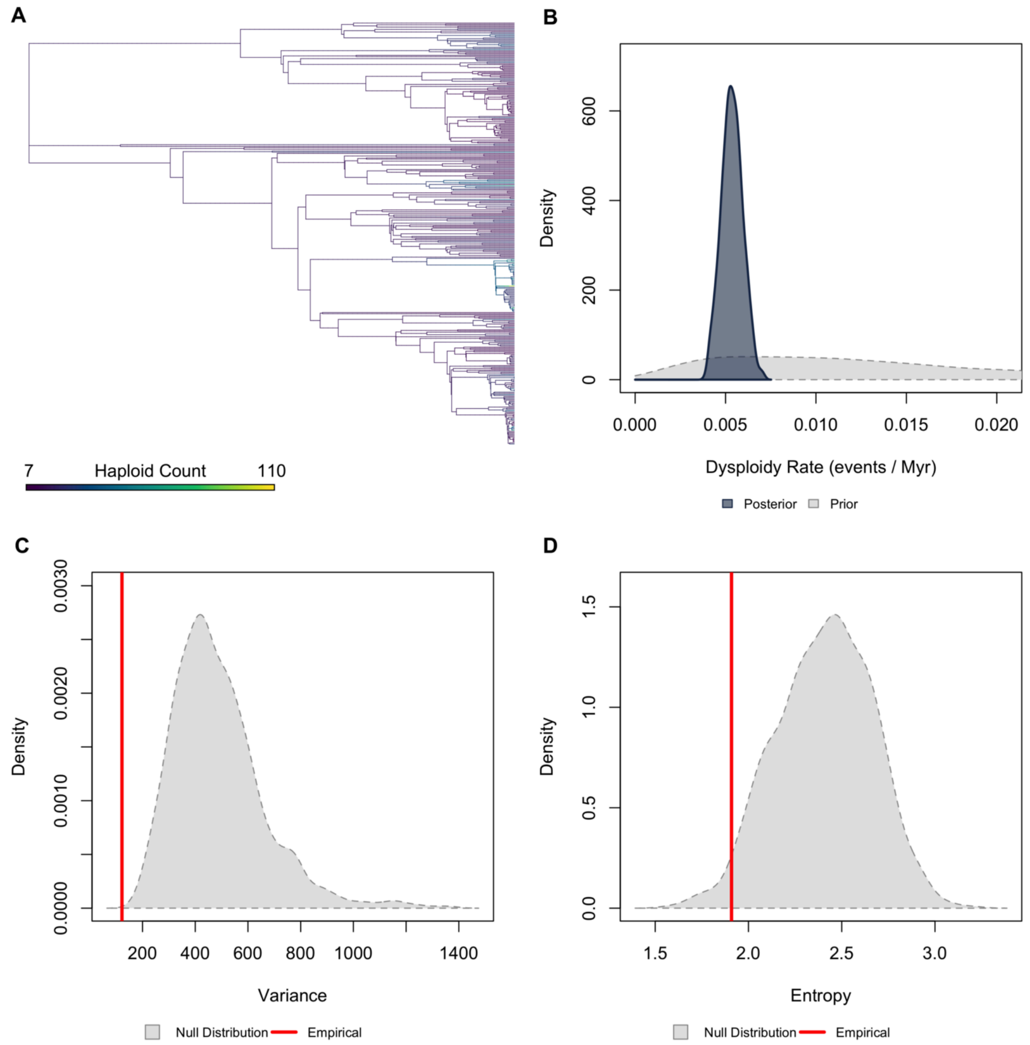


**Figure S15.** ***Validation plots for Rubiaceae.*** *(A) Continuous character map of haploid chromosome number across the phylogeny. (B) Prior versus posterior density distributions for dysploidy rate. (C) Posterior predictive simulation adequacy test for variance. (D) Posterior predictive simulation adequacy test for Shannon's entropy.*

Solanaceae | Higher Taxonomy: Angiospermae | Chromosomes sampled: 1,458 | Phylogenetic tips: 930 | Overlap with phylogeny: 930 species | Root age: 51 Ma | Taxonomic tips: TRUE | Unresolved: 48.28%

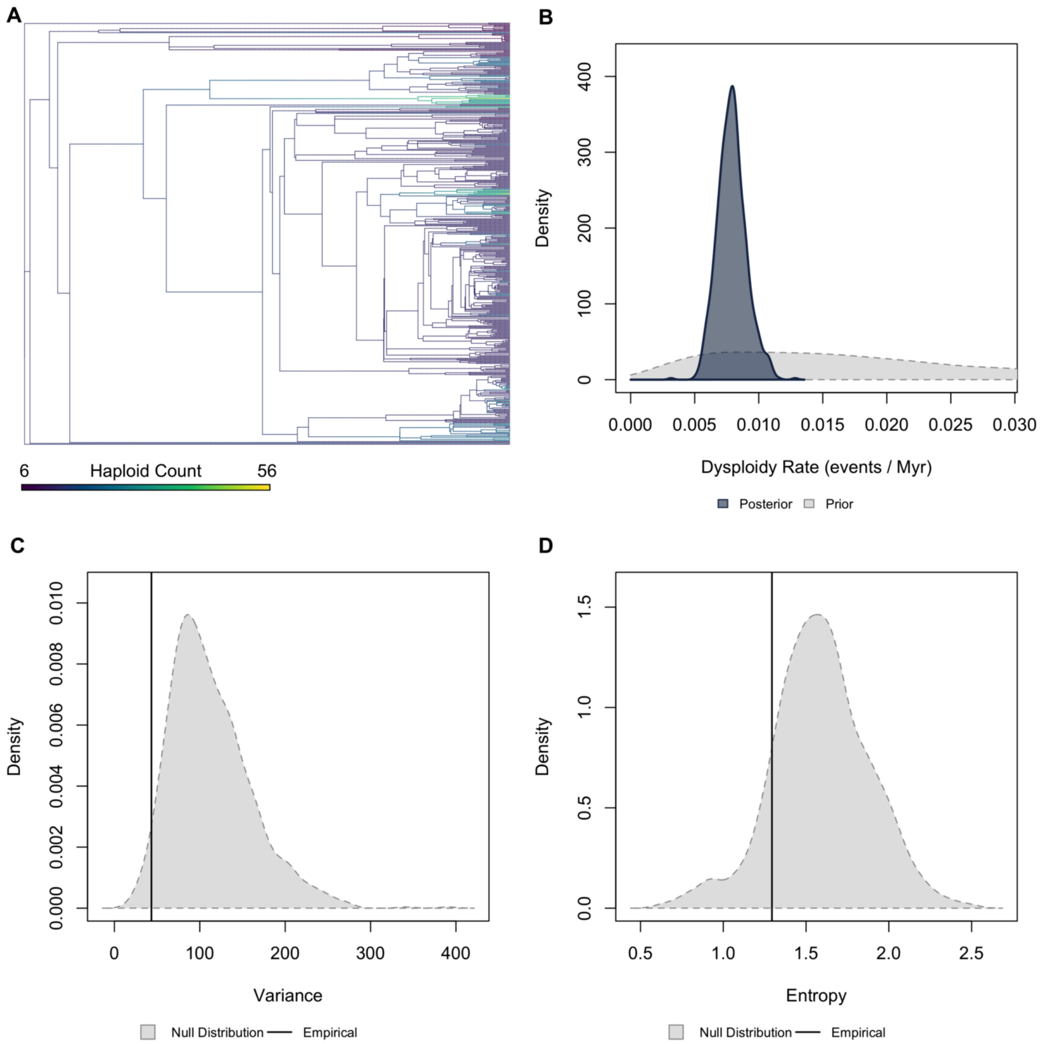


**Figure S16.** ***Validation plots for Solanaceae.*** *(A) Continuous character map of haploid chromosome number across the phylogeny. (B) Prior versus posterior density distributions for dysploidy rate. (C) Posterior predictive simulation adequacy test for variance. (D) Posterior predictive simulation adequacy test for Shannon's entropy.*

Fungi | Higher Taxonomy: Fungi | Chromosomes sampled: 151 | Phylogenetic tips: 5,418 | Overlap with phylogeny: 40 species | Root age: 642 Ma | Taxonomic tips: FALSE | Unresolved: N/A

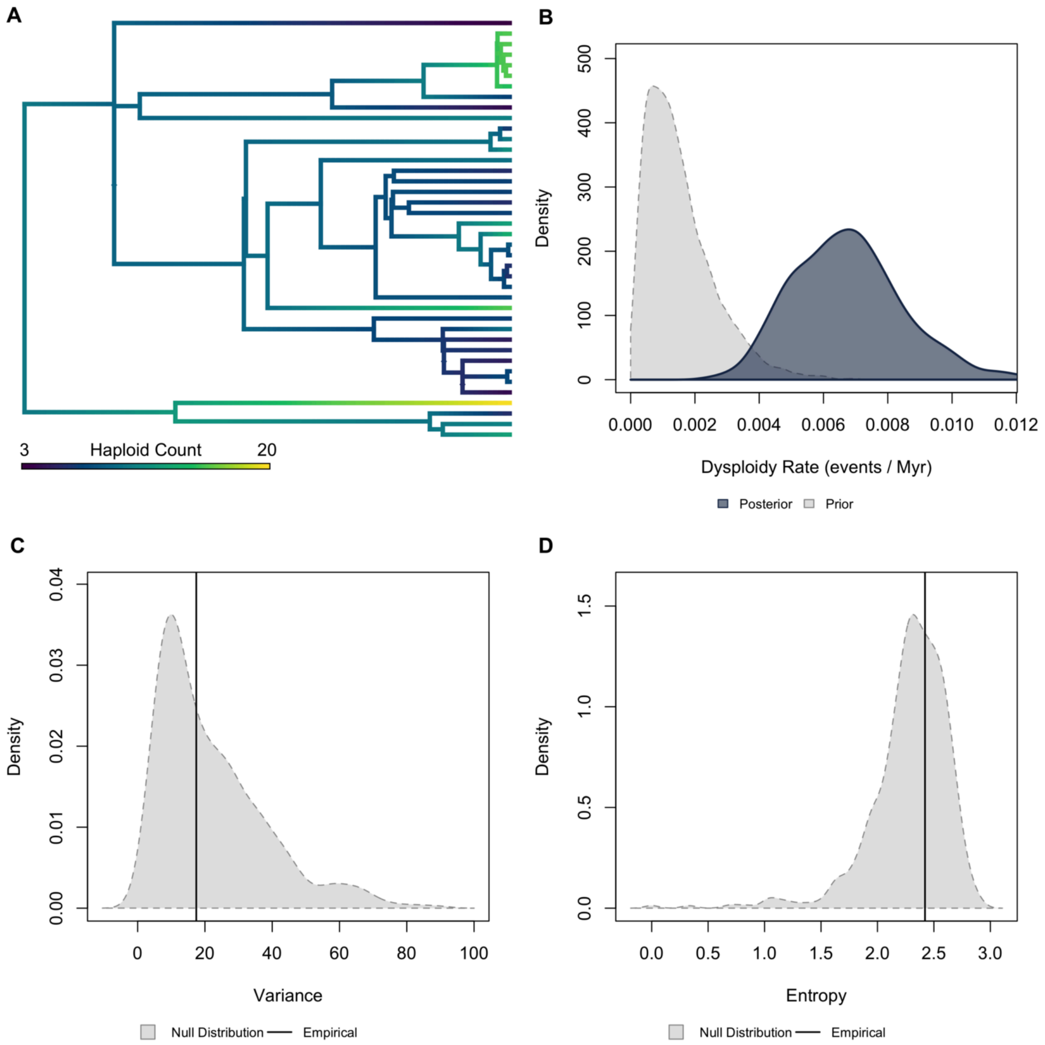


**Figure S17.** ***Validation plots for Fungi.*** *(A) Continuous character map of haploid chromosome number across the phylogeny. (B) Prior versus posterior density distributions for dysploidy rate. (C) Posterior predictive simulation adequacy test for variance. (D) Posterior predictive simulation adequacy test for Shannon's entropy.*

Blattodea | Higher Taxonomy: Insecta | Chromosomes sampled: 190 | Phylogenetic tips: 151 | Overlap with phylogeny: 30 species | Root age: 270 Ma | Taxonomic tips: FALSE | Unresolved: N/A

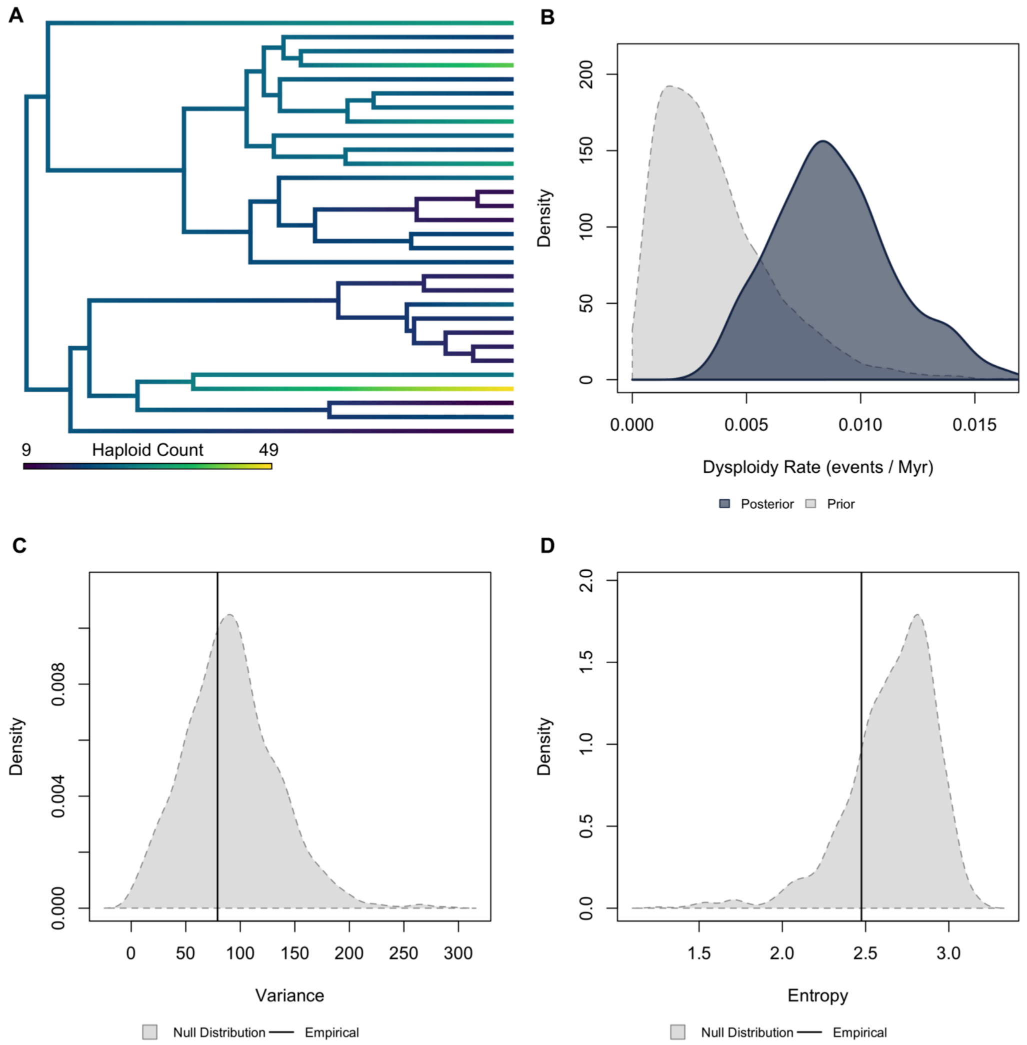


**Figure S18.** ***Validation plots for Blattodea.*** *(A) Continuous character map of haploid chromosome number across the phylogeny. (B) Prior versus posterior density distributions for dysploidy rate. (C) Posterior predictive simulation adequacy test for variance. (D) Posterior predictive simulation adequacy test for Shannon's entropy.*

Phasmatodea | Higher Taxonomy: Insecta | Chromosomes sampled: 109 | Phylogenetic tips: 148 | Overlap with phylogeny: 11 species | Root age: 140 Ma | Taxonomic tips: FALSE | Unresolved: N/A

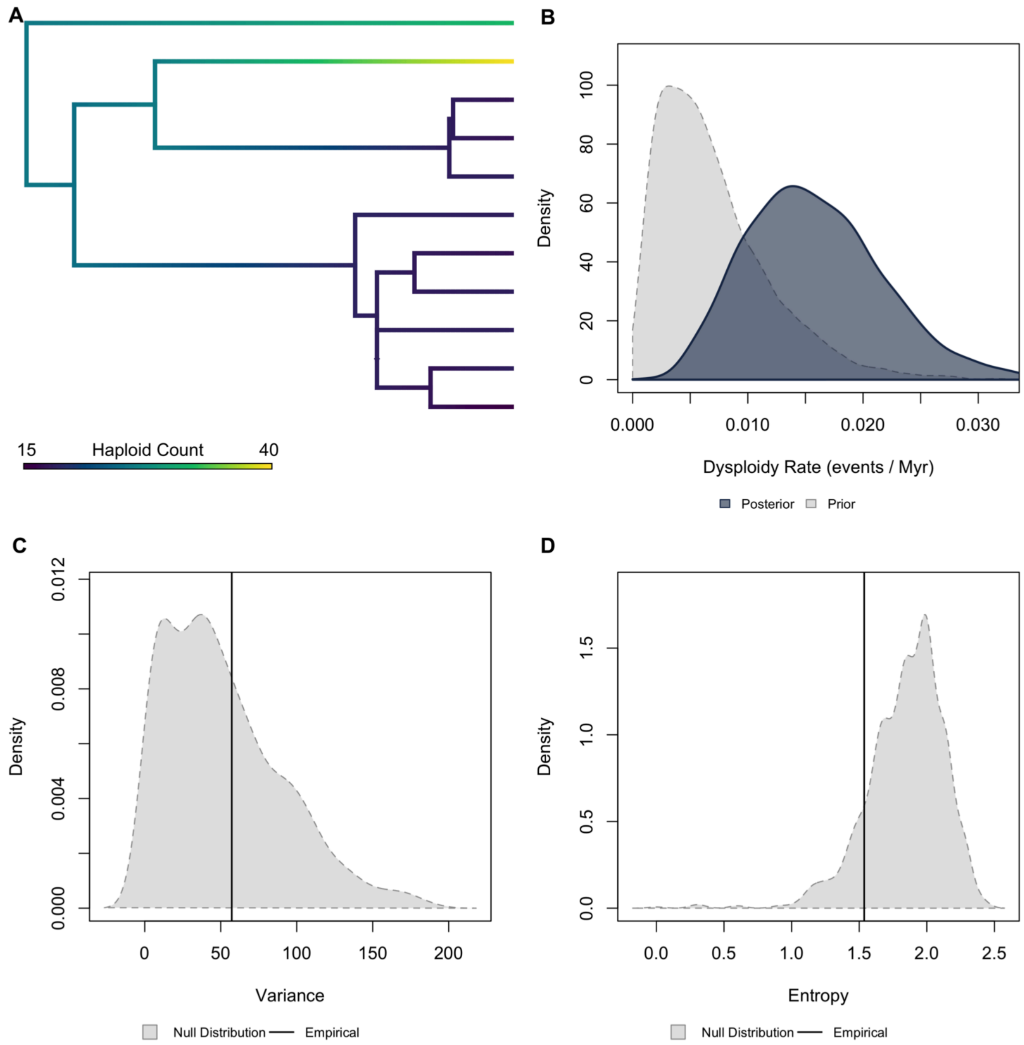


**Figure S19.** ***Validation plots for Phasmatodea.*** *(A) Continuous character map of haploid chromosome number across the phylogeny. (B) Prior versus posterior density distributions for dysploidy rate. (C) Posterior predictive simulation adequacy test for variance. (D) Posterior predictive simulation adequacy test for Shannon's entropy.*

Orthoptera | Higher Taxonomy: Insecta | Chromosomes sampled: 319 | Phylogenetic tips: 232 | Overlap with phylogeny: 36 species | Root age: 74 Ma | Taxonomic tips: FALSE | Unresolved: N/A

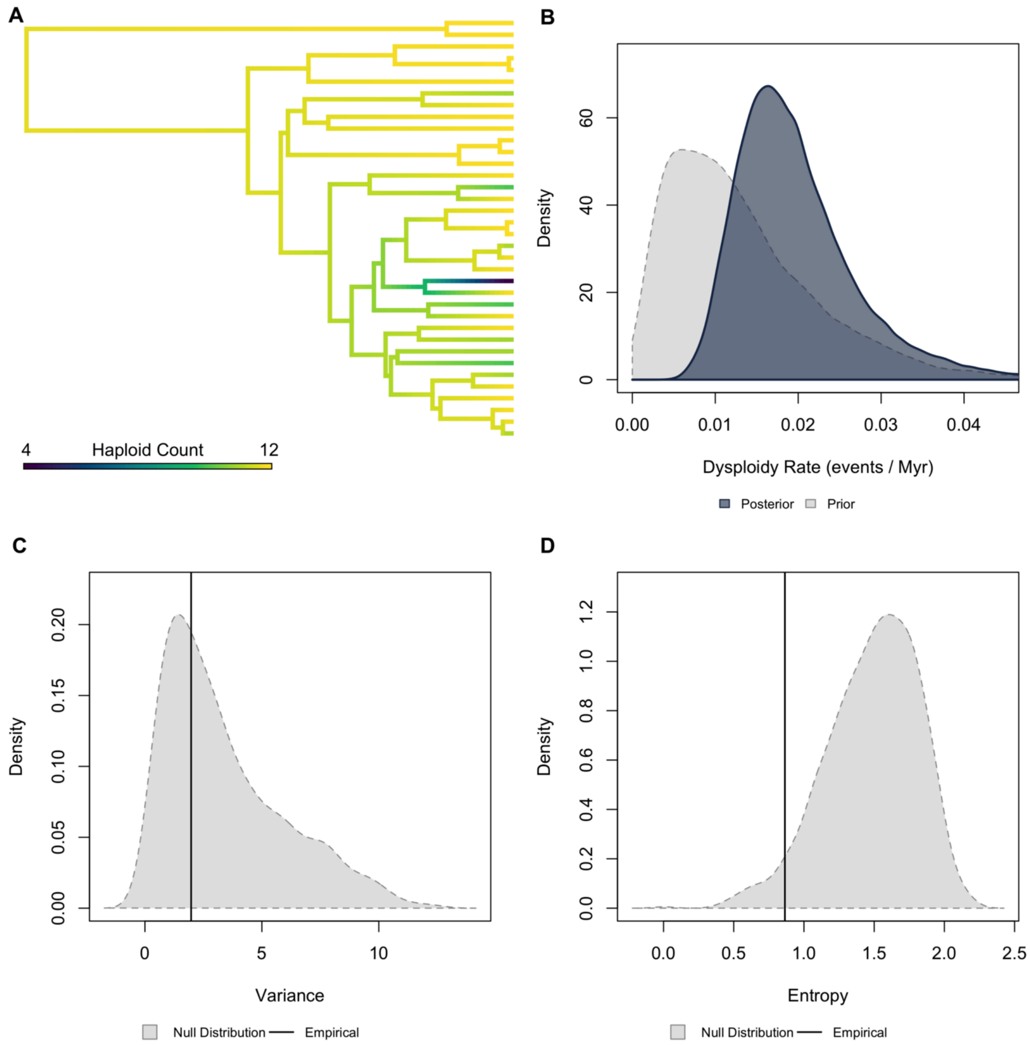


**Figure S20.** ***Validation plots for Orthoptera.*** *(A) Continuous character map of haploid chromosome number across the phylogeny. (B) Prior versus posterior density distributions for dysploidy rate. (C) Posterior predictive simulation adequacy test for variance. (D) Posterior predictive simulation adequacy test for Shannon's entropy.*

Hemiptera | Higher Taxonomy: Insecta | Chromosomes sampled: 1,744 | Phylogenetic tips: 1,967 | Overlap with phylogeny: 46 species | Root age: 258 Ma | Taxonomic tips: FALSE | Unresolved: N/A

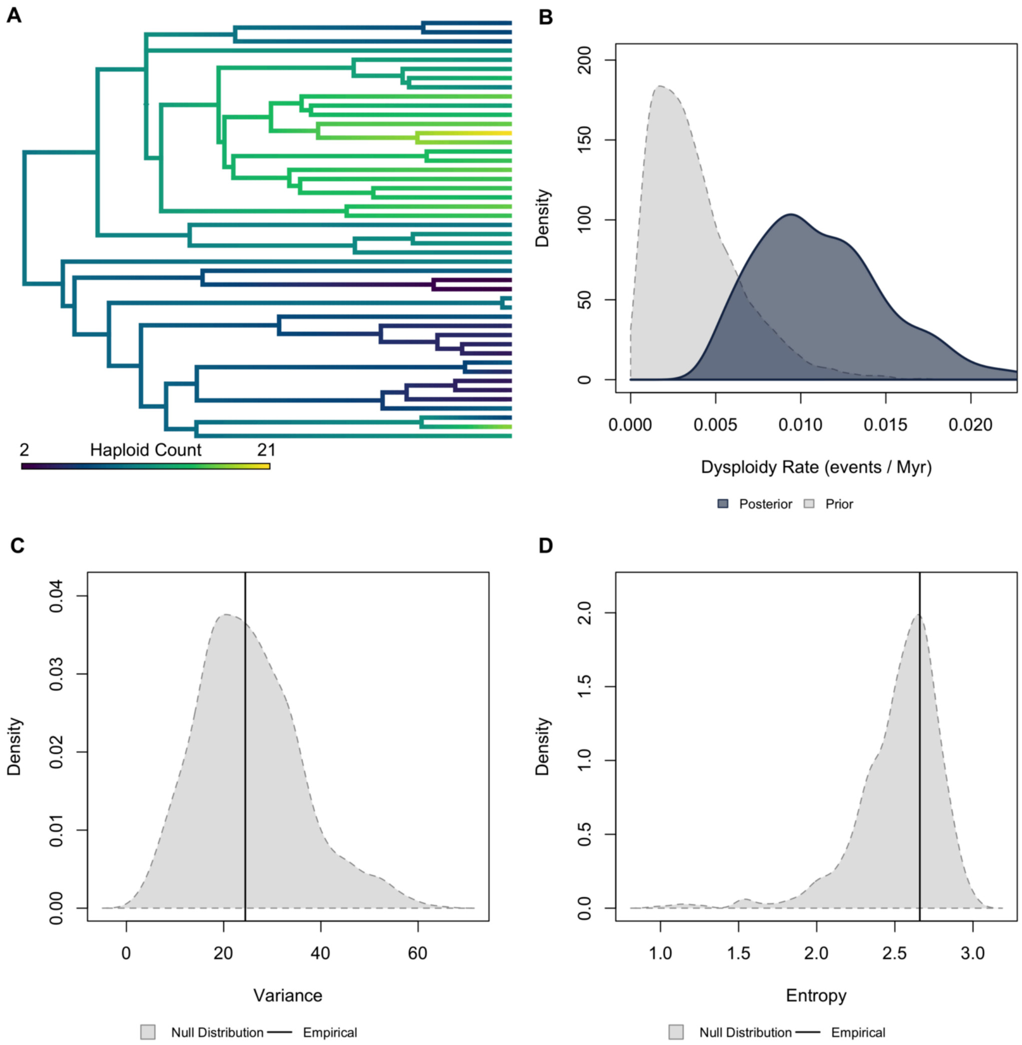


**Figure S21.** ***Validation plots for Hemiptera.*** *(A) Continuous character map of haploid chromosome number across the phylogeny. (B) Prior versus posterior density distributions for dysploidy rate. (C) Posterior predictive simulation adequacy test for variance. (D) Posterior predictive simulation adequacy test for Shannon's entropy.*

Hymenoptera | Higher Taxonomy: Insecta | Chromosomes sampled: 1,591 | Phylogenetic tips: 602 | Overlap with phylogeny: 345 species | Root age: 240 Ma | Taxonomic tips: FALSE | Unresolved: N/A

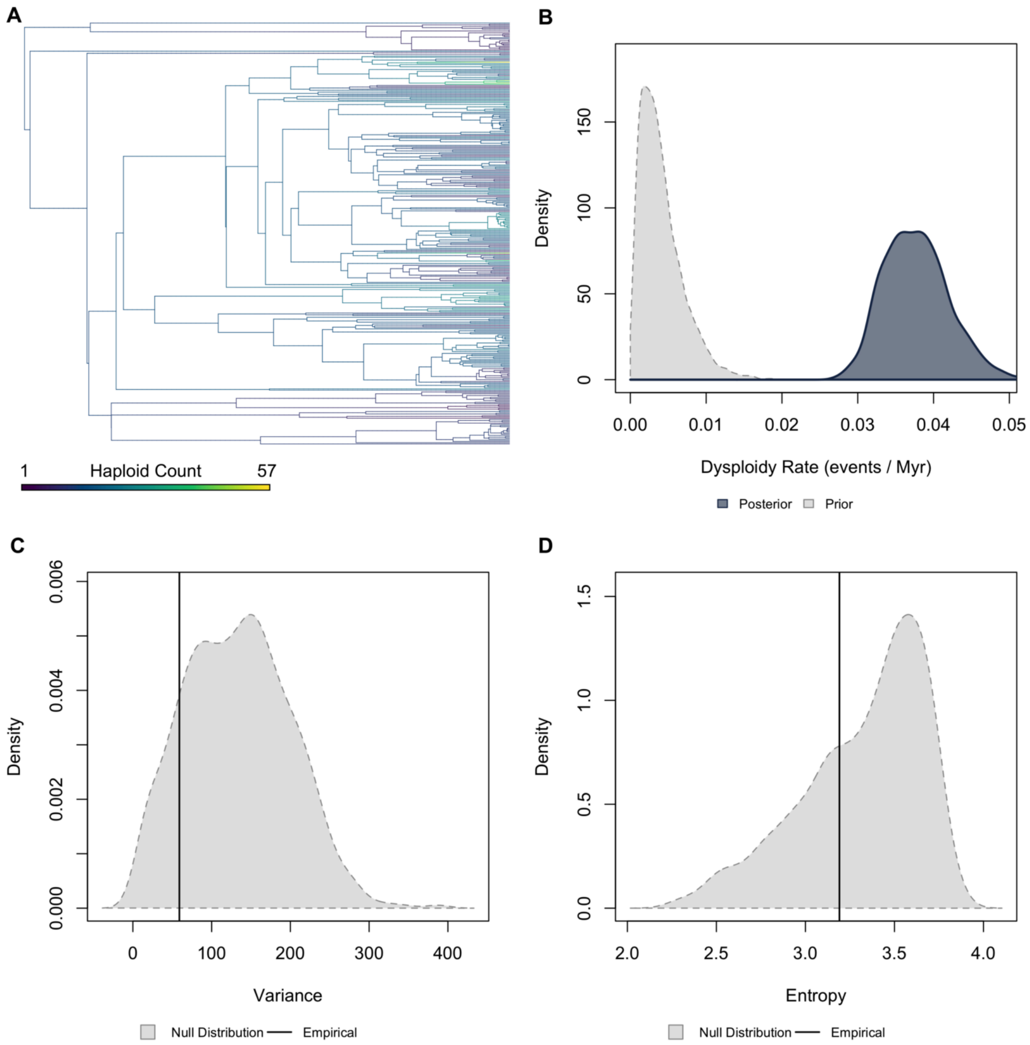


**Figure S22.** ***Validation plots for Hymenoptera.*** *(A) Continuous character map of haploid chromosome number across the phylogeny. (B) Prior versus posterior density distributions for dysploidy rate. (C) Posterior predictive simulation adequacy test for variance. (D) Posterior predictive simulation adequacy test for Shannon's entropy.*

Dytiscidae | Higher Taxonomy: Insecta | Chromosomes sampled: 85 | Phylogenetic tips: 973 | Overlap with phylogeny: 29 species | Root age: 144 Ma | Taxonomic tips: FALSE | Unresolved: N/A

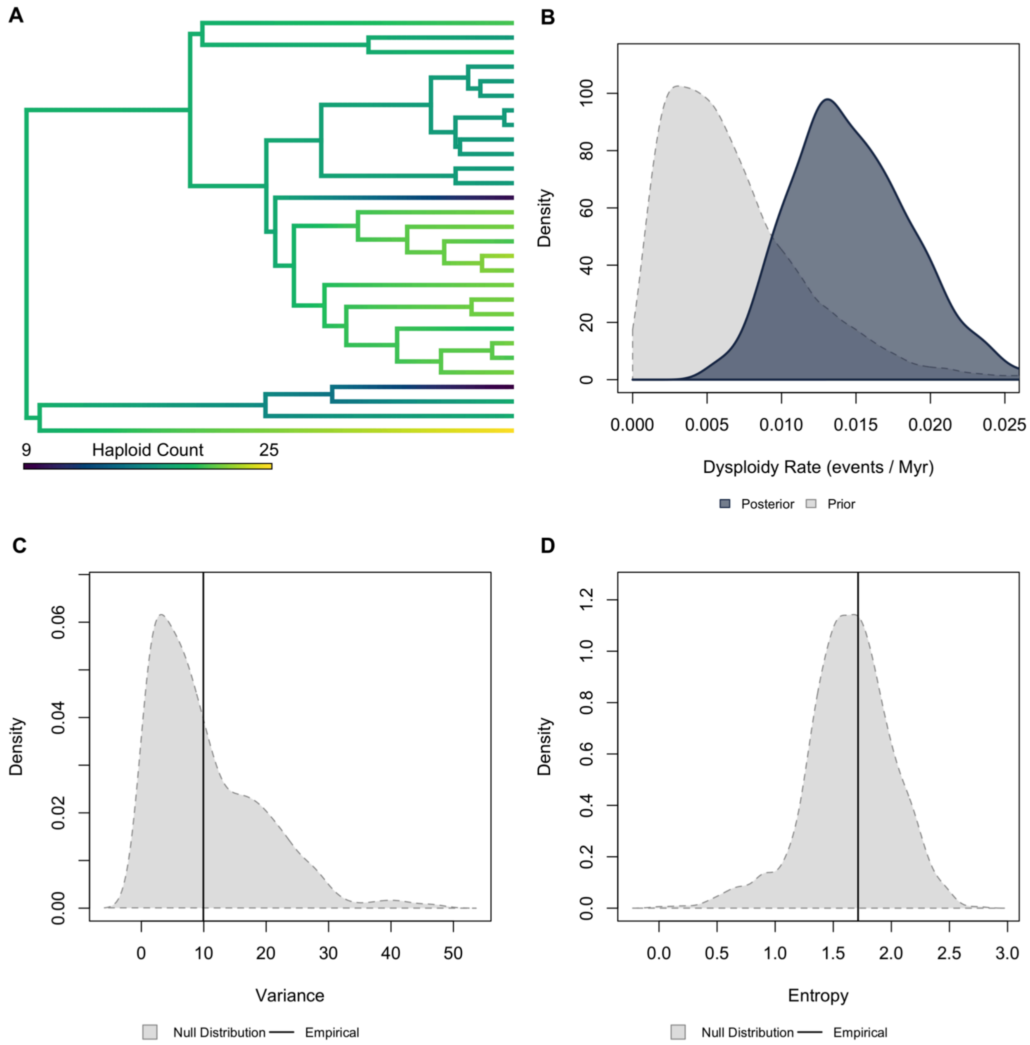


**Figure S23. *Validation plots for Dytiscidae.*** *(A) Continuous character map of haploid chromosome number across the phylogeny. (B) Prior versus posterior density distributions for dysploidy rate. (C) Posterior predictive simulation adequacy test for variance. (D) Posterior predictive simulation adequacy test for Shannon's entropy.*

Carabidae | Higher Taxonomy: Insecta | Chromosomes sampled: 777 | Phylogenetic tips: 104 | Overlap with phylogeny: 95 species | Root age: 103 Ma | Taxonomic tips: FALSE | Unresolved: N/A

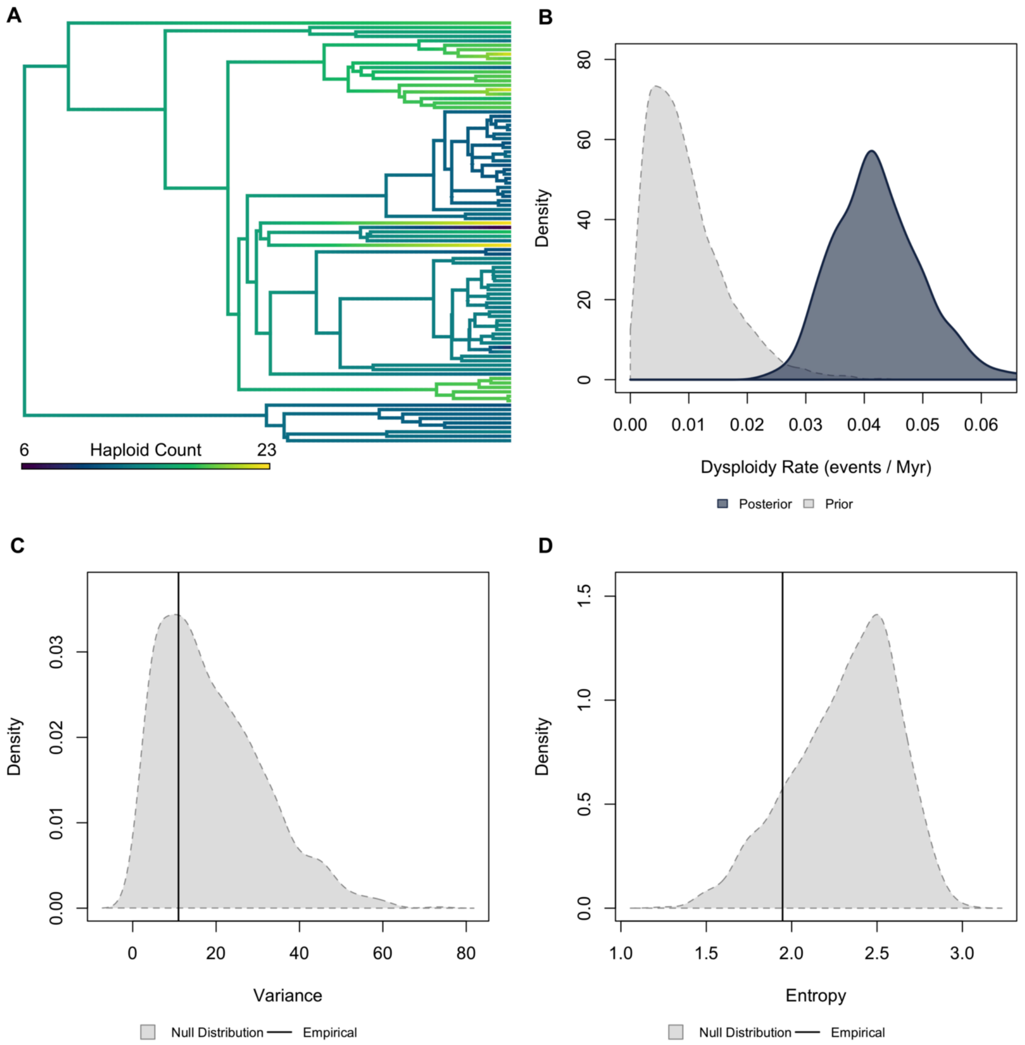


**Figure S24.** ***Validation plots for Carabidae.*** *(A) Continuous character map of haploid chromosome number across the phylogeny. (B) Prior versus posterior density distributions for dysploidy rate. (C) Posterior predictive simulation adequacy test for variance. (D) Posterior predictive simulation adequacy test for Shannon's entropy.*

Hydrophilidae | Higher Taxonomy: Insecta | Chromosomes sampled: 81 | Phylogenetic tips: 168 | Overlap with phylogeny: 21 species | Root age: 133 Ma | Taxonomic tips: FALSE | Unresolved: N/A

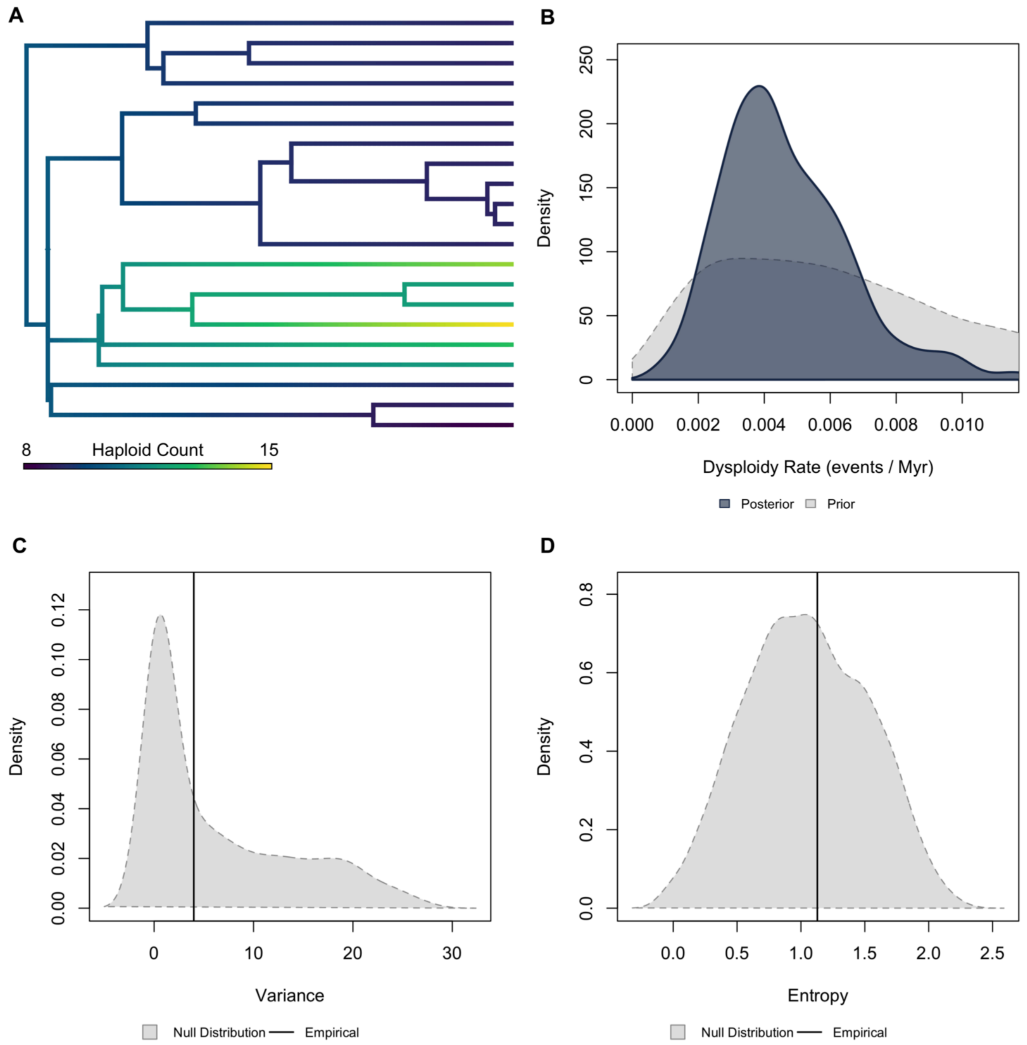


**Figure S25.** ***Validation plots for Hydrophilidae.*** *(A) Continuous character map of haploid chromosome number across the phylogeny. (B) Prior versus posterior density distributions for dysploidy rate. (C) Posterior predictive simulation adequacy test for variance. (D) Posterior predictive simulation adequacy test for Shannon's entropy.*

Scarabidae | Higher Taxonomy: Insecta | Chromosomes sampled: 478 | Phylogenetic tips: 211 | Overlap with phylogeny: 174 species | Root age: 180 Ma | Taxonomic tips: FALSE | Unresolved: N/A

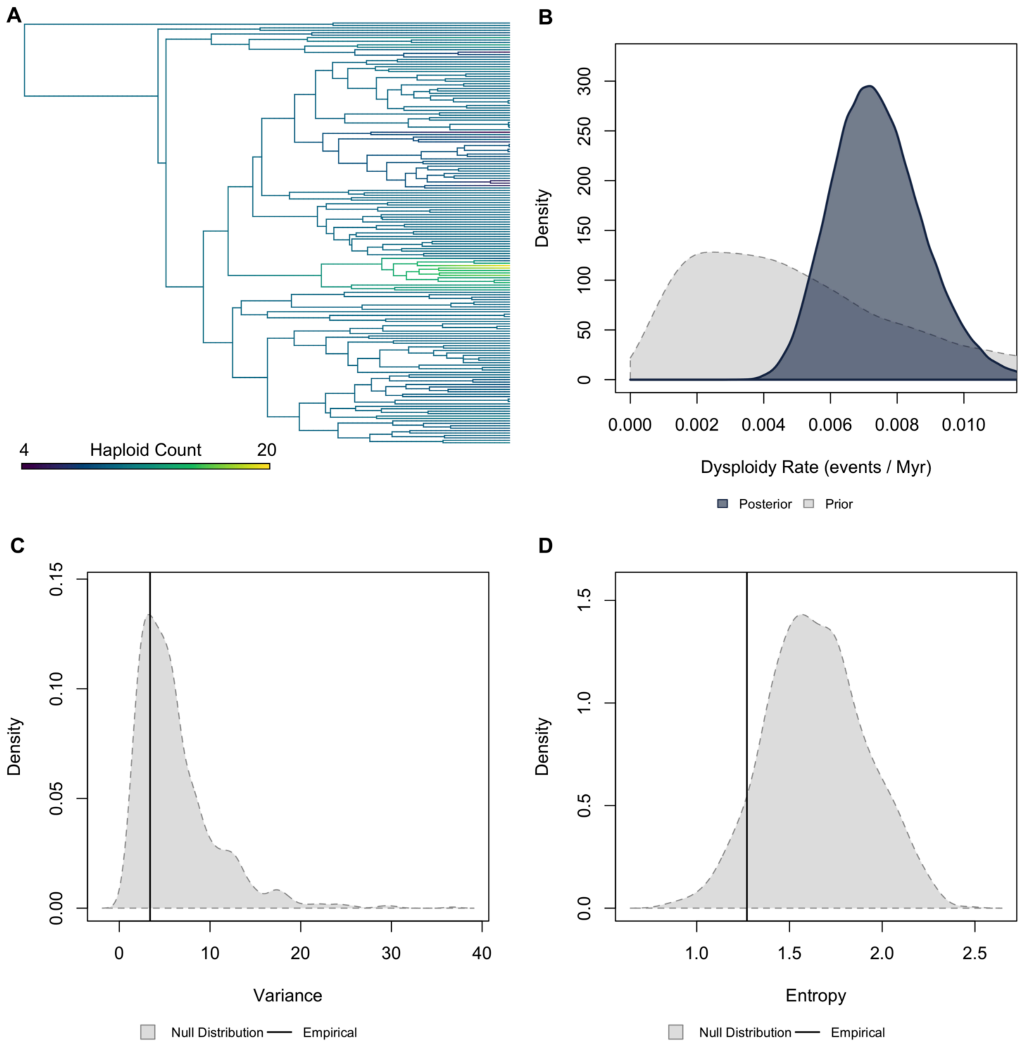


**Figure S26.** ***Validation plots for Scarabidae.*** *(A) Continuous character map of haploid chromosome number across the phylogeny. (B) Prior versus posterior density distributions for dysploidy rate. (C) Posterior predictive simulation adequacy test for variance. (D) Posterior predictive simulation adequacy test for Shannon's entropy.*

Coccinellidae | Higher Taxonomy: Insecta | Chromosomes sampled: 159 | Phylogenetic tips: 112 | Overlap with phylogeny: 24 species | Root age: 219 Ma | Taxonomic tips: FALSE | Unresolved: N/A

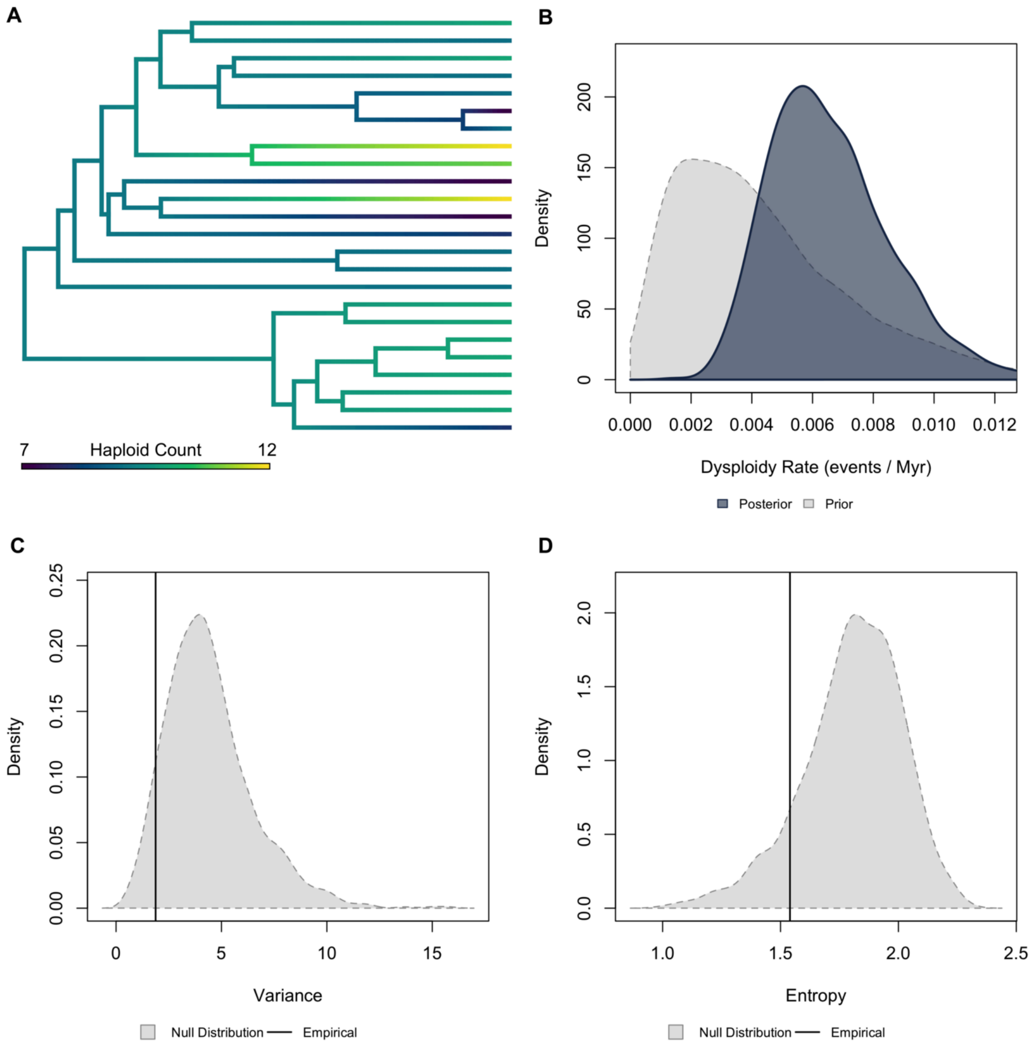


**Figure S27.** ***Validation plots for Coccinellidae.*** *(A) Continuous character map of haploid chromosome number across the phylogeny. (B) Prior versus posterior density distributions for dysploidy rate. (C) Posterior predictive simulation adequacy test for variance. (D) Posterior predictive simulation adequacy test for Shannon's entropy.*

Chrysomelidae | Higher Taxonomy: Insecta | Chromosomes sampled: 869 | Phylogenetic tips: 1,640 | Overlap with phylogeny: 182 species | Root age: 136 Ma | Taxonomic tips: FALSE | Unresolved: N/A

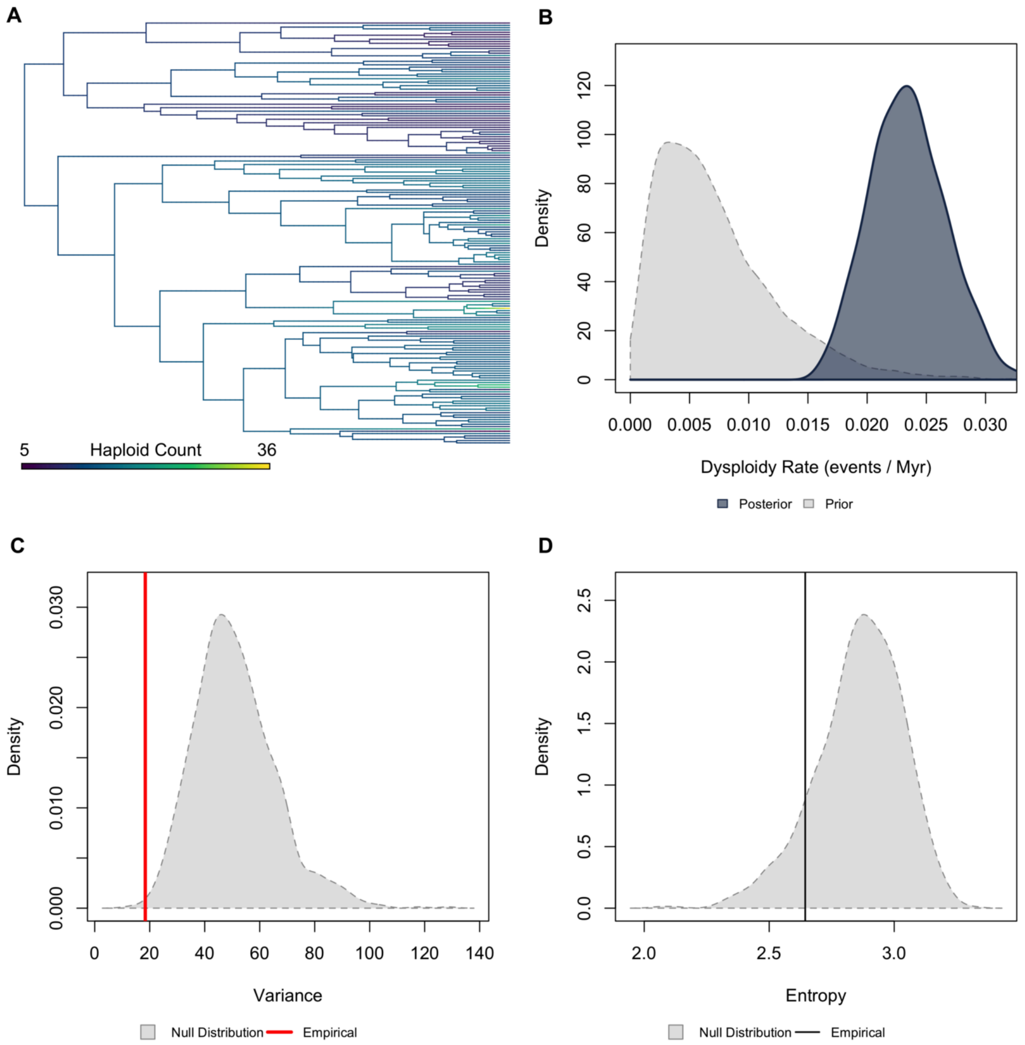


**Figure S28.** ***Validation plots for Chrysomelidae.*** *(A) Continuous character map of haploid chromosome number across the phylogeny. (B) Prior versus posterior density distributions for dysploidy rate. (C) Posterior predictive simulation adequacy test for variance. (D) Posterior predictive simulation adequacy test for Shannon's entropy.*

Curculionidae | Higher Taxonomy: Insecta | Chromosomes sampled: 617 | Phylogenetic tips: 1,492 | Overlap with phylogeny: 33 species | Root age: 95 Ma | Taxonomic tips: FALSE | Unresolved: N/A

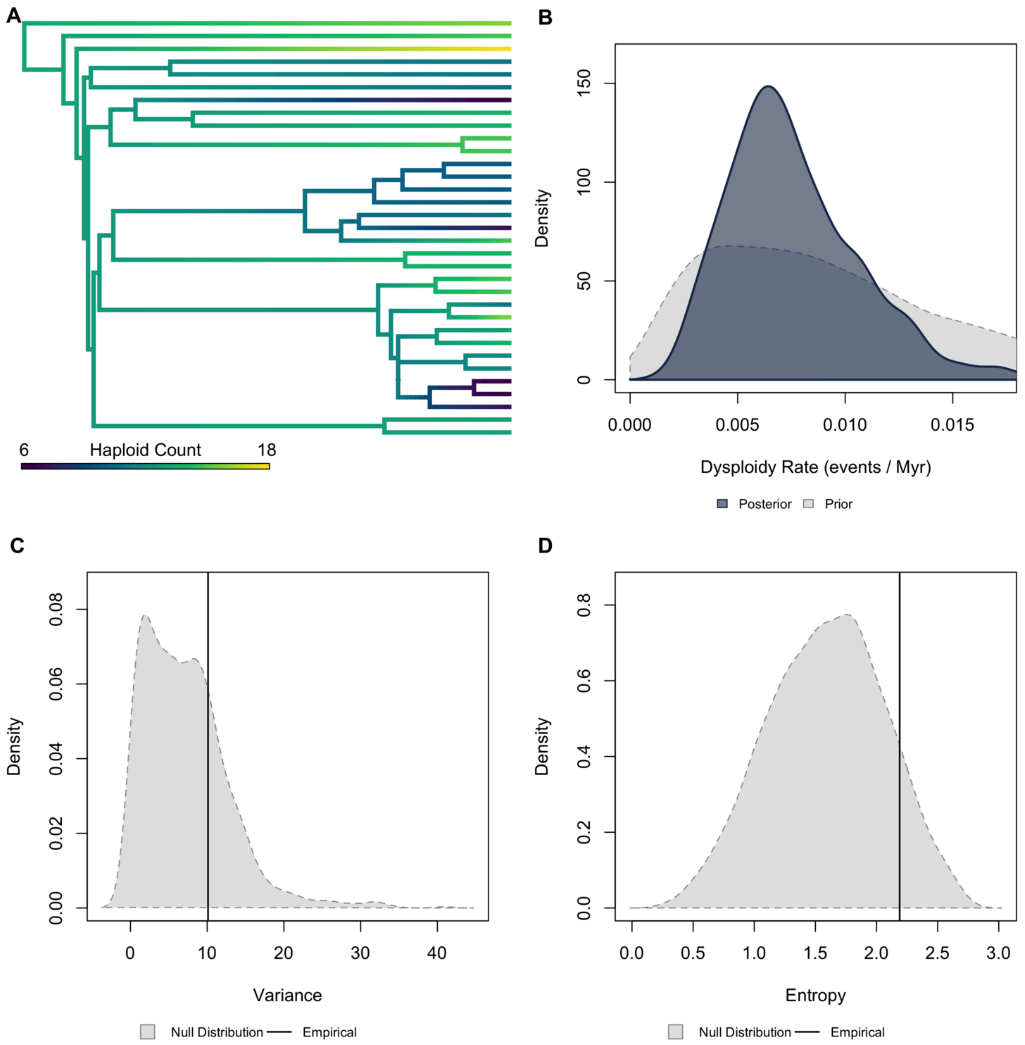


**Figure S29.** ***Validation plots for Curculionidae.*** *(A) Continuous character map of haploid chromosome number across the phylogeny. (B) Prior versus posterior density distributions for dysploidy rate. (C) Posterior predictive simulation adequacy test for variance. (D) Posterior predictive simulation adequacy test for Shannon's entropy.*

Tenebrionidae | Higher Taxonomy: Insecta | Chromosomes sampled: 239 | Phylogenetic tips: 318 | Overlap with phylogeny: 40 species | Root age: 180 Ma | Taxonomic tips: FALSE | Unresolved: N/A

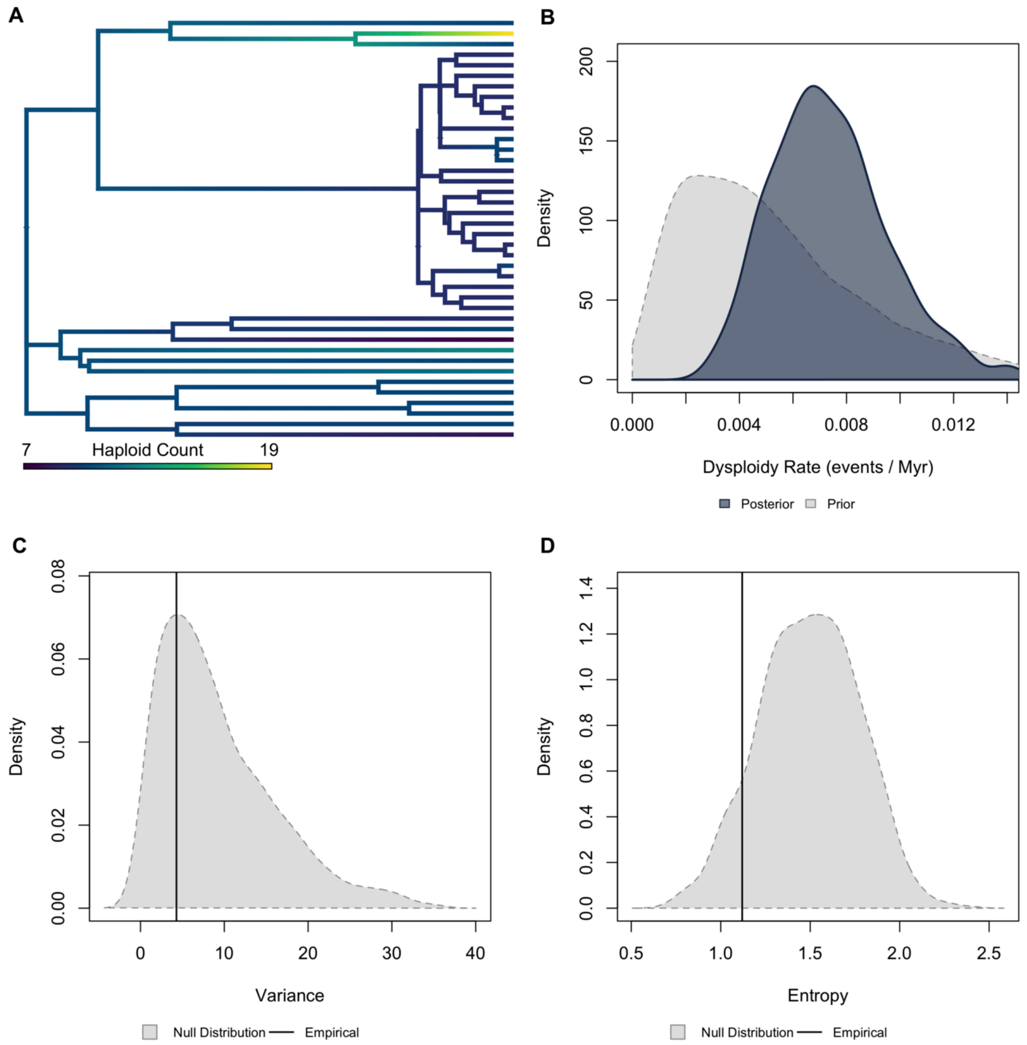


**Figure S30.** ***Validation plots for Tenebrionidae.*** *(A) Continuous character map of haploid chromosome number across the phylogeny. (B) Prior versus posterior density distributions for dysploidy rate. (C) Posterior predictive simulation adequacy test for variance. (D) Posterior predictive simulation adequacy test for Shannon's entropy.*

Drosophilidae | Higher Taxonomy: Insecta | Chromosomes sampled: 1,246 | Phylogenetic tips: 685 | Overlap with phylogeny: 352 species | Root age: 62 Ma | Taxonomic tips: FALSE | Unresolved: N/A

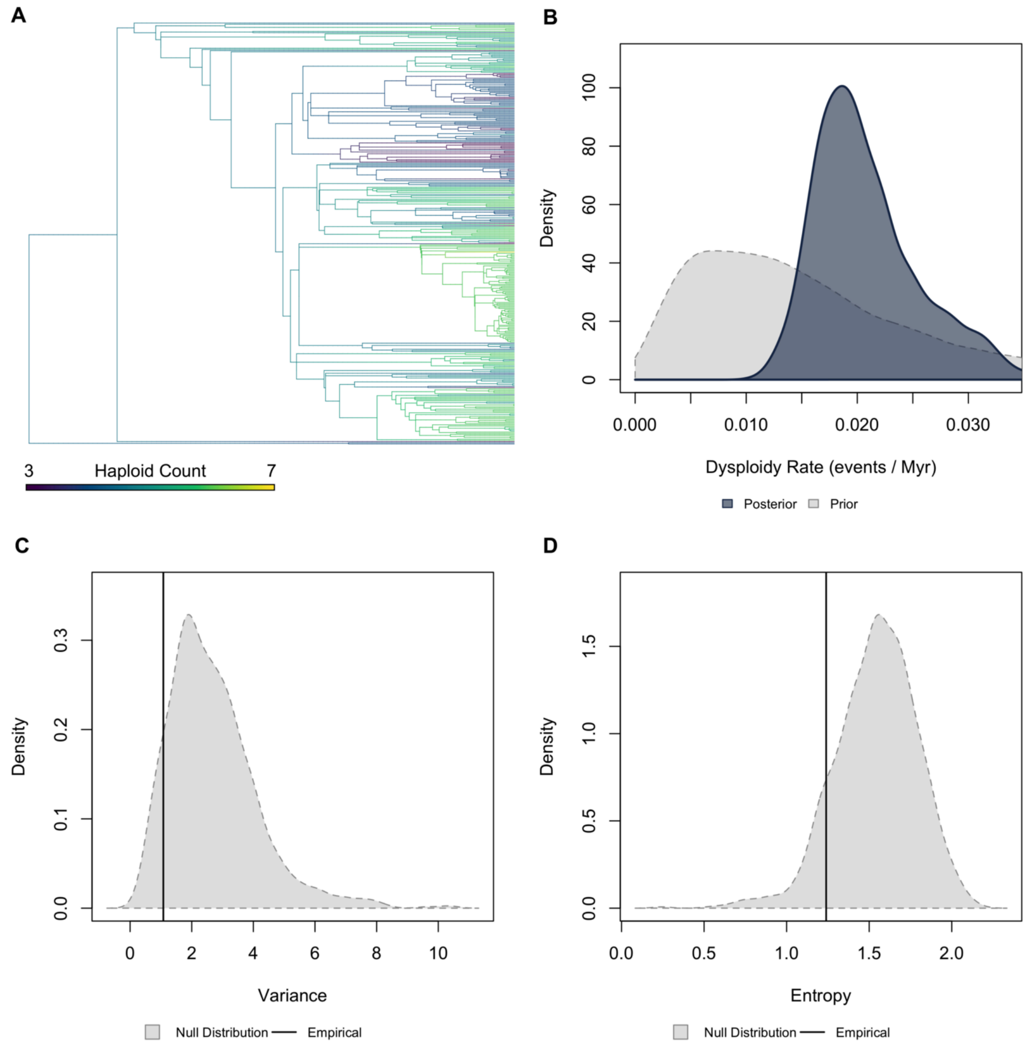


**Figure S31.** ***Validation plots for Drosophilidae.*** *(A) Continuous character map of haploid chromosome number across the phylogeny. (B) Prior versus posterior density distributions for dysploidy rate. (C) Posterior predictive simulation adequacy test for variance. (D) Posterior predictive simulation adequacy test for Shannon's entropy.*

Lepidoptera | Higher Taxonomy: Insecta | Chromosomes sampled: 3,370 | Phylogenetic tips: 2,255 | Overlap with phylogeny: 322 species | Root age: 102 Ma | Taxonomic tips: FALSE | Unresolved: N/A

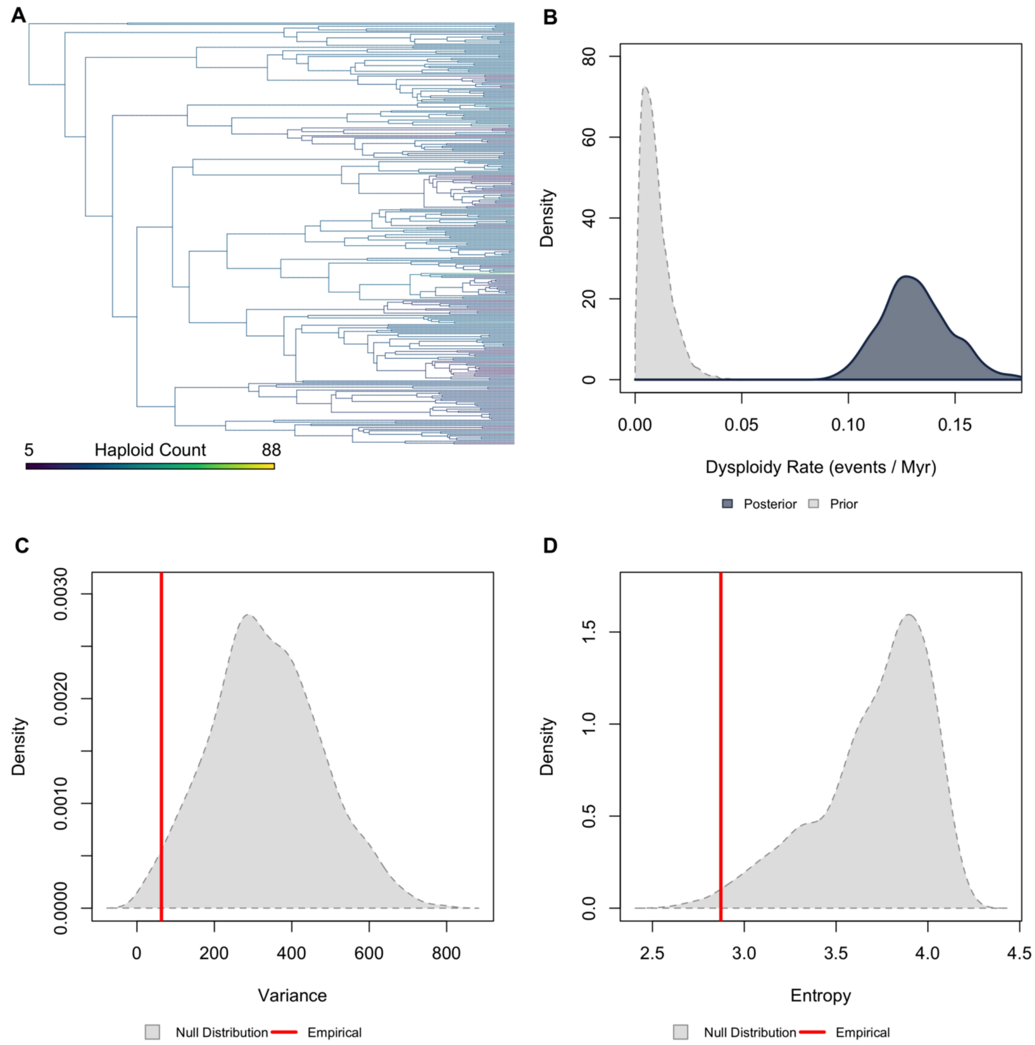


**Figure S32.** ***Validation plots for Lepidoptera.*** *(A) Continuous character map of haploid chromosome number across the phylogeny. (B) Prior versus posterior density distributions for dysploidy rate. (C) Posterior predictive simulation adequacy test for variance. (D) Posterior predictive simulation adequacy test for Shannon's entropy.*

Odonata | Higher Taxonomy: Insecta | Chromosomes sampled: 703 | Phylogenetic tips: 669 | Overlap with phylogeny: 84 species | Root age: 67 Ma | Taxonomic tips: FALSE | Unresolved: N/A

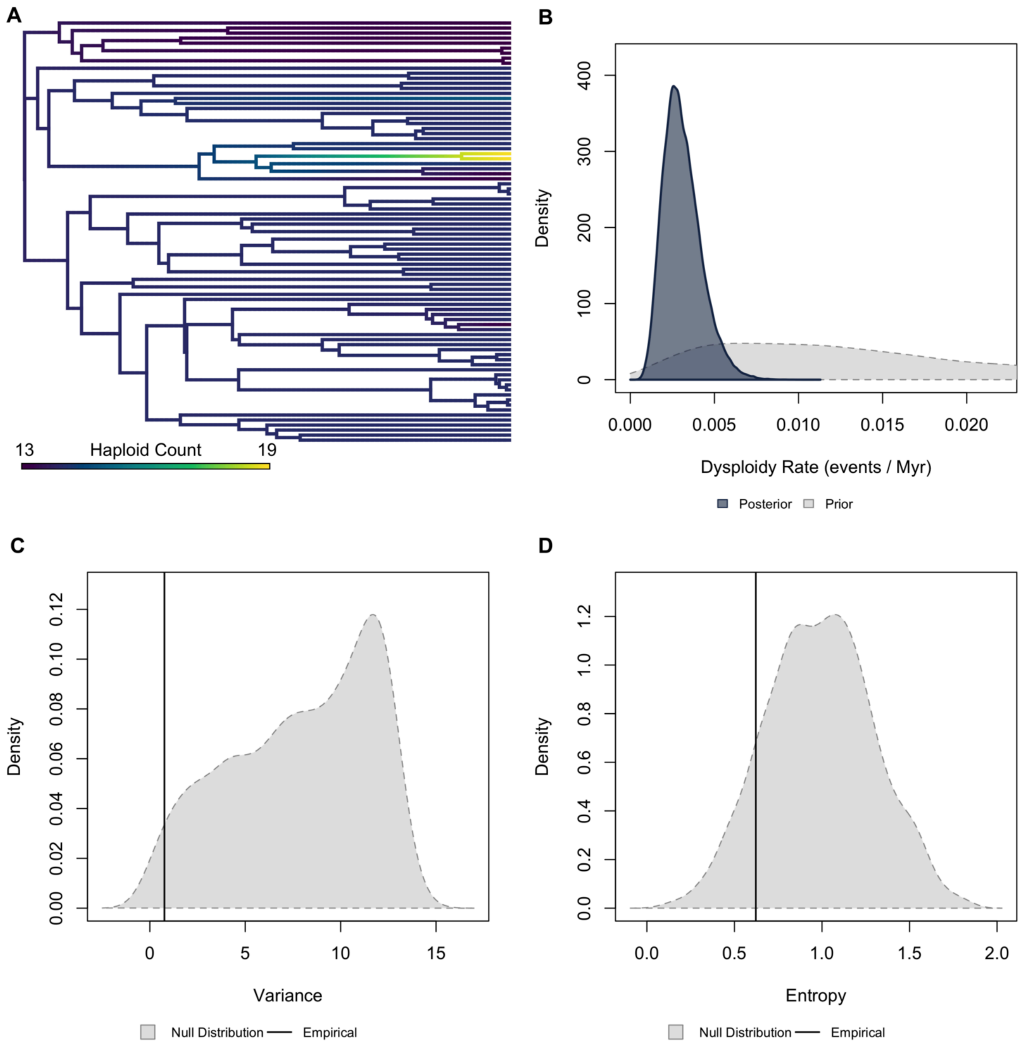


**Figure S33.** ***Validation plots for Odonata.*** *(A) Continuous character map of haploid chromosome number across the phylogeny. (B) Prior versus posterior density distributions for dysploidy rate. (C) Posterior predictive simulation adequacy test for variance. (D) Posterior predictive simulation adequacy test for Shannon's entropy.*

Araneae | Higher Taxonomy: Arachnida | Chromosomes sampled: 1,298 | Phylogenetic tips: 897 | Overlap with phylogeny: 87 species | Root age: 350 Ma | Taxonomic tips: FALSE | Unresolved: N/A

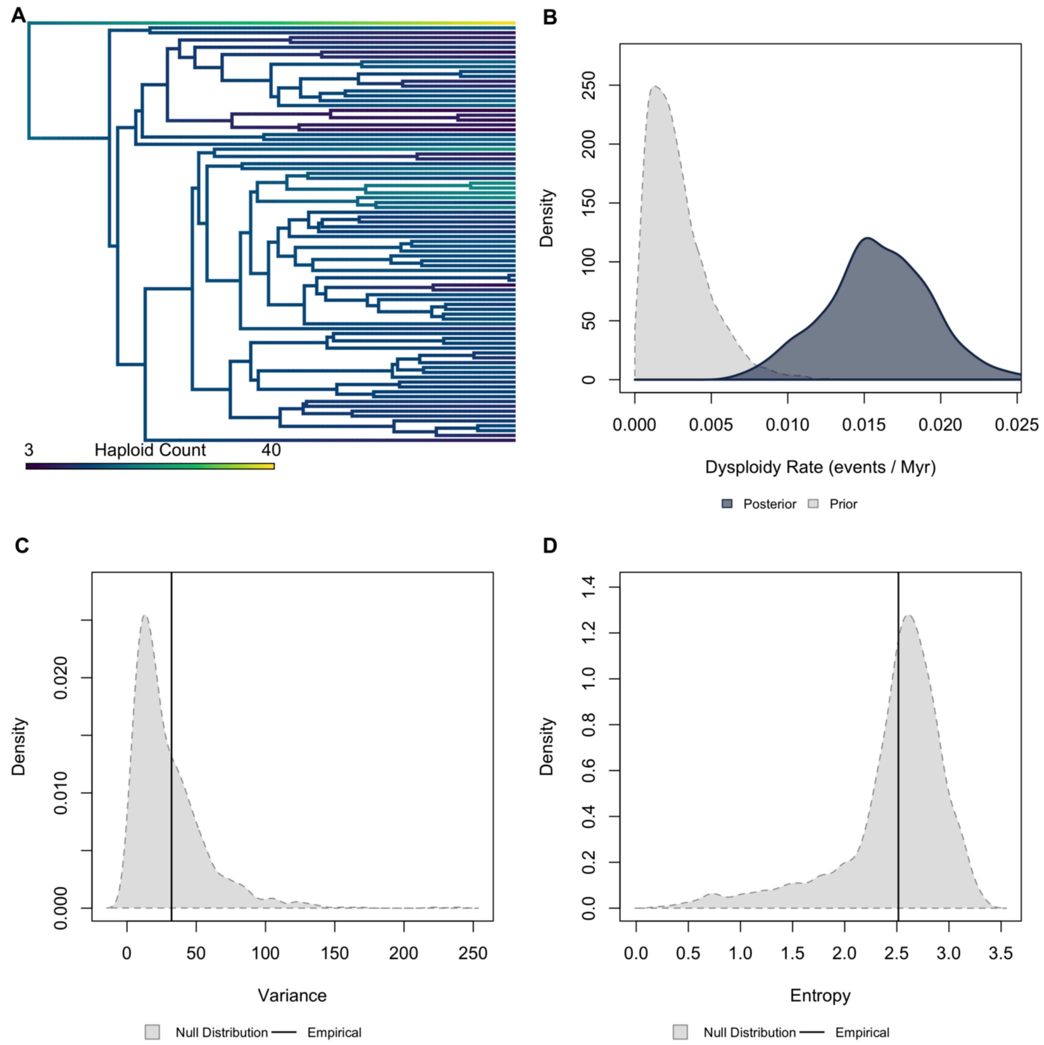


**Figure S34.** ***Validation plots for Araneae.*** *(A) Continuous character map of haploid chromosome number across the phylogeny. (B) Prior versus posterior density distributions for dysploidy rate. (C) Posterior predictive simulation adequacy test for variance. (D) Posterior predictive simulation adequacy test for Shannon's entropy.*

Scorpiones | Higher Taxonomy: Arachnida | Chromosomes sampled: 320 | Phylogenetic tips: 189 | Overlap with phylogeny: 36 species | Root age: 239 Ma | Taxonomic tips: FALSE | Unresolved: N/A

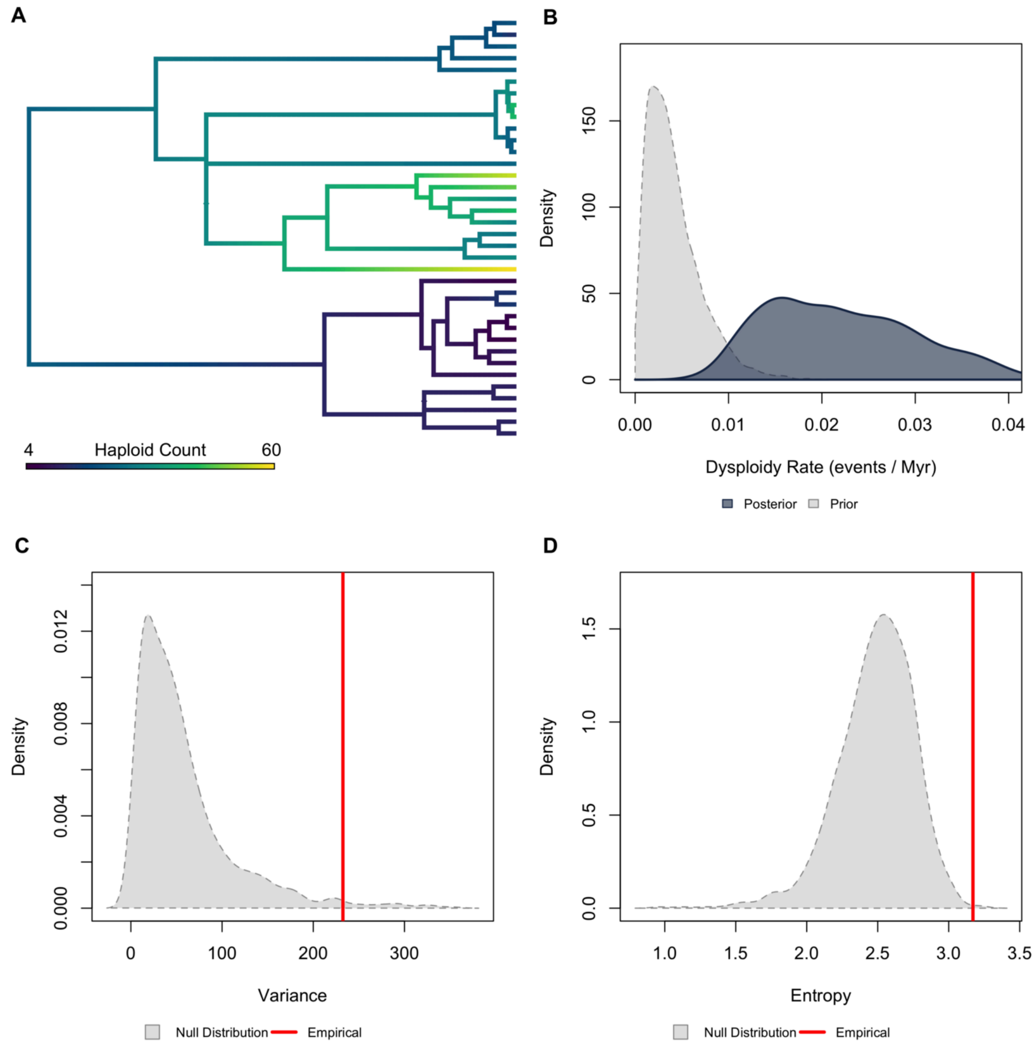


**Figure S35.** ***Validation plots for Scorpiones.*** *(A) Continuous character map of haploid chromosome number across the phylogeny. (B) Prior versus posterior density distributions for dysploidy rate. (C) Posterior predictive simulation adequacy test for variance. (D) Posterior predictive simulation adequacy test for Shannon's entropy.*

Carnivora | Higher Taxonomy: Mammalia | Chromosomes sampled: 167 | Phylogenetic tips: 294 | Overlap with phylogeny: 116 species | Root age: 94 Ma | Taxonomic tips: TRUE | Unresolved: 5.17%

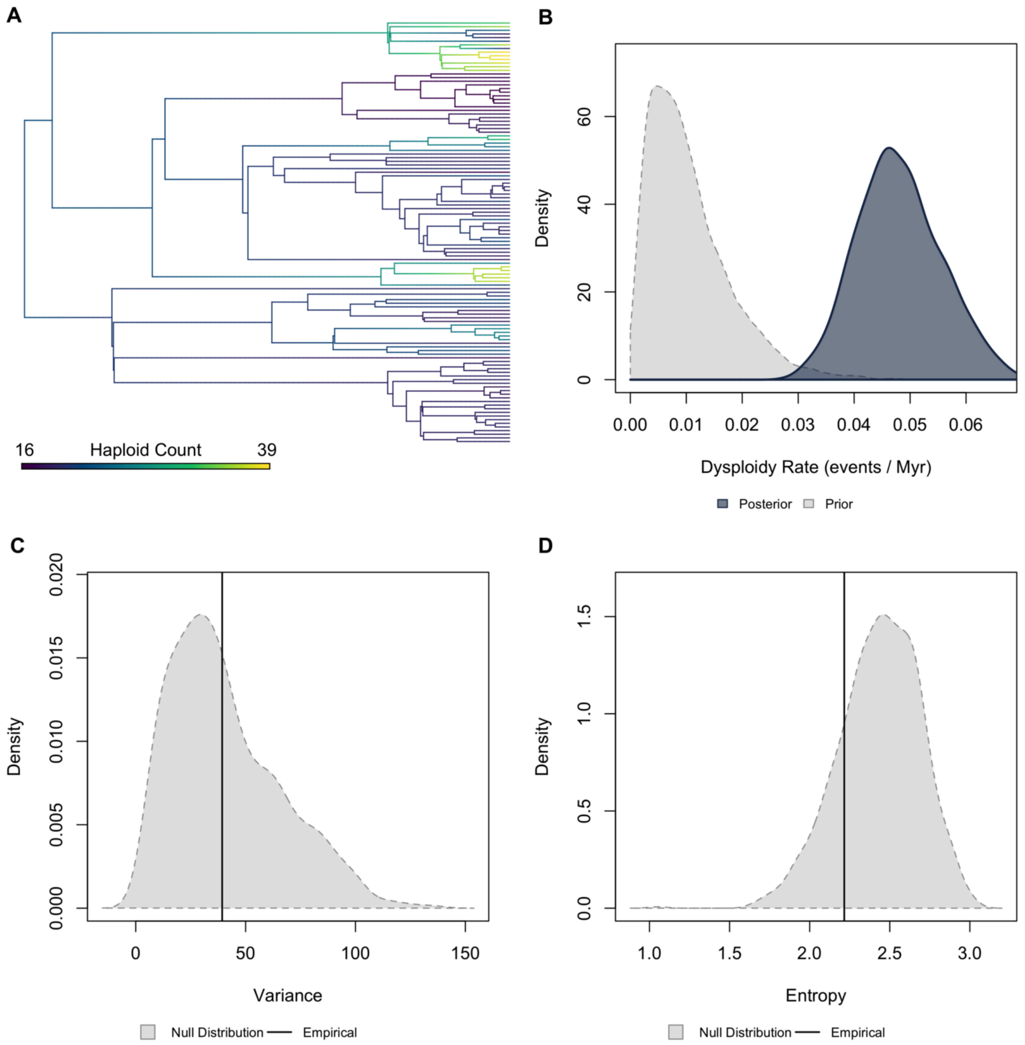


**Figure S36.** ***Validation plots for Carnivora.*** *(A) Continuous character map of haploid chromosome number across the phylogeny. (B) Prior versus posterior density distributions for dysploidy rate. (C) Posterior predictive simulation adequacy test for variance. (D) Posterior predictive simulation adequacy test for Shannon's entropy.*

Cetacea | Higher Taxonomy: Mammalia | Chromosomes sampled: 40 | Phylogenetic tips: 90 | Overlap with phylogeny: 34 species | Root age: 34 Ma | Taxonomic tips: FALSE | Unresolved: N/A

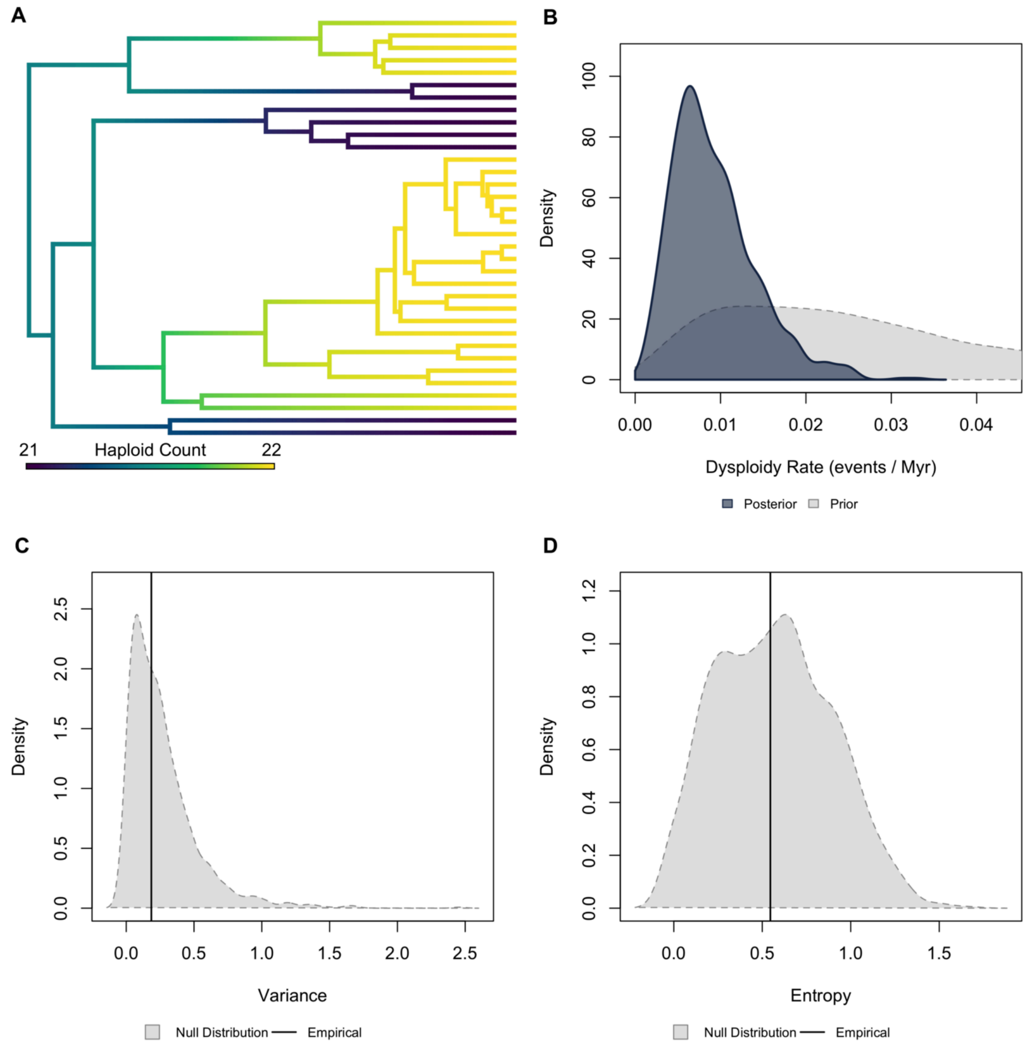


**Figure S37.** ***Validation plots for Cetacea.*** *(A) Continuous character map of haploid chromosome number across the phylogeny. (B) Prior versus posterior density distributions for dysploidy rate. (C) Posterior predictive simulation adequacy test for variance. (D) Posterior predictive simulation adequacy test for Shannon's entropy.*

Chiroptera | Higher Taxonomy: Mammalia | Chromosomes sampled: 204 | Phylogenetic tips: 815 | Overlap with phylogeny: 154 species | Root age: 62 Ma | Taxonomic tips: FALSE | Unresolved: N/A

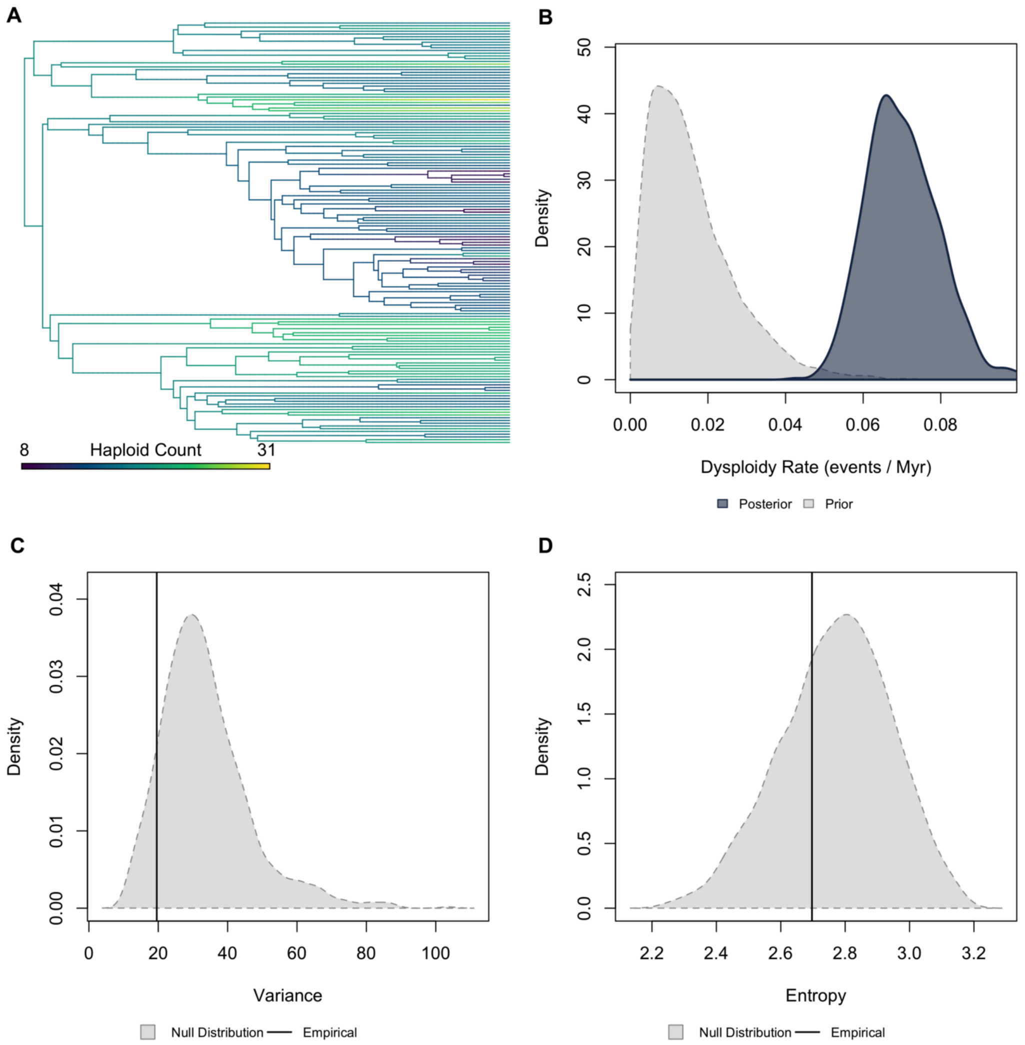


**Figure S38. *Validation plots for Chiroptera.*** *(A) Continuous character map of haploid chromosome number across the phylogeny. (B) Prior versus posterior density distributions for dysploidy rate. (C) Posterior predictive simulation adequacy test for variance. (D) Posterior predictive simulation adequacy test for Shannon's entropy.*

Cricetidae | Higher Taxonomy: Mammalia | Chromosomes sampled: 208 | Phylogenetic tips: 913 | Overlap with phylogeny: 103 species | Root age: 31 Ma | Taxonomic tips: FALSE | Unresolved: N/A

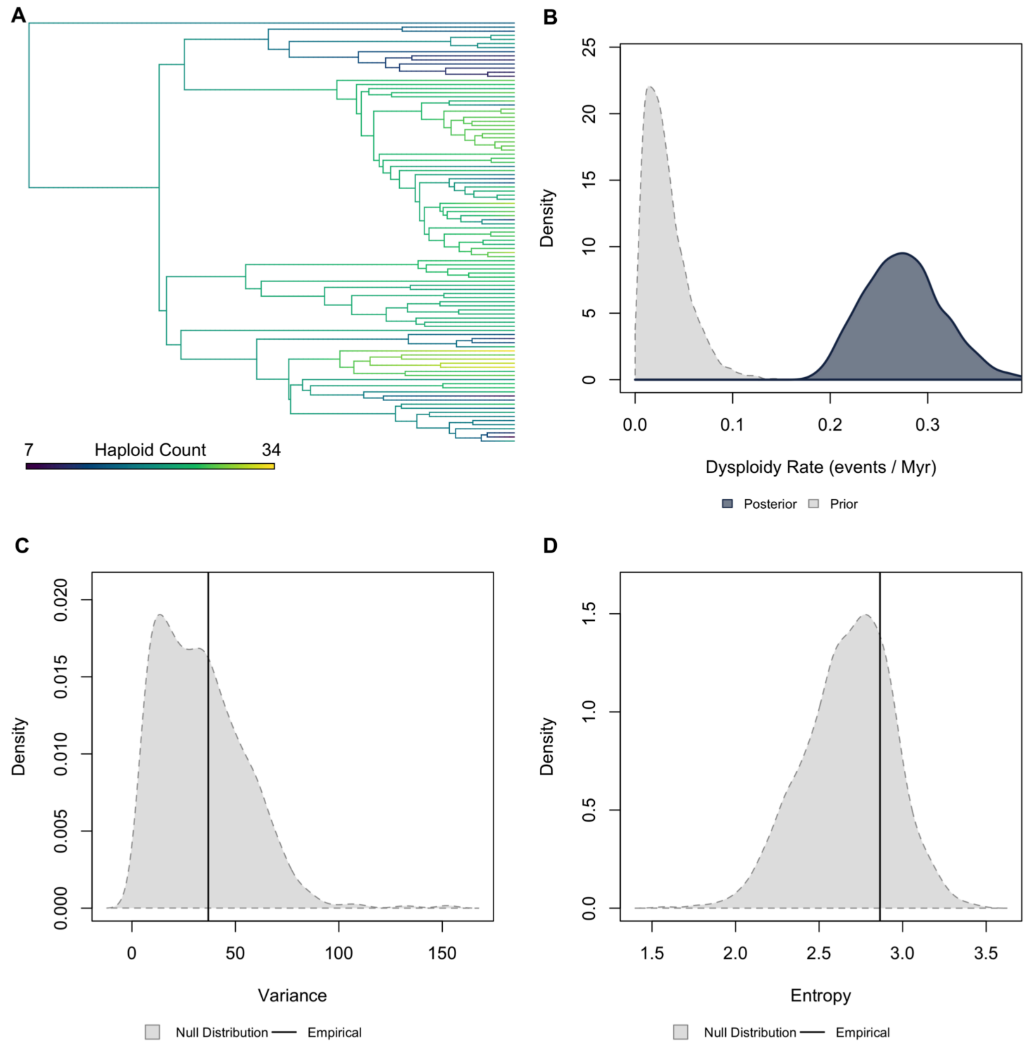


**Figure S39.** ***Validation plots for Cricetidae.*** *(A) Continuous character map of haploid chromosome number across the phylogeny. (B) Prior versus posterior density distributions for dysploidy rate. (C) Posterior predictive simulation adequacy test for variance. (D) Posterior predictive simulation adequacy test for Shannon's entropy.*

Muridae | Higher Taxonomy: Mammalia | Chromosomes sampled: 103 | Phylogenetic tips: 913 | Overlap with phylogeny: 57 species | Root age: 24.5 Ma | Taxonomic tips: FALSE | Unresolved: N/A

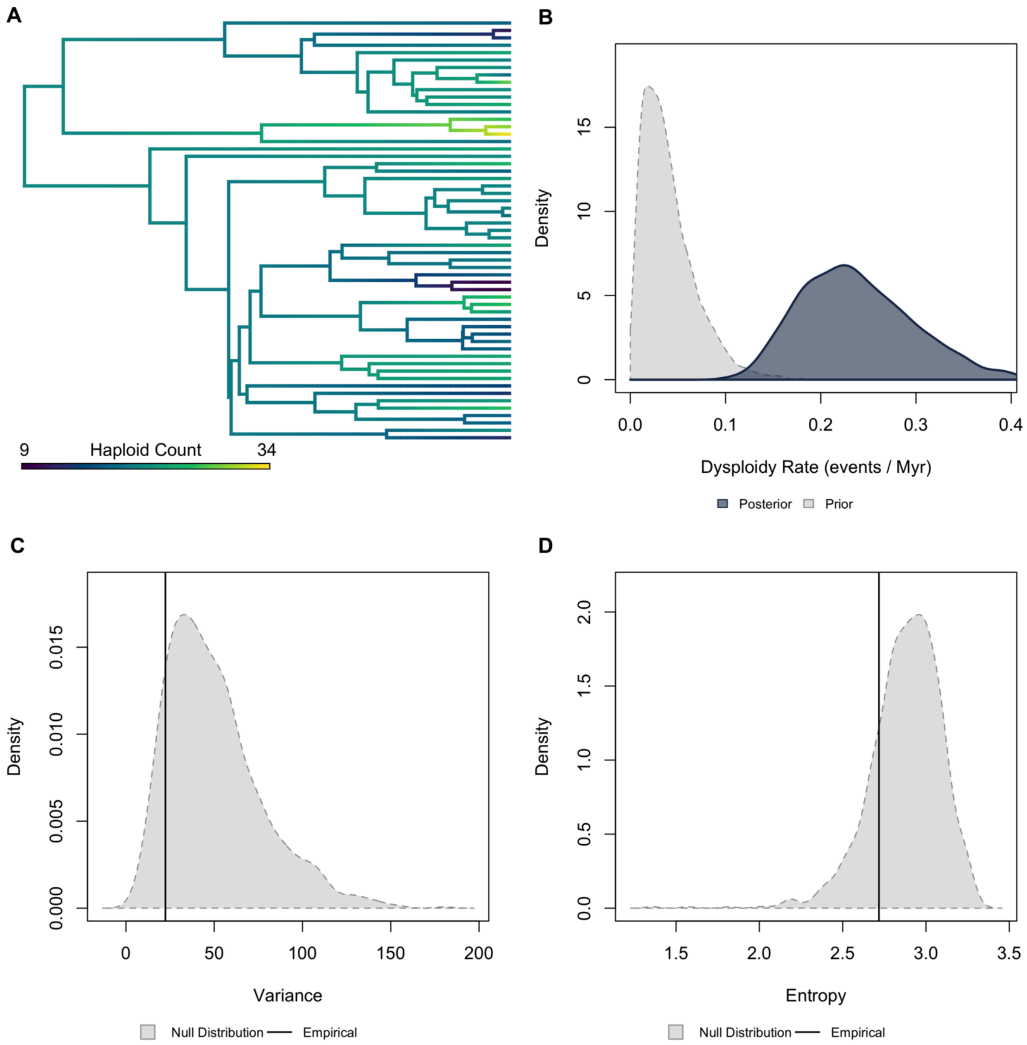


**Figure S40.** ***Validation plots for Muridae.*** *(A) Continuous character map of haploid chromosome number across the phylogeny. (B) Prior versus posterior density distributions for dysploidy rate. (C) Posterior predictive simulation adequacy test for variance. (D) Posterior predictive simulation adequacy test for Shannon's entropy.*

Primates | Higher Taxonomy: Mammalia | Chromosomes sampled: 148 | Phylogenetic tips: 455 | Overlap with phylogeny: 90 species | Root age: 74 Ma | Taxonomic tips: FALSE | Unresolved: N/A

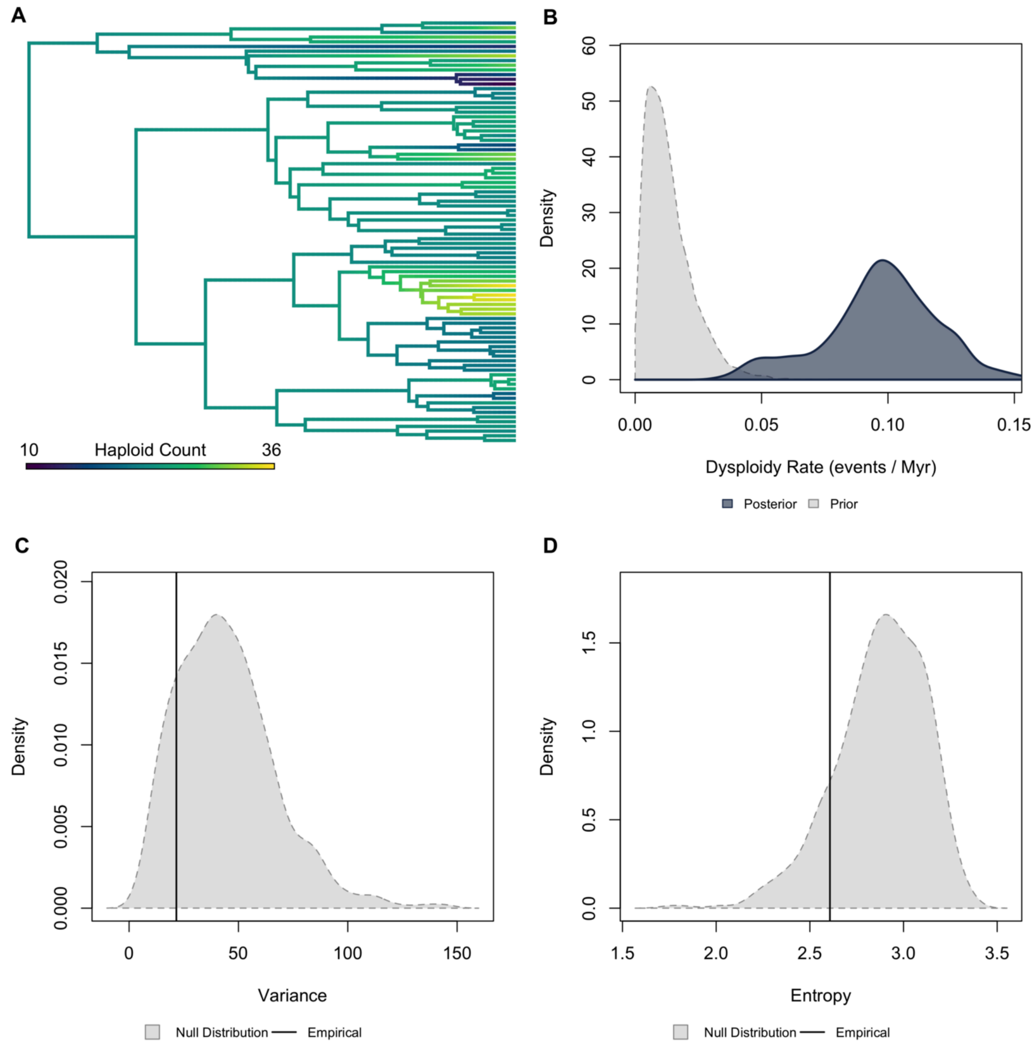


**Figure S41.** ***Validation plots for Primates.*** *(A) Continuous character map of haploid chromosome number across the phylogeny. (B) Prior versus posterior density distributions for dysploidy rate. (C) Posterior predictive simulation adequacy test for variance. (D) Posterior predictive simulation adequacy test for Shannon's entropy.*

Marsupialia | Higher Taxonomy: Mammalia | Chromosomes sampled: 40 | Phylogenetic tips: 40 | Overlap with phylogeny: 40 species | Root age: 78 Ma | Taxonomic tips: FALSE | Unresolved: N/A

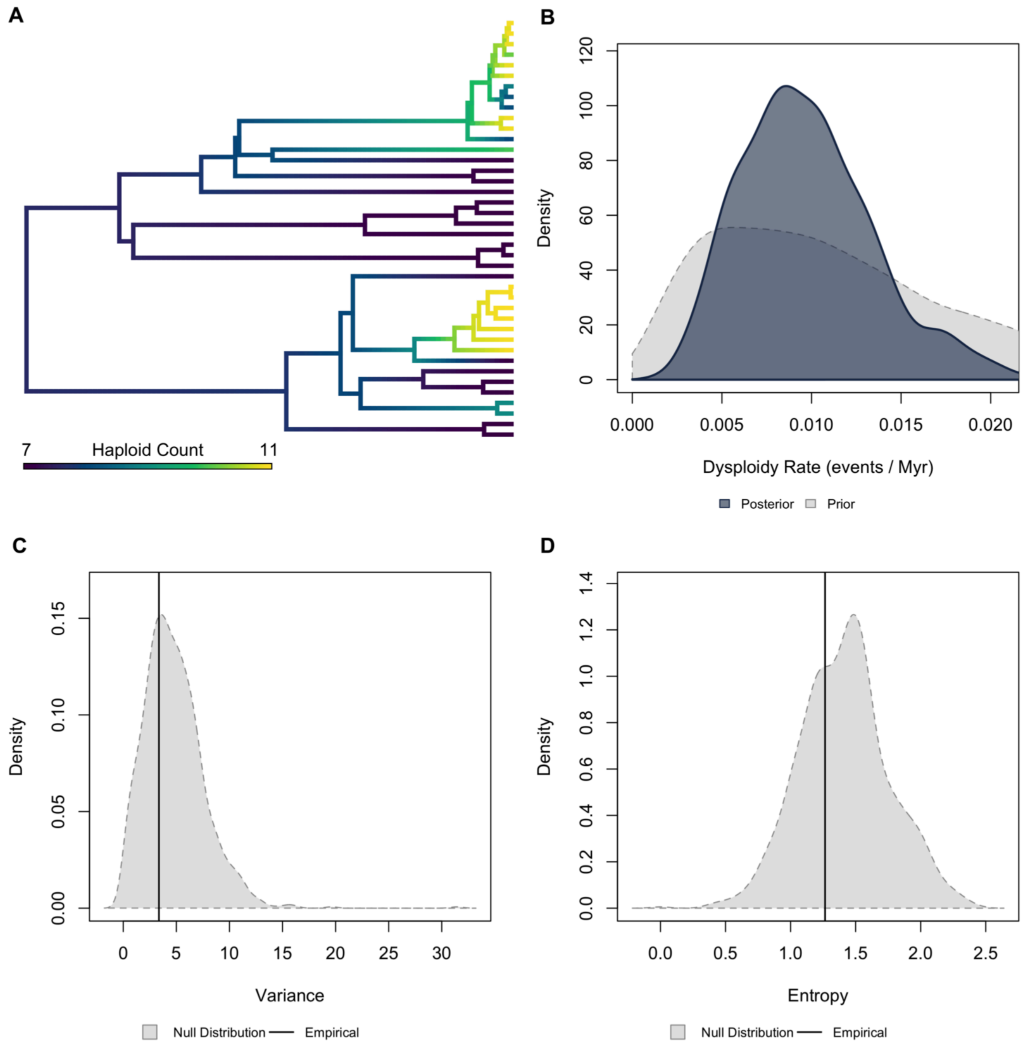


**Figure S42.** ***Validation plots for Marsupialia.*** *(A) Continuous character map of haploid chromosome number across the phylogeny. (B) Prior versus posterior density distributions for dysploidy rate. (C) Posterior predictive simulation adequacy test for variance. (D) Posterior predictive simulation adequacy test for Shannon's entropy.*

Accipitriformes | Higher Taxonomy: Reptilia | Chromosomes sampled: 67 | Phylogenetic tips: 237 | Overlap with phylogeny: 63 species | Root age: 60 Ma | Taxonomic tips: FALSE | Unresolved: N/A

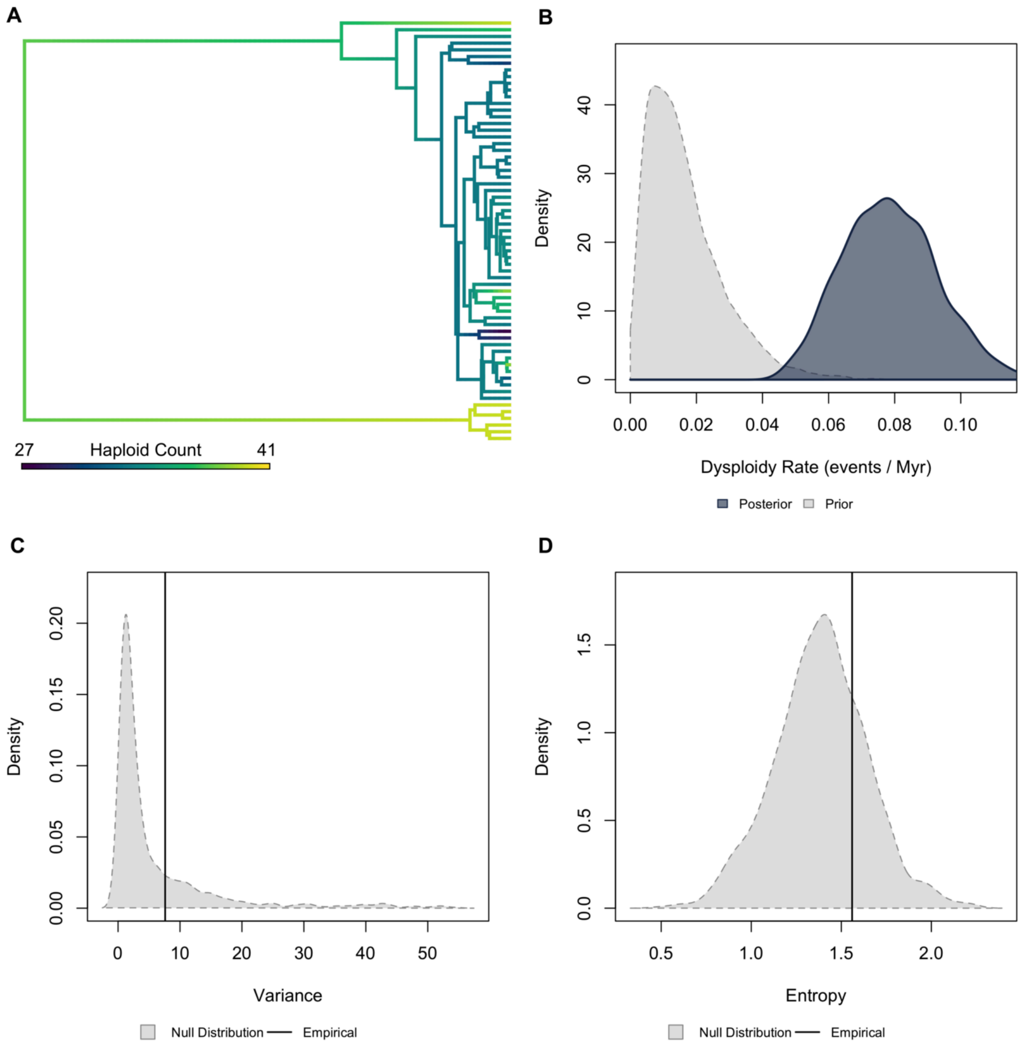


**Figure S43.** ***Validation plots for Accipitriformes.*** *(A) Continuous character map of haploid chromosome number across the phylogeny. (B) Prior versus posterior density distributions for dysploidy rate. (C) Posterior predictive simulation adequacy test for variance. (D) Posterior predictive simulation adequacy test for Shannon's entropy.*

Galliformes | Higher Taxonomy: Reptilia | Chromosomes sampled: 53 | Phylogenetic tips: 52 | Overlap with phylogeny: 52 species | Root age: 67 Ma | Taxonomic tips: FALSE | Unresolved: N/A

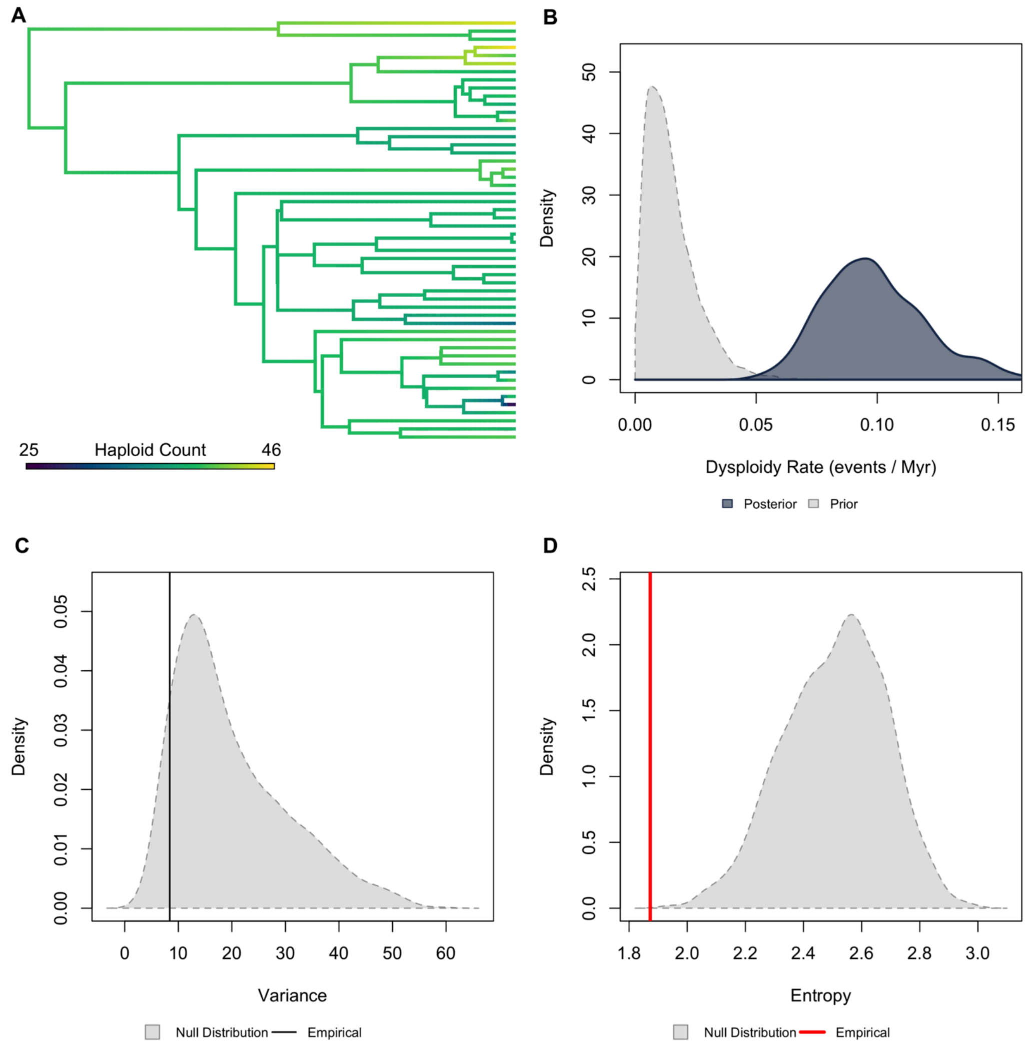


**Figure S44.** ***Validation plots for Galliformes.*** *(A) Continuous character map of haploid chromosome number across the phylogeny. (B) Prior versus posterior density distributions for dysploidy rate. (C) Posterior predictive simulation adequacy test for variance. (D) Posterior predictive simulation adequacy test for Shannon's entropy.*

Gekkonidae | Higher Taxonomy: Reptilia | Chromosomes sampled: 73 | Phylogenetic tips: 1,331 | Overlap with phylogeny: 59 species | Root age: 78 Ma | Taxonomic tips: FALSE | Unresolved: N/A

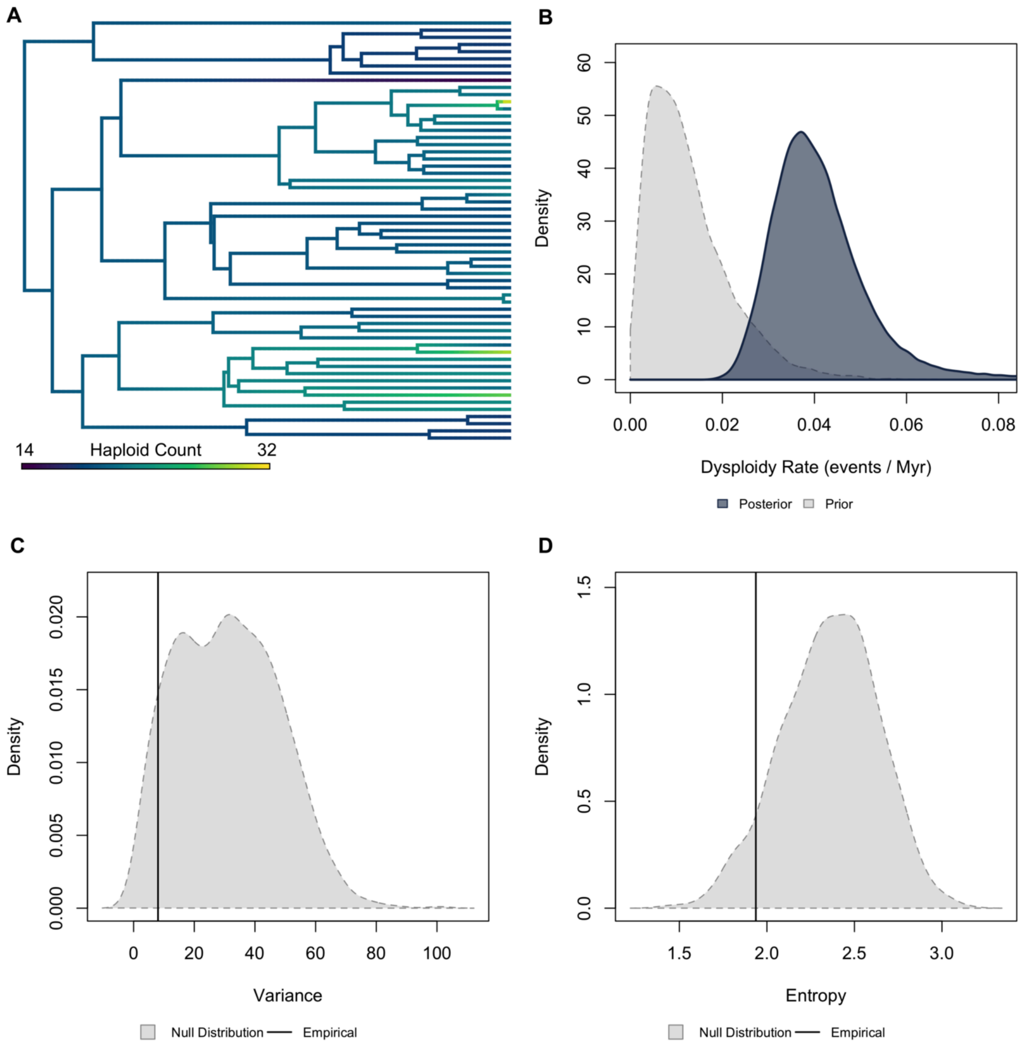


**Figure S45.** ***Validation plots for Gekkonidae.*** *(A) Continuous character map of haploid chromosome number across the phylogeny. (B) Prior versus posterior density distributions for dysploidy rate. (C) Posterior predictive simulation adequacy test for variance. (D) Posterior predictive simulation adequacy test for Shannon's entropy.*

Iguania | Higher Taxonomy: Reptilia | Chromosomes sampled: 382 | Phylogenetic tips: 1,416 | Overlap with phylogeny: 353 species | Root age: 146 Ma | Taxonomic tips: FALSE | Unresolved: N/A

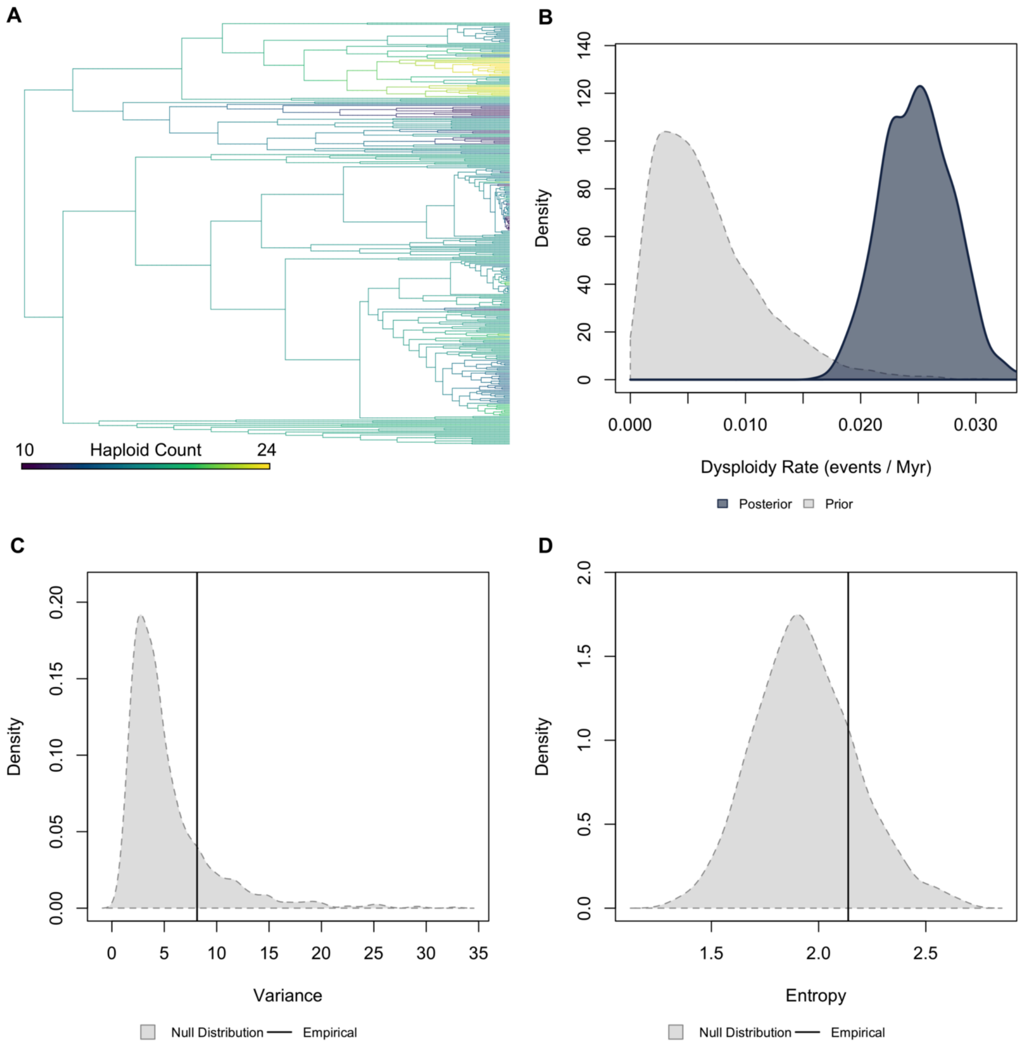


**Figure S46.** ***Validation plots for Iguania.*** *(A) Continuous character map of haploid chromosome number across the phylogeny. (B) Prior versus posterior density distributions for dysploidy rate. (C) Posterior predictive simulation adequacy test for variance. (D) Posterior predictive simulation adequacy test for Shannon's entropy.*

Passeriformes | Higher Taxonomy: Reptilia | Chromosomes sampled: 455 | Phylogenetic tips: 9,189 | Overlap with phylogeny: 449 species | Root age: 64 Ma | Taxonomic tips: FALSE | Unresolved: N/A

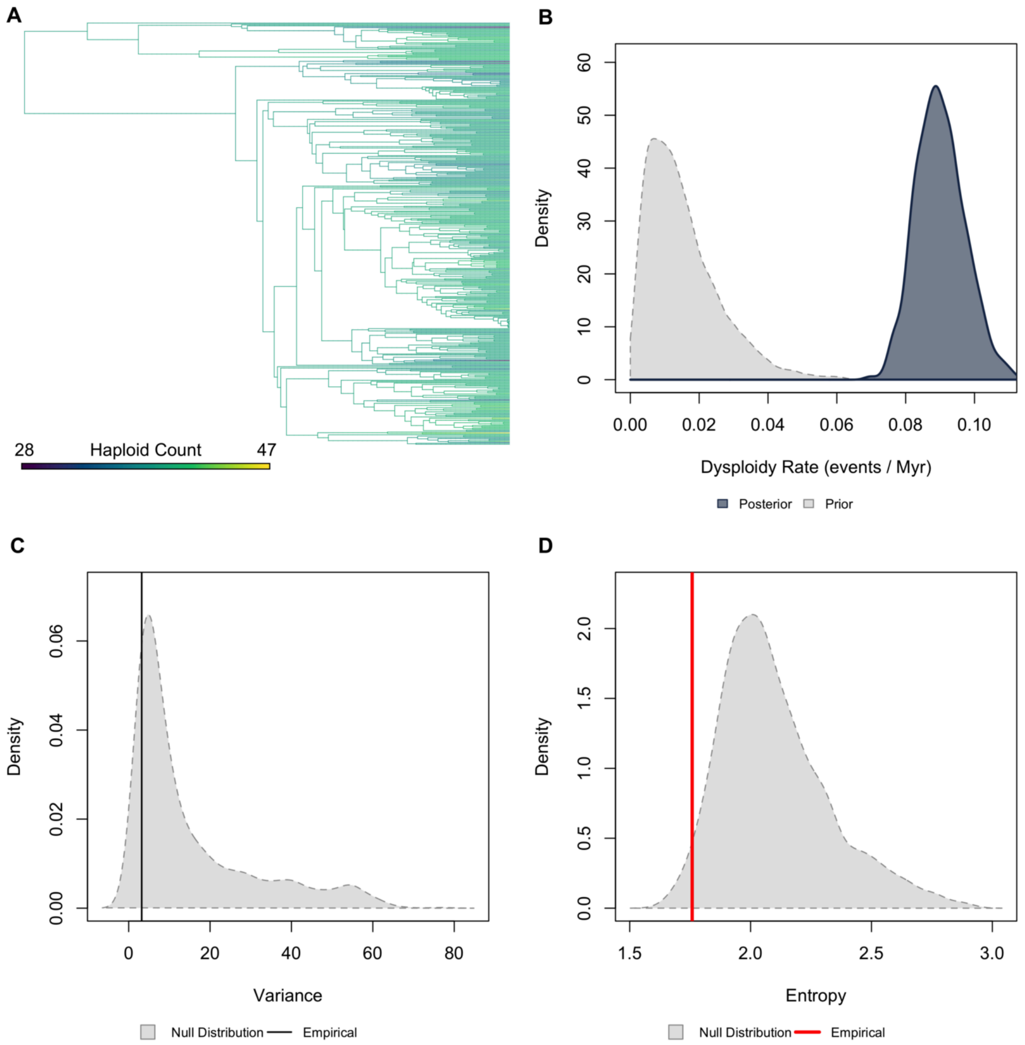


**Figure S47.** ***Validation plots for Passeriformes.*** *(A) Continuous character map of haploid chromosome number across the phylogeny. (B) Prior versus posterior density distributions for dysploidy rate. (C) Posterior predictive simulation adequacy test for variance. (D) Posterior predictive simulation adequacy test for Shannon's entropy.*

Scincoidea | Higher Taxonomy: Reptilia | Chromosomes sampled: 154 | Phylogenetic tips: 1,519 | Overlap with phylogeny: 134 species | Root age: 156 Ma | Taxonomic tips: FALSE | Unresolved: N/A

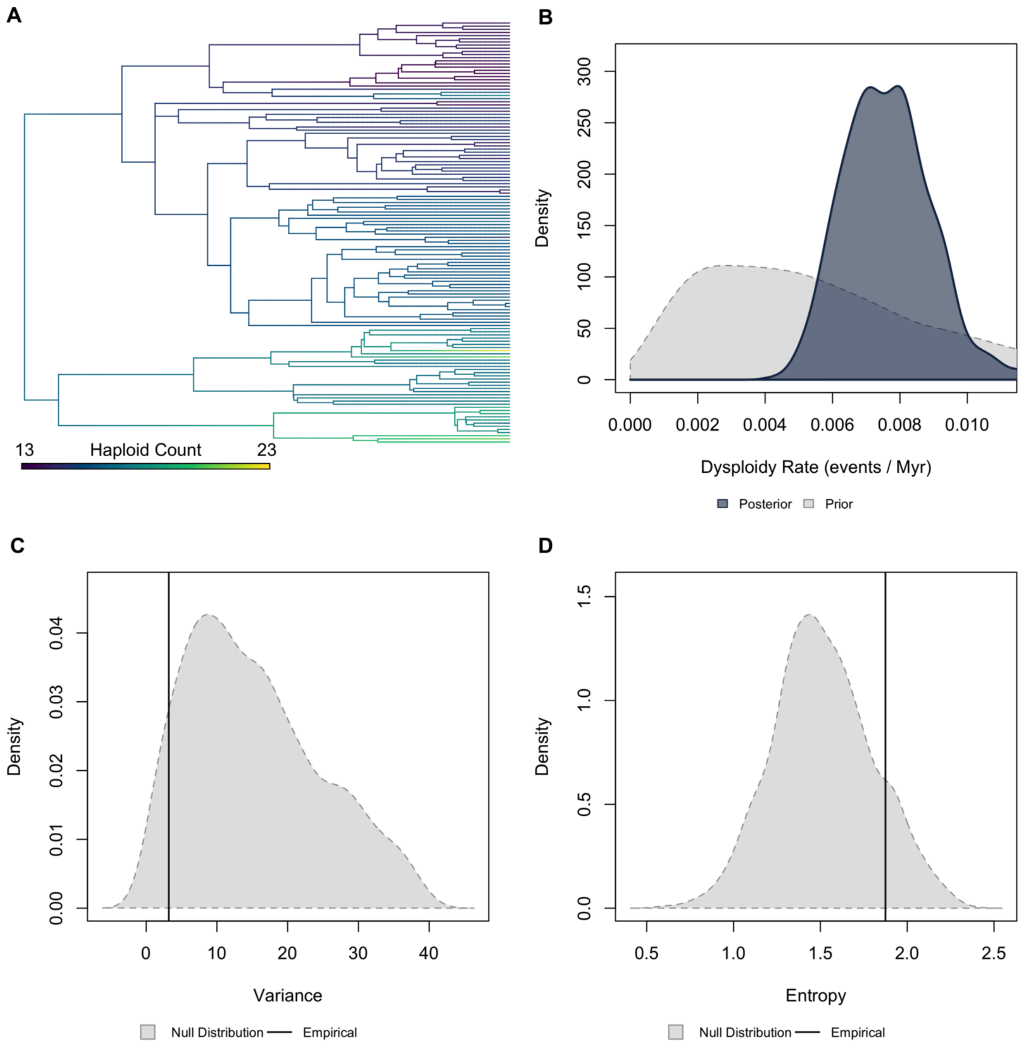


**Figure S48.** ***Validation plots for Scincoidea.*** *(A) Continuous character map of haploid chromosome number across the phylogeny. (B) Prior versus posterior density distributions for dysploidy rate. (C) Posterior predictive simulation adequacy test for variance. (D) Posterior predictive simulation adequacy test for Shannon's entropy.*

Serpentes | Higher Taxonomy: Reptilia | Chromosomes sampled: 256 | Phylogenetic tips: 1,877 | Overlap with phylogeny: 213 species | Root age: 124 Ma | Taxonomic tips: FALSE | Unresolved: N/A

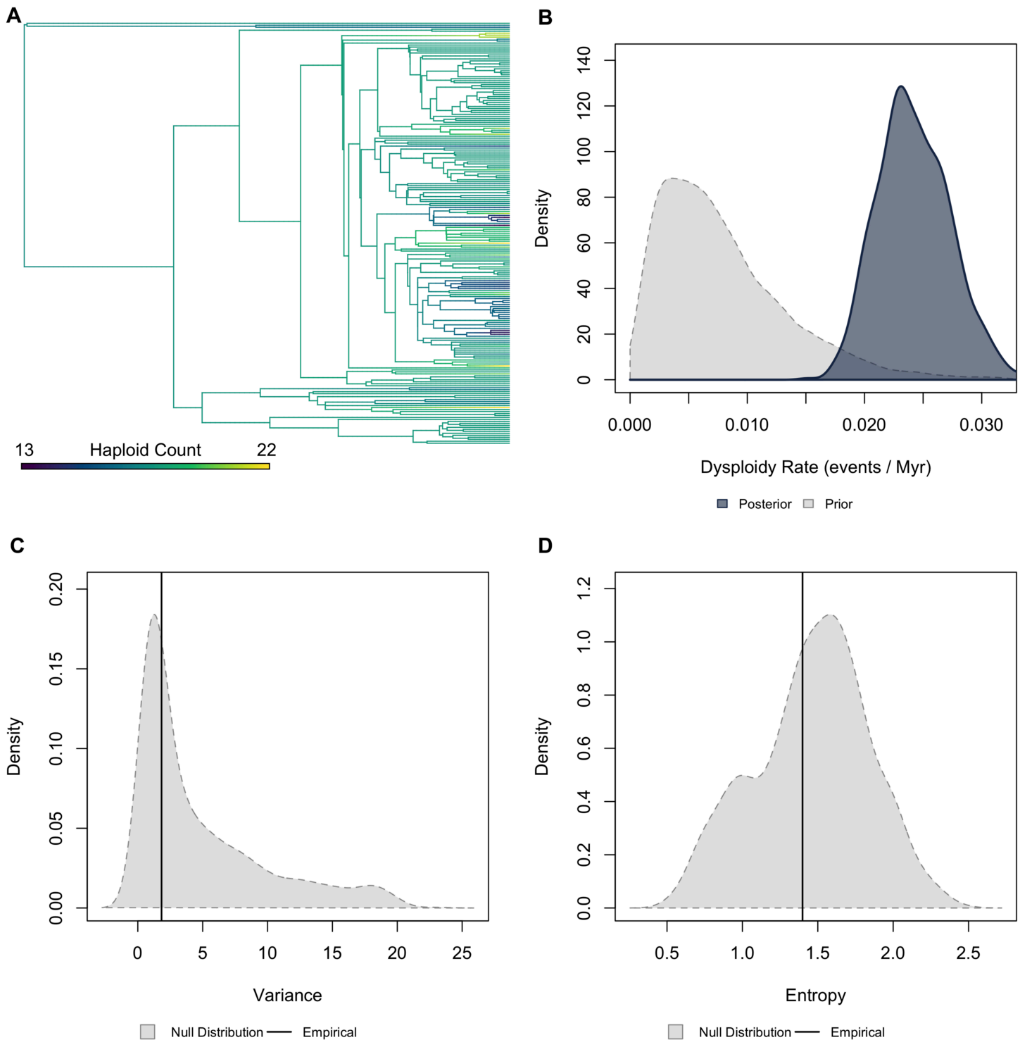


**Figure S49.** ***Validation plots for Serpentes.*** *(A) Continuous character map of haploid chromosome number across the phylogeny. (B) Prior versus posterior density distributions for dysploidy rate. (C) Posterior predictive simulation adequacy test for variance. (D) Posterior predictive simulation adequacy test for Shannon's entropy.*

Testudines | Higher Taxonomy: Reptilia | Chromosomes sampled: 141 | Phylogenetic tips: 286 | Overlap with phylogeny: 122 species | Root age: 201 Ma | Taxonomic tips: TRUE | Unresolved: 3.28%

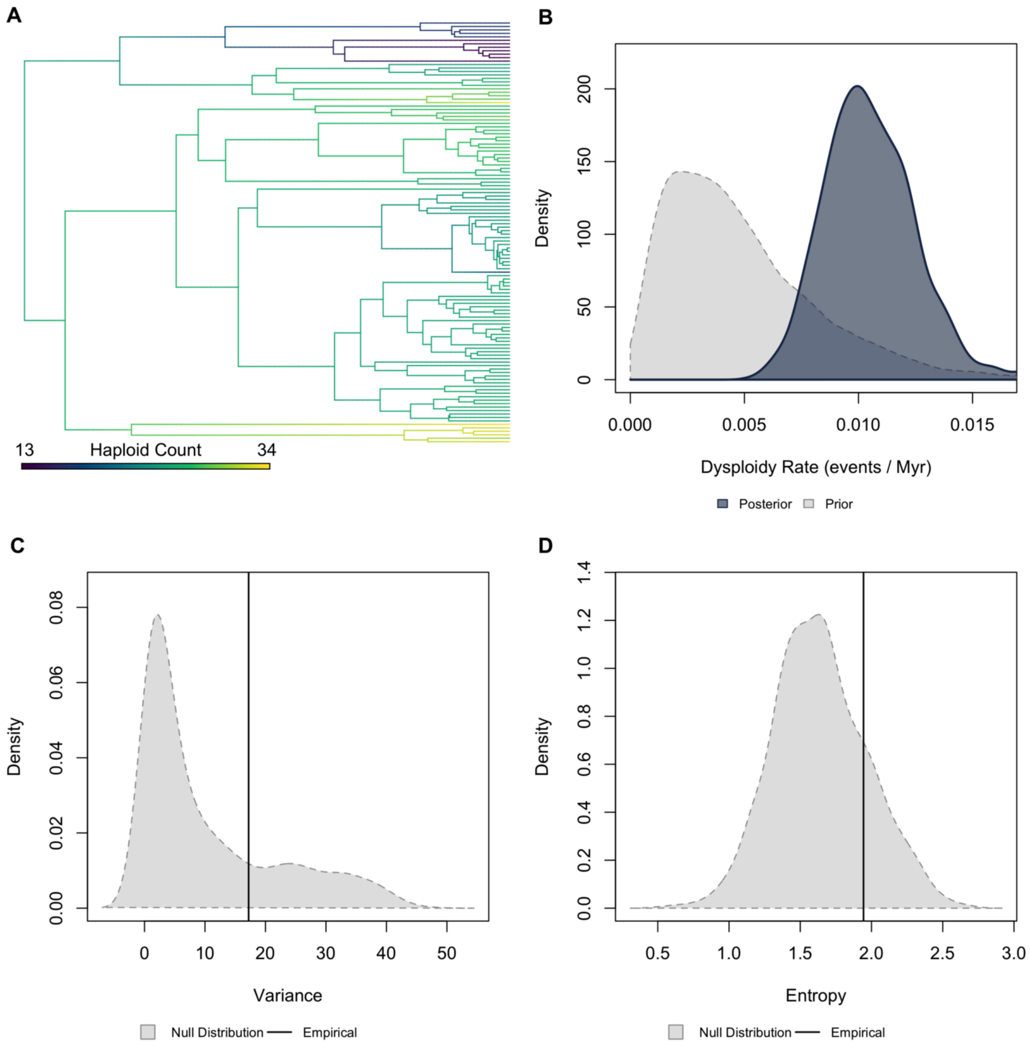


**Figure S50.** ***Validation plots for Testudines.*** *(A) Continuous character map of haploid chromosome number across the phylogeny. (B) Prior versus posterior density distributions for dysploidy rate. (C) Posterior predictive simulation adequacy test for variance. (D) Posterior predictive simulation adequacy test for Shannon's entropy.*

Caudata | Higher Taxonomy: Amphibia | Chromosomes sampled: 246 | Phylogenetic tips: 796 | Overlap with phylogeny: 204 species | Root age: 415 Ma | Taxonomic tips: FALSE | Unresolved: N/A

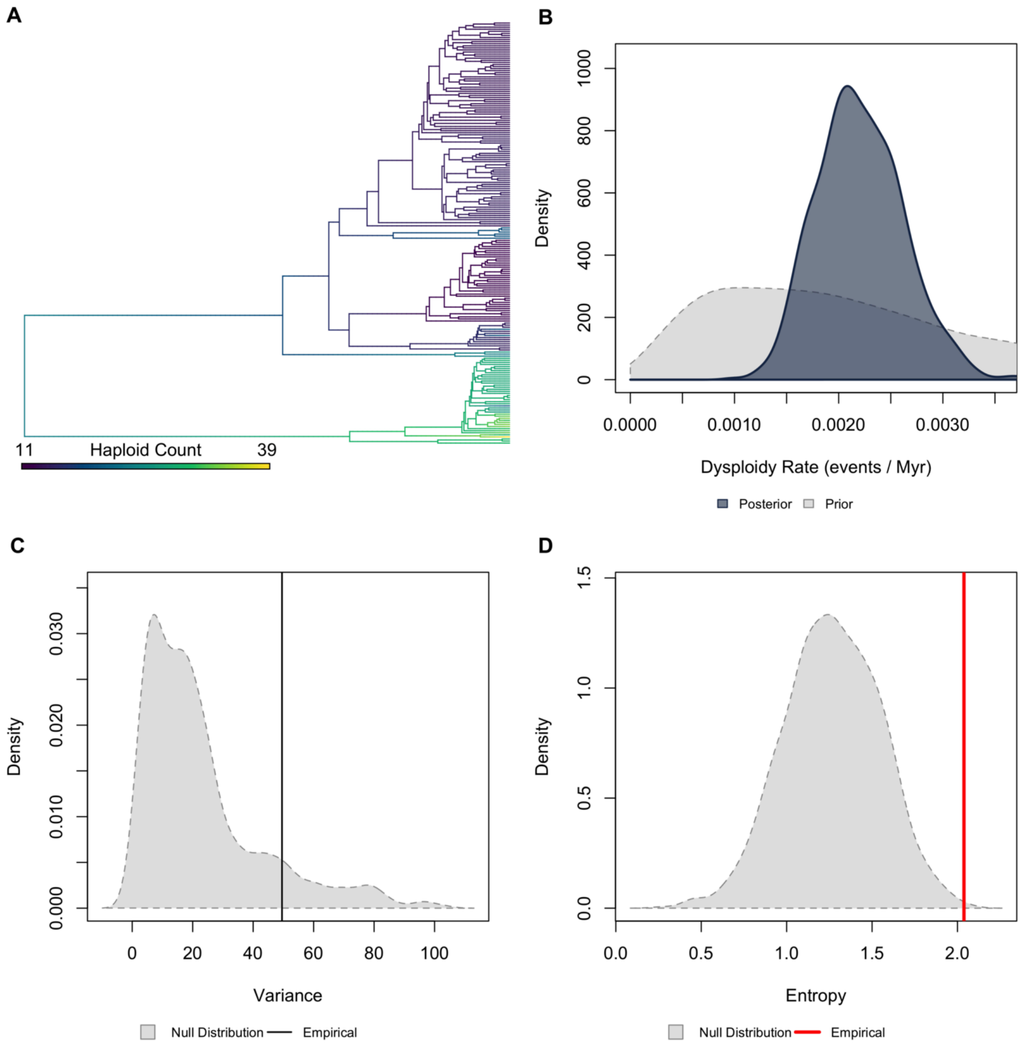


**Figure S51.** ***Validation plots for Caudata.*** *(A) Continuous character map of haploid chromosome number across the phylogeny. (B) Prior versus posterior density distributions for dysploidy rate. (C) Posterior predictive simulation adequacy test for variance. (D) Posterior predictive simulation adequacy test for Shannon's entropy.*

Anura | Higher Taxonomy: Amphibia | Chromosomes sampled: 1,831 | Phylogenetic tips: 5,326 | Overlap with phylogeny: 1,207 species | Root age: 210 Ma | Taxonomic tips: FALSE | Unresolved: N/A

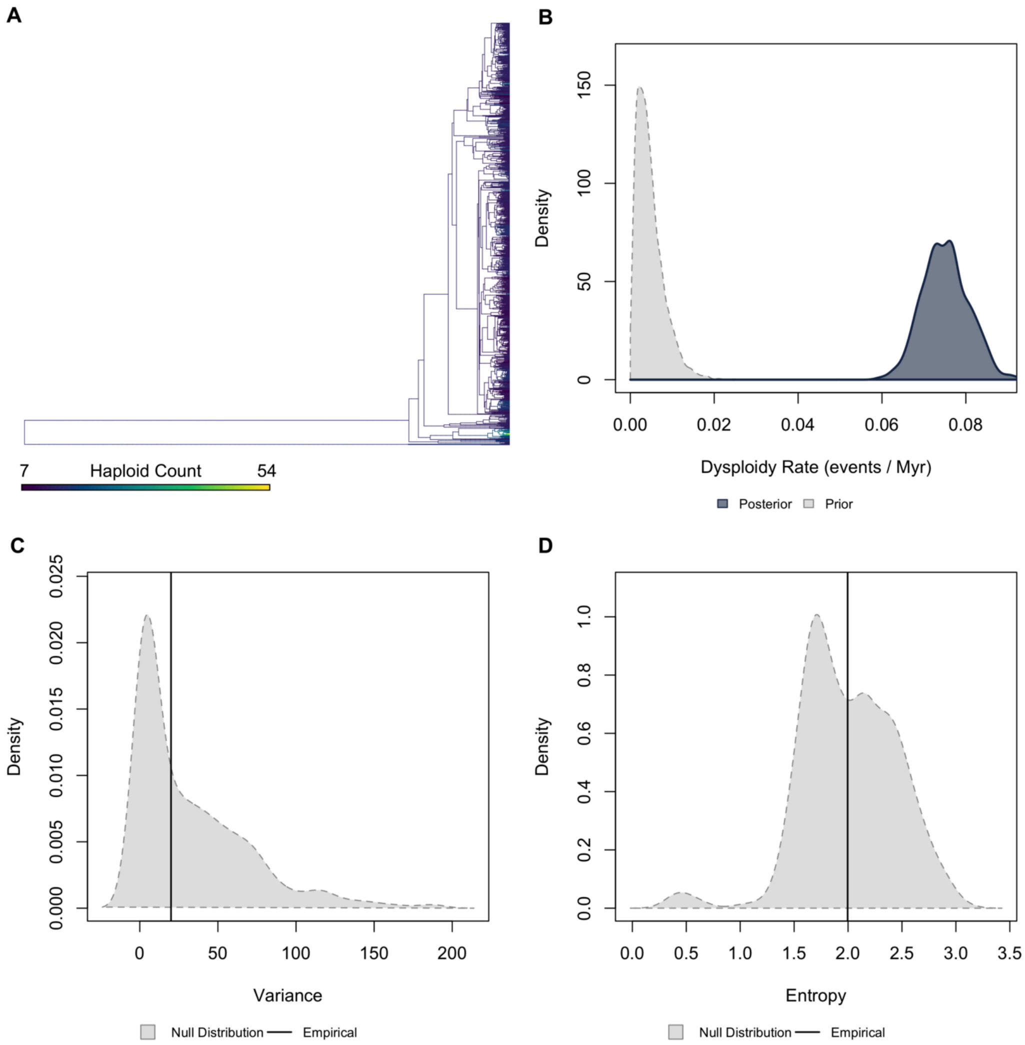


**Figure S52. *Validation plots for Anura.*** *(A) Continuous character map of haploid chromosome number across the phylogeny. (B) Prior versus posterior density distributions for dysploidy rate. (C) Posterior predictive simulation adequacy test for variance. (D) Posterior predictive simulation adequacy test for Shannon's entropy.*

Anabantiformes | Higher Taxonomy: Actinopterygii | Chromosomes sampled: 61 | Phylogenetic tips: 196 | Overlap with phylogeny: 41 species | Root age: 73 Ma | Taxonomic tips: FALSE | Unresolved: N/A

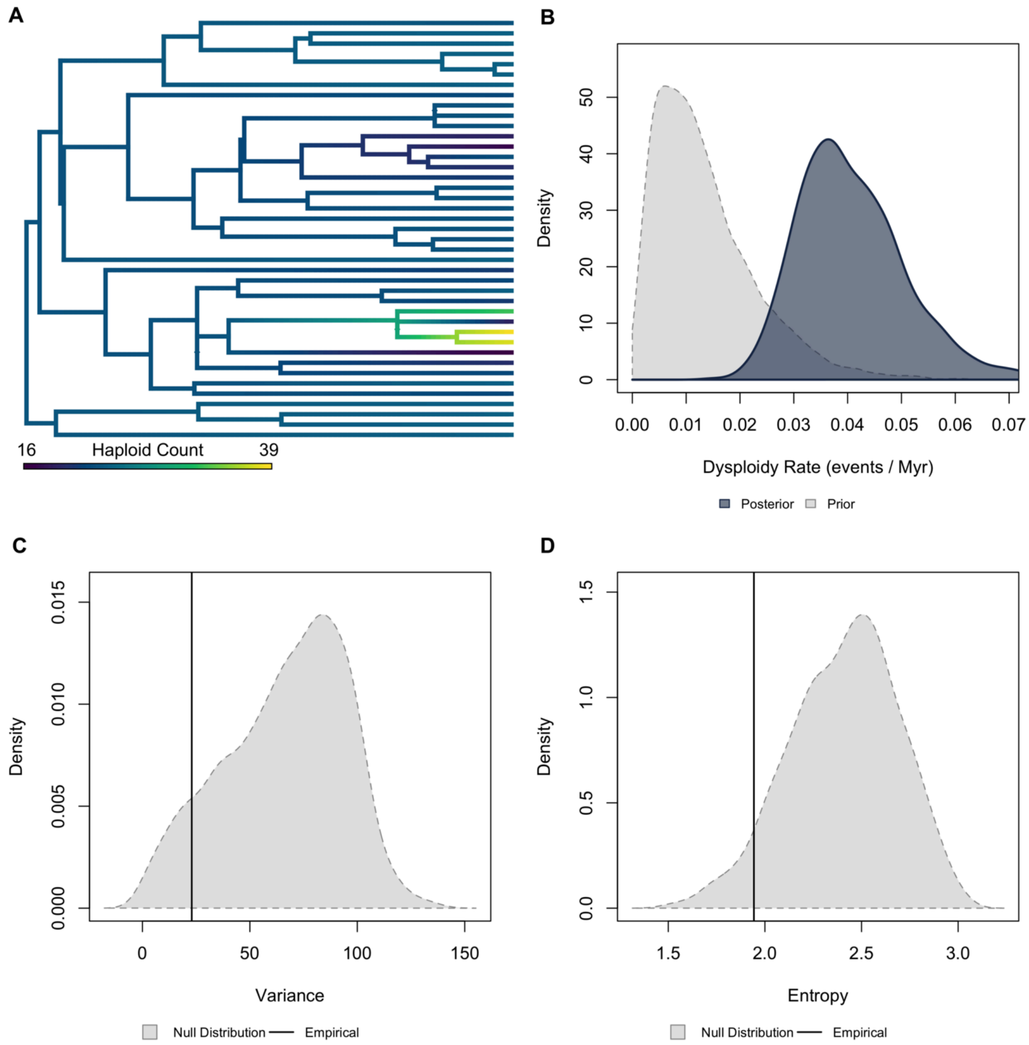


**Figure S53. *Validation plots for Anabantiformes.*** *(A) Continuous character map of haploid chromosome number across the phylogeny. (B) Prior versus posterior density distributions for dysploidy rate. (C) Posterior predictive simulation adequacy test for variance. (D) Posterior predictive simulation adequacy test for Shannon's entropy.*

Cichlidae | Higher Taxonomy: Actinopterygii | Chromosomes sampled: 281 | Phylogenetic tips: 750 | Overlap with phylogeny: 88 species | Root age: 60 Ma | Taxonomic tips: FALSE | Unresolved: N/A

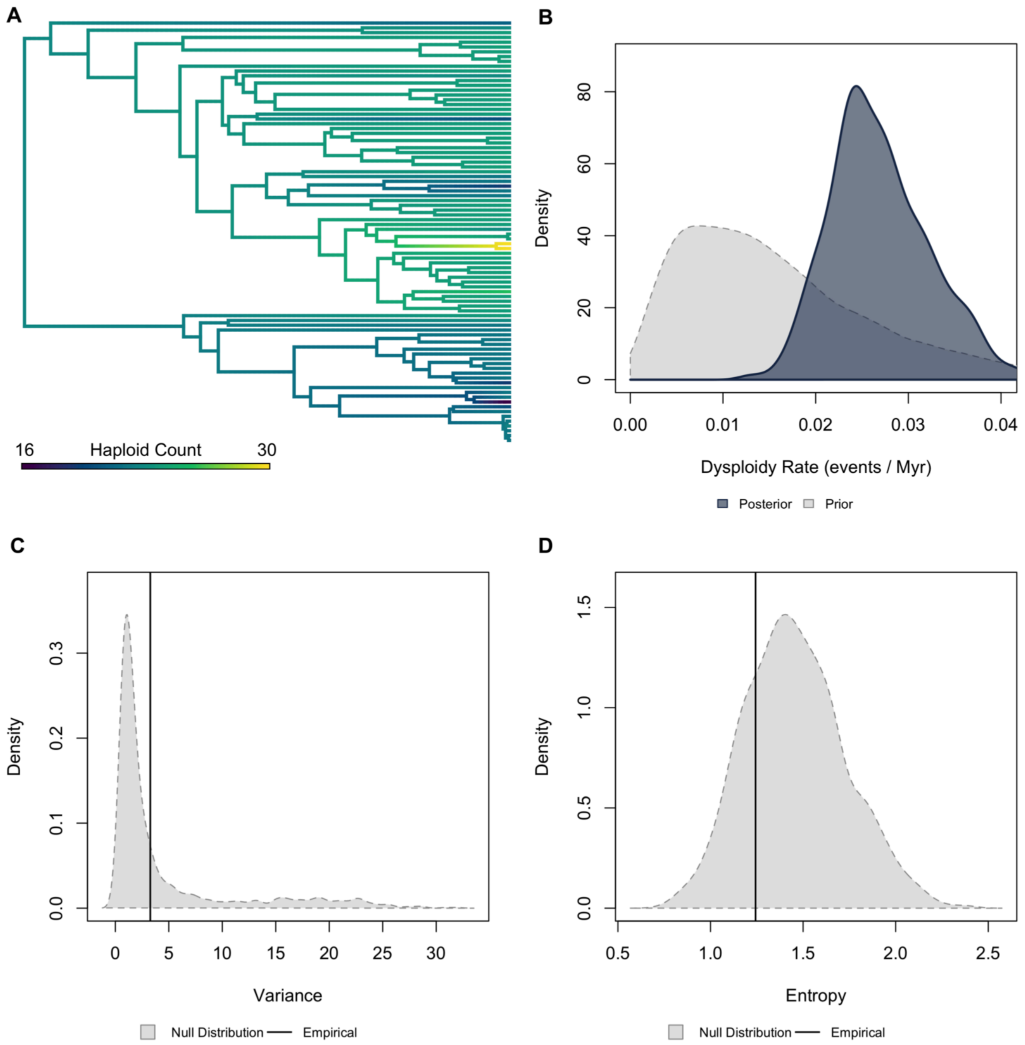


**Figure S54.** ***Validation plots for Cichlidae.*** *(A) Continuous character map of haploid chromosome number across the phylogeny. (B) Prior versus posterior density distributions for dysploidy rate. (C) Posterior predictive simulation adequacy test for variance. (D) Posterior predictive simulation adequacy test for Shannon's entropy.*

Nothobranchiidae | Higher Taxonomy: Actinopterygii | Chromosomes sampled: 242 | Phylogenetic tips: 11,638 | Overlap with phylogeny: 79 species | Root age: 73 Ma | Taxonomic tips: FALSE | Unresolved: N/A

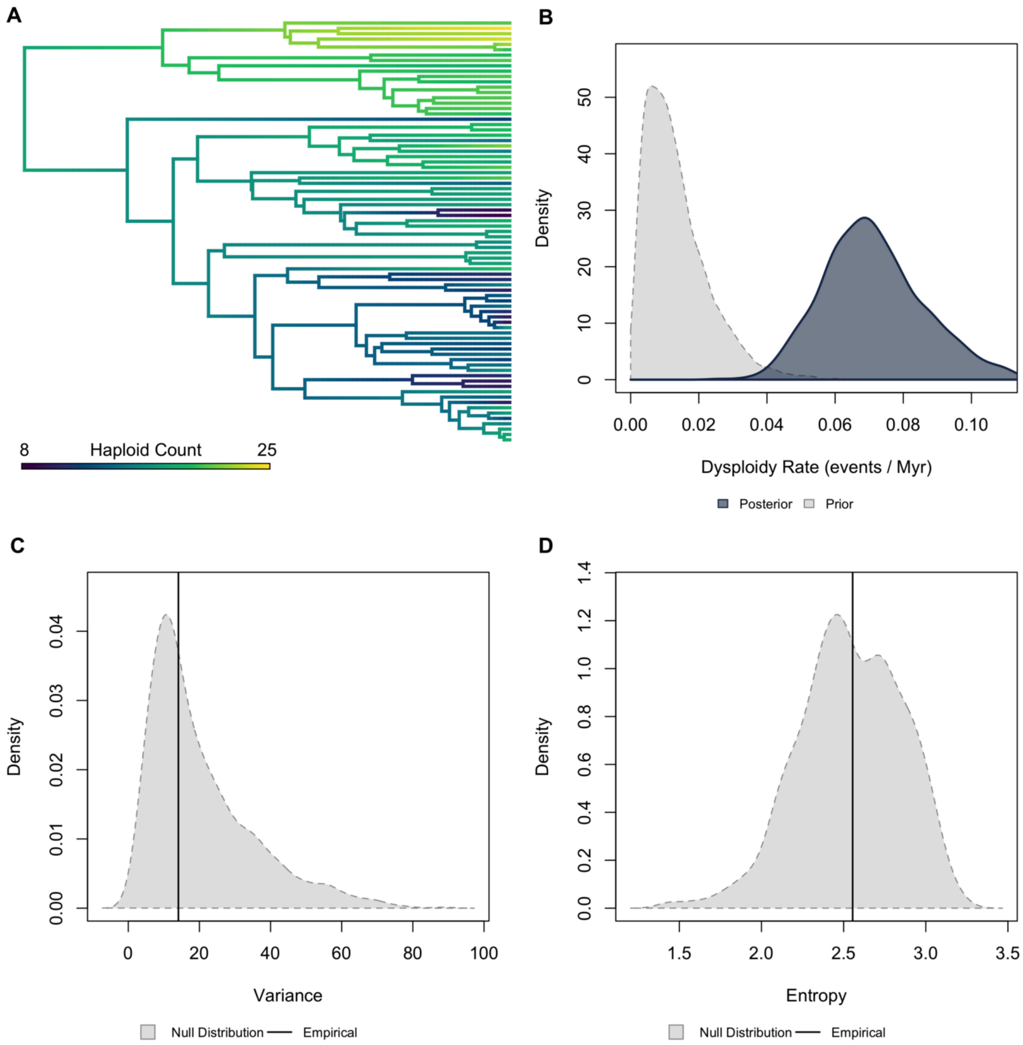


**Figure S55.** ***Validation plots for Nothobranchiidae.*** *(A) Continuous character map of haploid chromosome number across the phylogeny. (B) Prior versus posterior density distributions for dysploidy rate. (C) Posterior predictive simulation adequacy test for variance. (D) Posterior predictive simulation adequacy test for Shannon's entropy.*

Gobiidae | Higher Taxonomy: Actinopterygii | Chromosomes sampled: 232 | Phylogenetic tips: 827 | Overlap with phylogeny: 65 species | Root age: 52 Ma | Taxonomic tips: FALSE | Unresolved: N/A

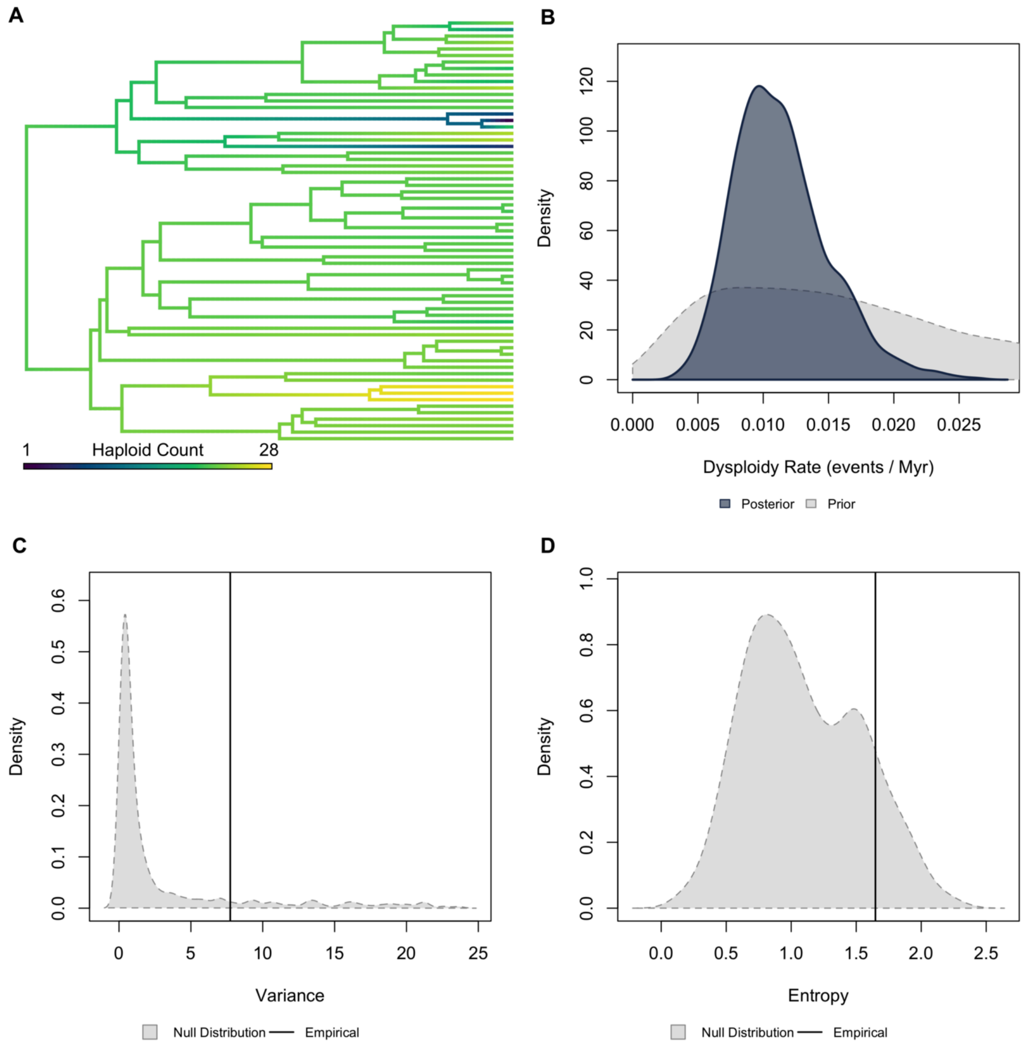


**Figure S56.** ***Validation plots for Gobiidae.*** *(A) Continuous character map of haploid chromosome number across the phylogeny. (B) Prior versus posterior density distributions for dysploidy rate. (C) Posterior predictive simulation adequacy test for variance. (D) Posterior predictive simulation adequacy test for Shannon's entropy.*

Siluriformes | Higher Taxonomy: Actinopterygii | Chromosomes sampled: 165 | Phylogenetic tips: 2,009 | Overlap with phylogeny: 131 species | Root age: 96 Ma | Taxonomic tips: FALSE | Unresolved: N/A

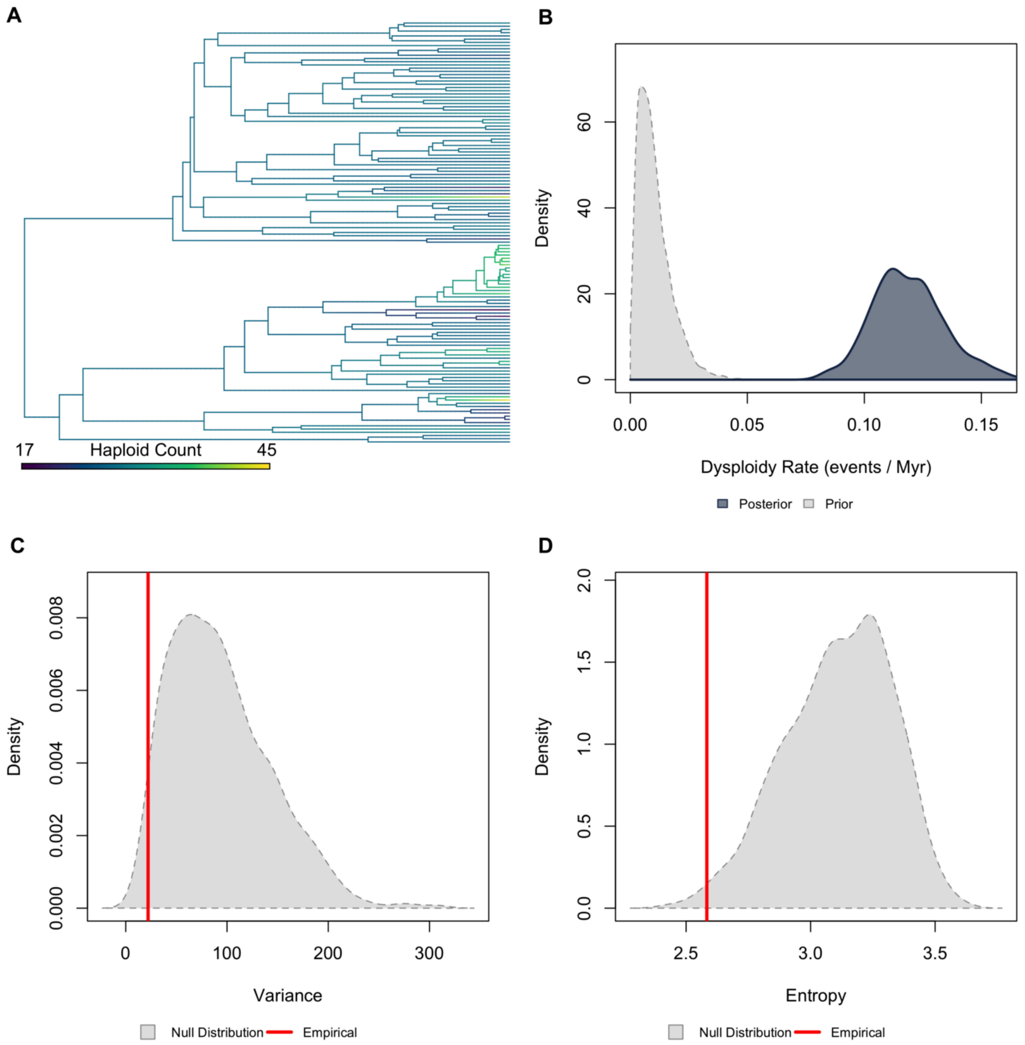


**Figure S57.** ***Validation plots for Siluriformes.*** *(A) Continuous character map of haploid chromosome number across the phylogeny. (B) Prior versus posterior density distributions for dysploidy rate. (C) Posterior predictive simulation adequacy test for variance. (D) Posterior predictive simulation adequacy test for Shannon's entropy.*

Characidae | Higher Taxonomy: Actinopterygii | Chromosomes sampled: 663 | Phylogenetic tips: 1,019 | Overlap with phylogeny: 238 species | Root age: 108 Ma | Taxonomic tips: FALSE | Unresolved: N/A

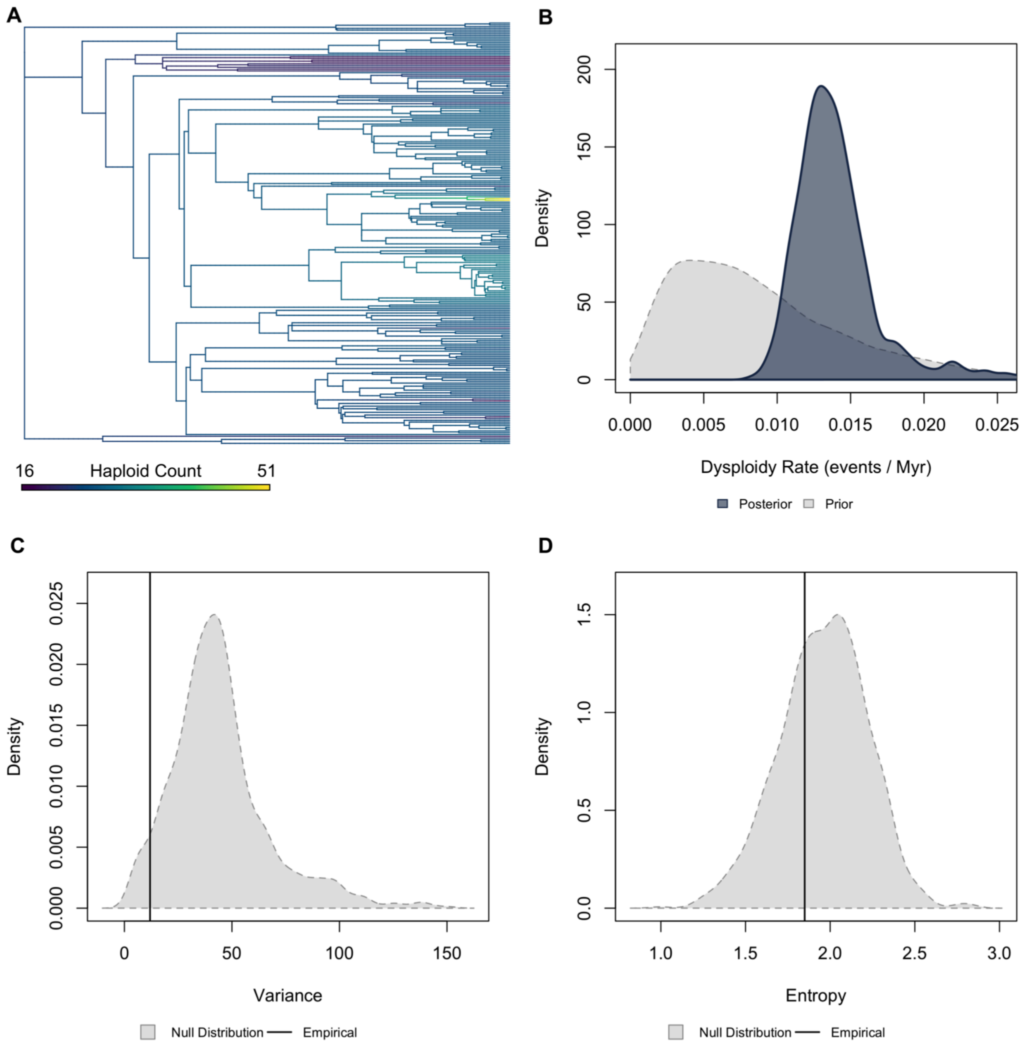


**Figure S58.** ***Validation plots for Characidae.*** *(A) Continuous character map of haploid chromosome number across the phylogeny. (B) Prior versus posterior density distributions for dysploidy rate. (C) Posterior predictive simulation adequacy test for variance. (D) Posterior predictive simulation adequacy test for Shannon's entropy.*

Cyprinidae | Higher Taxonomy: Actinopterygii | Chromosomes sampled: 1,111 | Phylogenetic tips: 1,368 | Overlap with phylogeny: 398 species | Root age: 90 Ma | Taxonomic tips: FALSE | Unresolved: N/A

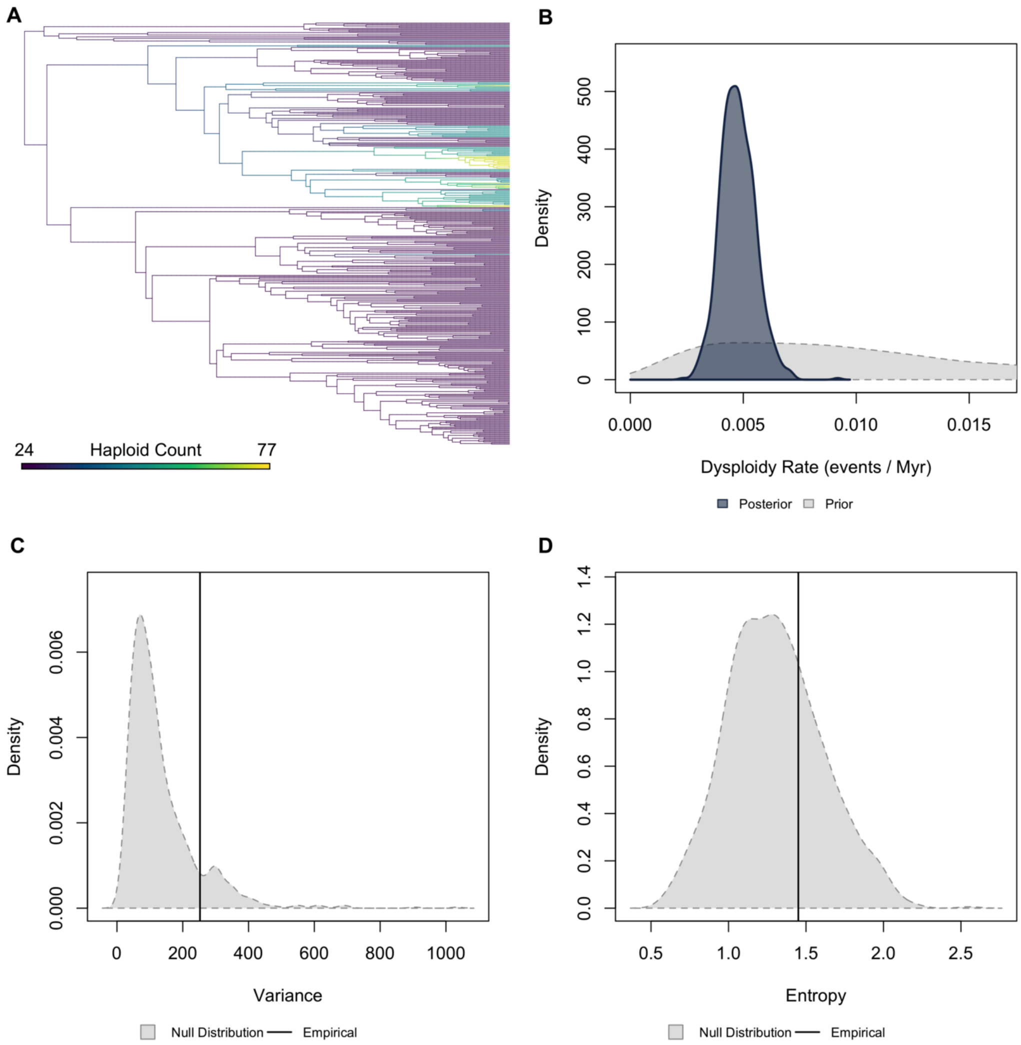


**Figure S59.** ***Validation plots for Cyprinidae.*** *(A) Continuous character map of haploid chromosome number across the phylogeny. (B) Prior versus posterior density distributions for dysploidy rate. (C) Posterior predictive simulation adequacy test for variance. (D) Posterior predictive simulation adequacy test for Shannon's entropy.*

Chondrichthyes | Higher Taxonomy: Chondrichthyes | Chromosomes sampled: 176 | Phylogenetic tips: 1,192 | Overlap with phylogeny: 77 species | Root age: 413 Ma | Taxonomic tips: FALSE | Unresolved: N/A

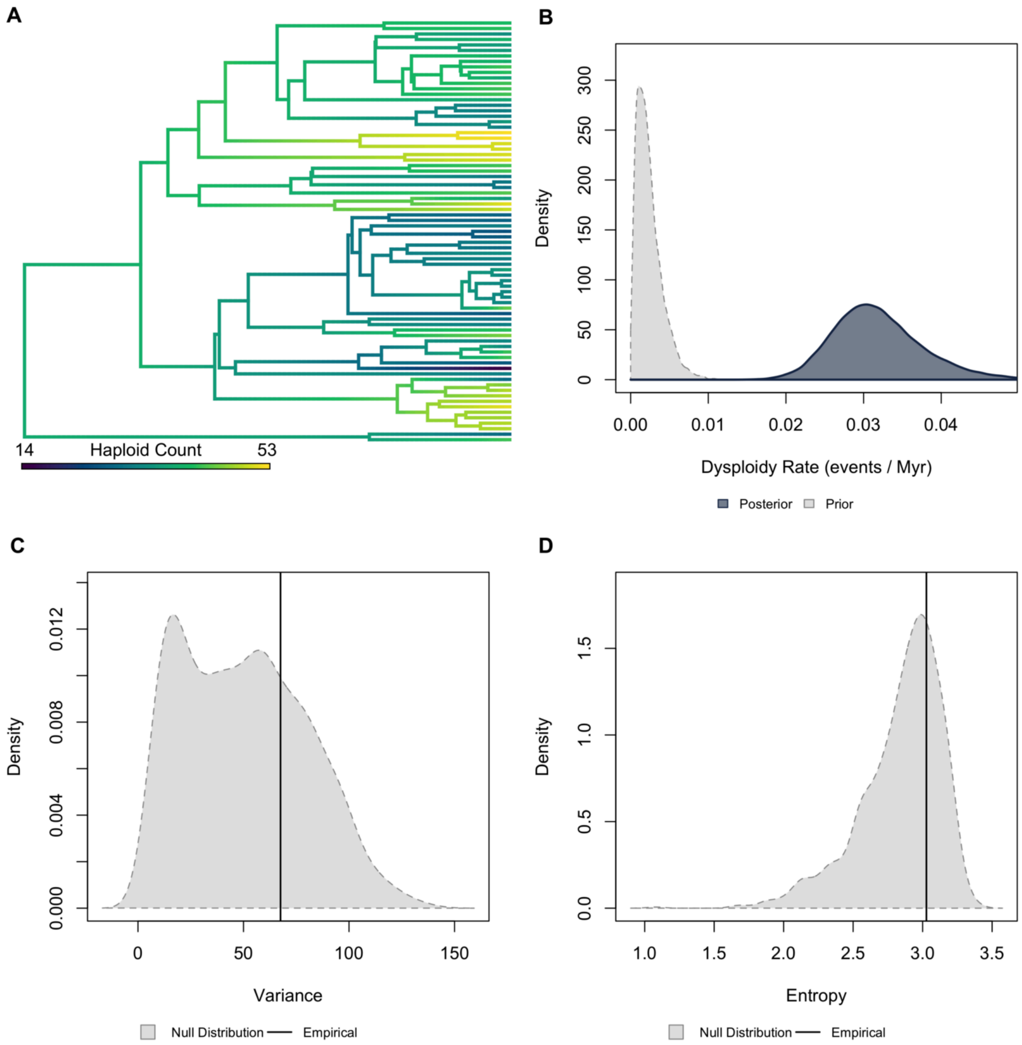


**Figure S60.** ***Validation plots for Chondrichthyes.*** *(A) Continuous character map of haploid chromosome number across the phylogeny. (B) Prior versus posterior density distributions for dysploidy rate. (C) Posterior predictive simulation adequacy test for variance. (D) Posterior predictive simulation adequacy test for Shannon's entropy.*

**Table S4. Summary of PPS adequacy analysis.** False values indicate that the empirical statistic fell within the 95% highest posterior density of the simulated null distribution. In contrast, True values indicate that the empirical statistic was outside of this region. The final column pior overlap describes the proportion of overlap between the prior and posterior distribution.

| Clade | Variance | Entropy | Prior Overlap |
|---|---|---|---|
| Accipitriformes | FALSE | FALSE | 2.8% |
| Anabantiformes | FALSE | FALSE | 15% |
| Anura | FALSE | FALSE | 0% |
| Araneae | FALSE | FALSE | 3.4% |
| Asteraceae | FALSE | TRUE | 0% |
| Blattodea | FALSE | FALSE | 31% |
| Brassicaceae | TRUE | FALSE | 15% |
| Bryophyta | FALSE | FALSE | 0% |
| Carabidae | FALSE | FALSE | 3.5% |
| Carnivora | FALSE | FALSE | 2.5% |
| Caudata | FALSE | TRUE | 43% |
| Cetacea | FALSE | FALSE | 39% |
| Characidae | FALSE | FALSE | 36% |
| Chiroptera | FALSE | FALSE | 2% |
| Chondrichthyes | FALSE | FALSE | 0% |
| Chrysomelidae | TRUE | FALSE | 6.7% |
| Cichlidae | FALSE | FALSE | 35% |
| Coccinellidae | FALSE | FALSE | 53% |
| Cricetidae | FALSE | FALSE | 0.03% |
| Curculionidae | FALSE | FALSE | 68% |
| Cyprinidae | FALSE | FALSE | 25% |
| Drosophilidae | FALSE | FALSE | 44% |
| Dytiscidae | FALSE | FALSE | 33% |
| Fabaceae | TRUE | TRUE | 9.3% |
| Fungi | FALSE | FALSE | 7% |
| Galliformes | FALSE | TRUE | 1.1% |
| Gekkonidae | FALSE | FALSE | 11% |

| Clade | Variance | Entropy | Prior Overlap |
|---|---|---|---|
| Gobiidae | FALSE | FALSE | 51% |
| Gymnospermae | FALSE | FALSE | 14% |
| Hemiptera | FALSE | FALSE | 25% |
| Hydrophilidae | FALSE | FALSE | 63% |
| Hymenoptera | FALSE | FALSE | 0% |
| Iguania | FALSE | FALSE | 3.6% |
| Lepidoptera | TRUE | TRUE | 0% |
| Liliaceae | FALSE | FALSE | 7.6% |
| Magnoliaceae* | TRUE | FALSE | 5.2% |
| Marsupialia | FALSE | FALSE | 68% |
| Muridae | FALSE | FALSE | 2.5% |
| Nothobranchiidae | FALSE | FALSE | 2.8% |
| Odonata | FALSE | FALSE | 20% |
| Orchidaceae | FALSE | TRUE | 0% |
| Orthoptera | FALSE | FALSE | 57% |
| Passeriformes | FALSE | TRUE | 0.17% |
| Passifloraceae | FALSE | FALSE | 47% |
| Phasmatodea | FALSE | FALSE | 40% |
| Primates | FALSE | FALSE | 2.6% |
| Pteridophyta | FALSE | FALSE | 0.08% |
| Rubiaceae | TRUE | TRUE | 15% |
| Scarabidae | FALSE | FALSE | 39% |
| Scincoidea | FALSE | FALSE | 39% |
| Scorpiones | TRUE | TRUE | 8.5% |
| Serpentes | FALSE | FALSE | 7.2% |
| Siluriformes | TRUE | TRUE | 0% |
| Solanaceae | FALSE | FALSE | 21% |
| Tenebrionidae | FALSE | FALSE | 57% |
| Testudines | FALSE | FALSE | 24% |

* Review of the Magnoliacea dataset revealed that the majority of species were included based strictly on taxonomy and we removed this clade from downstream analyses and discussion as we do not believe the rates estimates are robust.

### Tree Resolution Effects on Dysploidy Rates

Phylogenies for plant clades were obtained from the Smith and Brown (2018) seed plant megaphylogeny, which contains unresolved polytomies in several regions of the tree. In the main analyses, these polytomies were resolved using multi2di, producing fully bifurcating trees for downstream modeling. To assess whether this resolution step influenced estimates of dysploidy rates, we conducted a sensitivity analysis in which all tips descending directly from polytomies were removed prior to analysis, thereby avoiding arbitrary resolution of unresolved relationships. Using these pruned trees, we reran all chromosome-number evolution analyses under the same models, priors, and MCMC settings as in the main analysis. Dysploidy rate posteriors from the rerun analyses were then compared to those from the original analyses, both in the context of the full set of clades and through direct comparison of old versus rerun posterior distributions for individual plant families. This approach allowed us to evaluate the extent to which inferred dysploidy rates are sensitive to tree resolution, while preserving comparability with the primary results.

Across the full dataset, updating the plant clade estimates produces little change in the overall placement of dysploidy rates. The relative ordering of major clades remains stable, and broad-scale differences among plants, animals, and fungi are preserved. Within plants, Orchidaceae shows a modest leftward shift in its posterior distribution, but it continues to exhibit the highest dysploidy rates among plant families. In contrast, Solanaceae and Rubiaceae shift slightly to the right, indicating higher inferred rates in the rerun analyses, though these changes are small relative to the global range of dysploidy rates across all clades (Fig. S61).

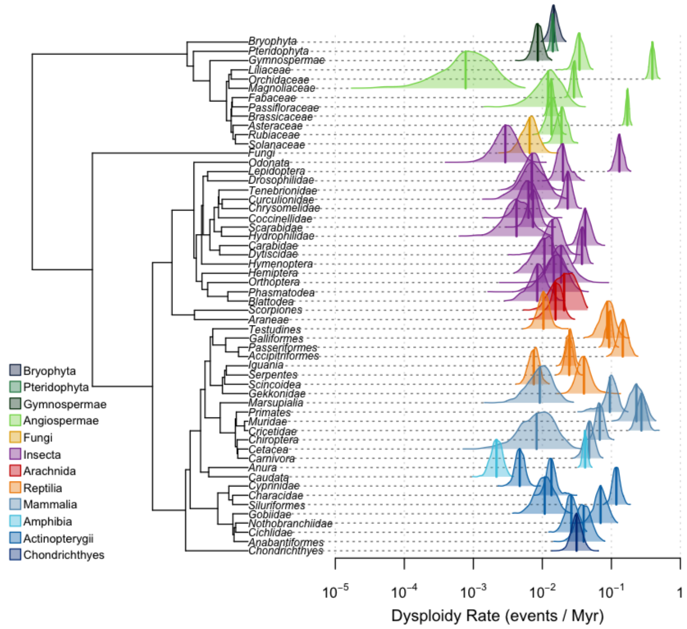


**Figure S61. Tree-wide comparison of dysploidy rate estimates after updating plant clades.** Tree-wide posterior distributions of dysploidy rates across major eukaryotic clades after substituting rerun MCMC analyses for plant families in which phylogenies were modified by pruning tips that directly descended from polytomies.

Direct comparison of posterior dysploidy rate distributions highlights clade-specific responses to the alternative tree treatment. Asteraceae and Brassicaceae show substantial overlap between original and rerun posteriors, with only minor shifts in mean rate estimates, indicating low sensitivity to tree resolution. Fabaceae and Liliaceae exhibit moderate shifts, with rerun posteriors displaced relative to the original analyses but retaining overlapping support. In contrast, Orchidaceae, Solanaceae, and Rubiaceae show pronounced changes, with little overlap between original and rerun posterior distributions. For Orchidaceae, the rerun analysis shifts inferred dysploidy rates toward lower values, whereas Solanaceae and Rubiaceae shift toward higher rates (Fig. S62). Despite these clade-specific differences, the relative placement of these families within the broader distribution of dysploidy rates remains consistent, and large-scale comparative patterns are preserved.

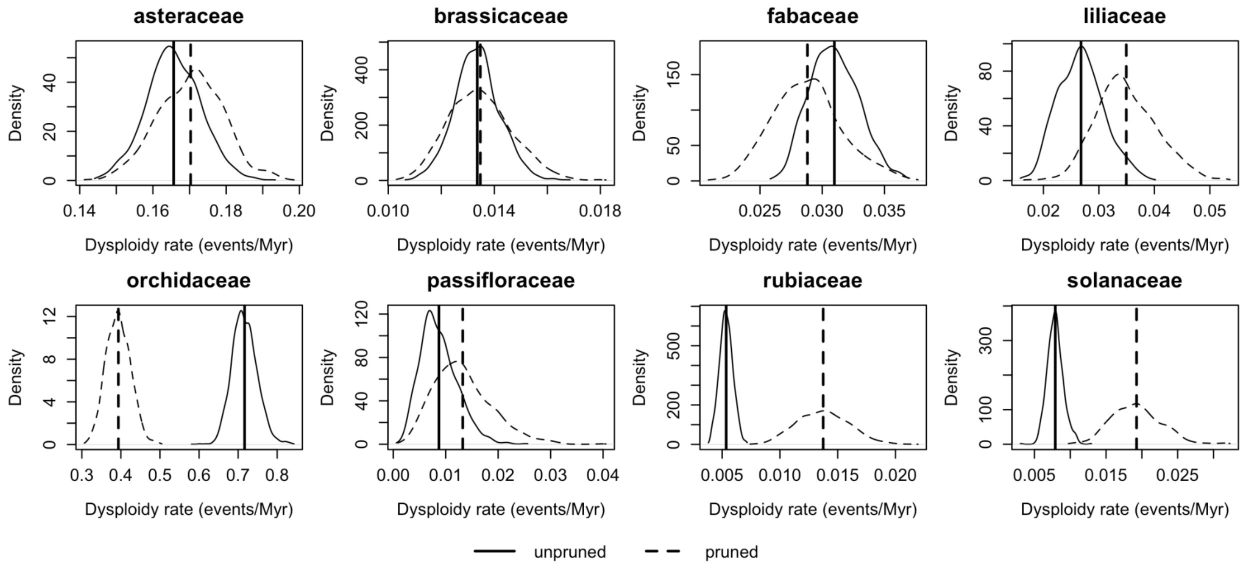


**Figure S62**. **Clade-specific comparison of original and rerun dysploidy rate posteriors.** Density comparison of dysploidy rates for eight plant families comparing original MCMC analyses (solid lines) and rerun analyses based on alternative tree treatments where tips descending from polytomies were removed (dashed lines). Rates are shown (events per Myr), with vertical lines indicating posterior means. Asteraceae and Brassicaceae show substantial posterior overlap, Fabaceae and Liliaceae exhibit moderate shifts, and Orchidaceae, Solanaceae, and Rubiaceae show pronounced shifts with limited overlap. Despite these differences, relative placement of families is consistent with the tree-wide analysis.

### Genome Size Effects on Dysploidy Rates

Genome size data (haploid nuclear DNA content; C-values) were compiled from publicly available databases for animals, plants, and fungi. Animal genome size estimates were obtained from the Animal Genome Size Database, while plant and fungal C-values were sourced from the Plant DNA C-values Database (Royal Botanic Gardens, Kew) and the Fungal Genome Size Database, respectively. All genome sizes were retained in their reported C-value units.

C-value data were matched to the clade names used in this study to retrieve all available genome size entries for focal clades. Analyses were restricted to clades that were present in both datasets and for which genome size data could be readily queried using available taxonomic identifiers, resulting in 48 of the 55 clades being included. All C-value entries associated with each retained clade were converted to megabases and plotted against the corresponding estimated dysploidy rate (Fig. S63A). The broad overlap of genome sizes across clades, coupled with large differences in dysploidy rates, indicates that genome size alone does not predict the tempo of chromosome number evolution. However, to investigate further, we performed a phylogenetic generalized least squares (PGLS) model assuming a Brownian motion correlation structure. This analysis showed that genome size does not predict dysploidy rate (p = 0.41, Fig. S63B).

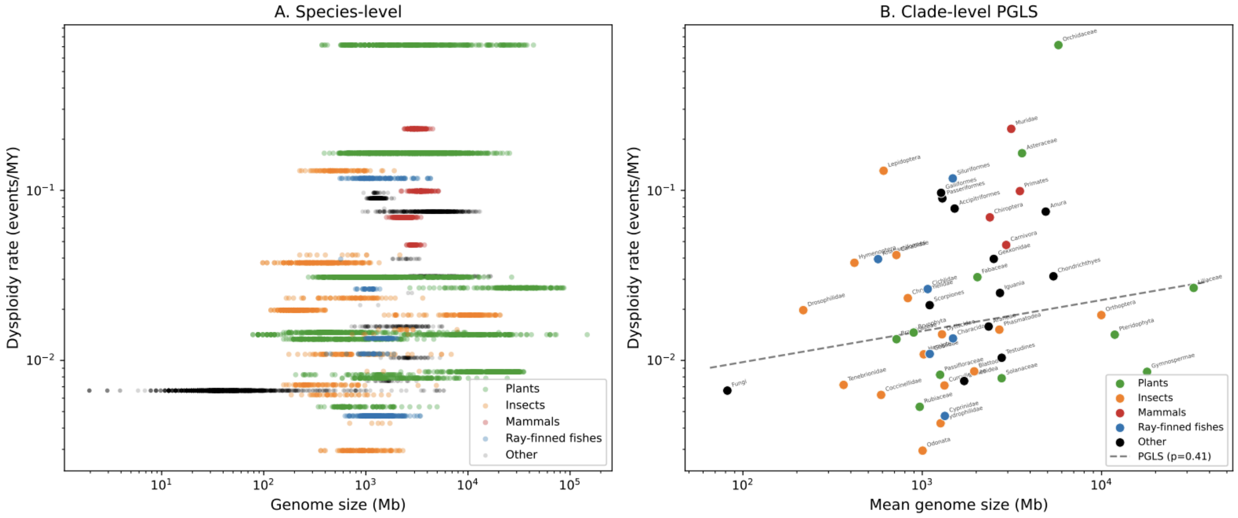


**Figure S63. Genome size and dysploidy rate across taxa at species and clade levels.** A) Species-level relationship between genome size (Mb, log scale) and dysploidy rate (events per million years, log scale). Each point represents a species, colored by major taxonomic groups (plants, insects, mammals, ray-finned fishes, and other taxa). Distributions reveal substantial within-group variation, with plants spanning the widest range of genome sizes and dysploidy rates. B) Clade-level analysis using phylogenetic generalized least squares (PGLS). Points represent clade-level means for genome size and dysploidy rate, labeled by major lineages. The dashed line indicates the fitted PGLS regression ($\rho = 0.41$), showing a non-significant, positive association between genome size and dysploidy rate

#### Variance in Chromosome Number (Plants vs. Animals)

Haploid chromosome number data were compiled for each clade and cleaned to resolve ranges or multiple reported values by randomly sampling a single integer value when necessary. For each clade, the variance in haploid chromosome number was calculated and paired with the clade's median dysploidy rate. Clades were categorized as plants or animals, and distributions of chromosome number variance were compared between these two groups (Fig. S64).

A Spearman rank correlations test revealed no association between dysploidy rate and chromosome number variance in plants ($\rho = 0.12$, $p = 0.734$). In animals, dysploidy rate showed a weak positive trend with chromosome number variance ($\rho = 0.27$, $p = 0.074$), although this relationship was not statistically significant.

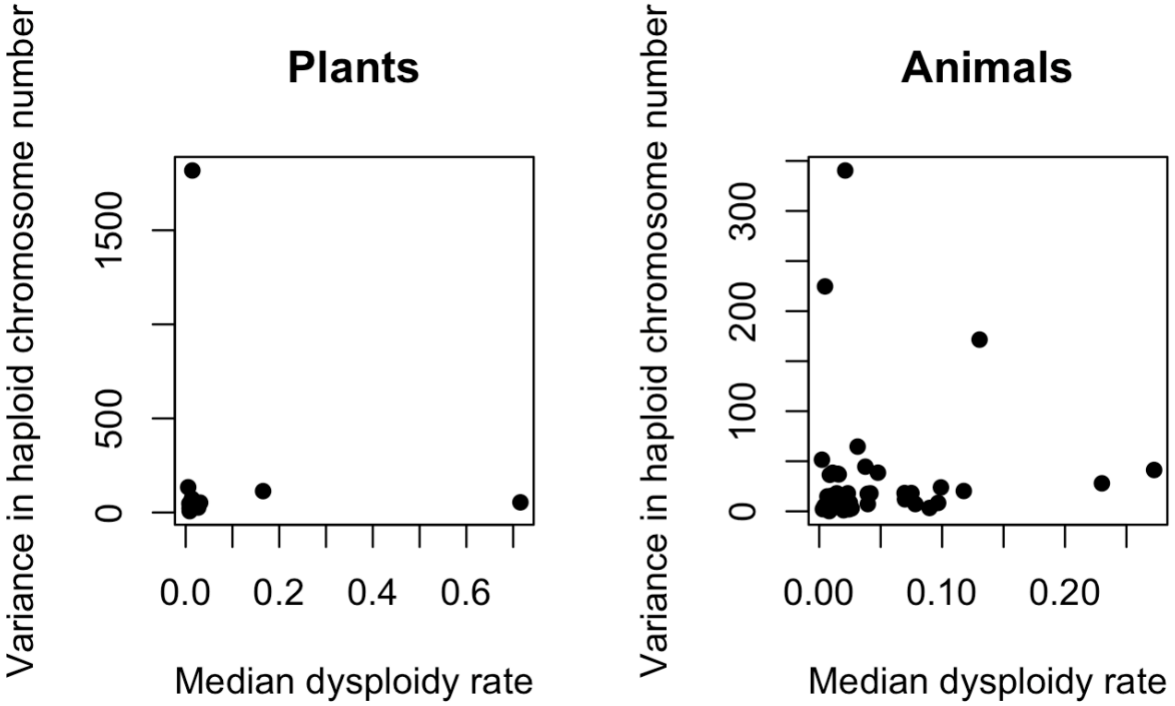


**Figure S64**. **Chromosome number variance in relation to dysploidy rate in plants and animals.** Variance in haploid chromosome number was calculated for each clade using species-level chromosome count data. Each point represents a clade plotted by its median dysploidy rate (horizontal axis) and the variance in haploid chromosome number (vertical axis). Clades were separated into plants and animals and plotted in adjacent panels.

#### Effective Population Size Effects on Dysploidy Rates

Effective population size (Ne) influences the relative strength of selection versus drift and may therefore affect rates of chromosome number change. Because Ne cannot be directly measured for most clades, we used two proxy variables: genome-wide dN/dS ratios (which scale inversely with Ne under nearly neutral theory) and geographic range size (a demographic correlate of census population size).

Genome-wide dN/dS estimates were compiled for 45 of the 56 focal clades from three published sources. For animal clades, values were obtained from Marino et al. (2024), who estimated dN/dS from BUSCO single-copy orthologs across 807 animal species using codon models with phylogenetic filtering. Where a focal clade corresponded directly to a taxonomic order or family represented in that dataset (29 clades), the median dN/dS across all species in that taxon was used. For five beetle families lacking family-level representation, the median across all Coleoptera was used as an order-level proxy. For Orthoptera and Phasmatodea, which belong to orders not represented in Marino et al. (2024), a class-level proxy was calculated as the median across all Insecta species excluding orders that had direct estimates. For Characidae, a superorder-level proxy was computed from all Otophysi species (Siluriformes, Cypriniformes, Characiformes, and Gymnotiformes). Plant dN/dS values were obtained from De La Torre et al. (2017), who estimated genome-wide dN/dS from 42 nuclear gene orthologs: Brassicaceae (0.093) and Fabaceae (0.121) were assigned directly, while remaining angiosperm clades without clade-specific estimates were assigned the mean of these two values (~0.11) as a group-level proxy. Gymnosperm dN/dS (0.314) was taken from Buschiazzo et al. (2012), who compared 968 orthologous gene pairs between *Picea glauca* and *Pinus taeda*. Eleven clades (amphibians, reptiles, arachnids, sharks, and fungi) lacked any published genome-wide dN/dS estimates and were excluded from this analysis. Full provenance for each clade estimate, including species lists and taxonomic matching details, is provided in Table S5.

Geographic range sizes were estimated using occurrence records from the Global Biodiversity Information Facility (GBIF). For each clade, up to 30 species were randomly sampled, and georeferenced occurrence records were retrieved. Range size was approximated as the area of the bounding box enclosing all occurrence points for each species, and the clade-level value was taken as the median across sampled species. Range data were obtained for 55 of the 56 focal clades.

**Table S5. Effective population size proxy data for phylogenetic comparative analyses.** For each of the 56 focal clades, this table reports the median dysploidy rate (events/Myr), genome-wide dN/dS ratio (an inverse proxy for effective population size under nearly neutral theory), and median geographic range size (km²). Genome-wide dN/dS estimates were compiled from three published sources: Marino et al. (2024), who estimated dN/dS from BUSCO single-copy orthologs across animal species (*39*); De La Torre et al. (2017), who estimated dN/dS from 42 nuclear gene orthologs in angiosperms (*40*); and Buschiazzo et al. (2012), who compared orthologous gene pairs in gymnosperms (*41*). For each clade, the table records the number of species contributing to the dN/dS estimate (dNdS_N_Species), the source dataset and taxonomic matching method (dNdS_Source), the estimate quality (direct = clade-level match; order_proxy, class_proxy, or group_proxy = estimate borrowed from a broader taxonomic group; missing = no estimate available), the full literature citation, the species used, and methodological notes. Of the 56 clades, 29 have direct taxonomic matches, 16 have proxy estimates from higher taxonomic levels, and 11 lack published dN/dS data entirely. Geographic range sizes were estimated from GBIF occurrence records for up to 30 randomly sampled species per clade, with range approximated as the bounding-box area enclosing all georeferenced occurrences; range data were obtained for 55 of 56 clades.

| Clade | Higher Classification | Kingdom | Median Dysploidy Rate | Median dNdS | dNdS N Species | dNdS Source | dNdS Quality | Median Range km2 | Range N Species | dNdS_Reference |
|---|---|---|---|---|---|---|---|---|---|---|
| **accipitriformes** | Reptilia | Animal | 0.0781956 | 0.175 | 6 | Marino_order | direct | 28873352 | 29.0 | Marino A, Karagyan G, Hidalgo O, Pellicer J, Barker MS, Leitch IJ. 2024. Genome-wide dN/dS supports an N-centric view of genome size and chromosome evolution. eLife 13:RP98386. DOI: 10.7554/eLife.98386 |
| **anabantiformes** | Actinopterygii | Animal | 0.0394373 | 0.1296 | 4 | Marino_order | direct | 30781705 | 25.0 | Marino A, Karagyan G, Hidalgo O, Pellicer J, Barker MS, Leitch IJ. 2024. Genome-wide dN/dS supports an N-centric view of genome size and chromosome evolution. eLife 13:RP98386. DOI: 10.7554/eLife.98386 |
| **anura** | Amphibia | Animal | 0.0750225 | | | | | 1951053 | 28.0 | |
| **araneae** | Arachnida | Animal | 0.0158285 | | | | | 3141271 | 11.0 | |
| **asteraceae** | Angiosperm | Plant | 0.1654461 | 0.11 | | Angiosperm_average_proxy | group_proxy | 1044223 | 19.0 | De La Torre AR, Li Z, Van de Peer Y, Ingvarsson PK. 2017. Contrasting rates of molecular evolution and patterns of selection among gymnosperms and flowering plants. Molecular Biology and Evolution 34(6):1363-1377. DOI: 10.1093/molbev/msx069 |
| **blattodea** | Insecta | Animal | 0.0086281 | 0.0451 | 3 | Marino_order | direct | 9605702 | 14.0 | Marino A, Karagyan G, Hidalgo O, Pellicer J, Barker MS, Leitch IJ. 2024. Genome-wide dN/dS supports an N-centric view of genome size and chromosome evolution. eLife 13:RP98386. DOI: 10.7554/eLife.98386 |
| **brassicaceae** | Angiosperm | Plant | 0.0133323 | 0.0925 | 42 | DeLaTorre2017 | direct | 3013412 | 12.0 | De La Torre AR, Li Z, Van de Peer Y, Ingvarsson PK. 2017. Contrasting rates of molecular evolution and patterns of selection among gymnosperms and flowering plants. Molecular Biology and Evolution 34(6):1363-1377. DOI: 10.1093/molbev/msx069 |

| Clade | Higher Classification | Kingdom | Median Dysploidy Rate | Median dNdS | dNdS N Species | dNdS Source | dNdS Quality | Median Range km2 | Range N Species | dNdS_Reference |
|---|---|---|---|---|---|---|---|---|---|---|
| **bryophyta** | Bryophyta | Plant | 0.0145887 | 0.11 | 0 | Bryophyte_lit_proxy | group_proxy | 35873284 | 20.0 | Szövényi P, Rensing SA, Lang D, Wray GA, Shaw AJ. 2011. Generation-time effect on genomes of haploid-dominant land plants. Journal of Evolutionary Biology 24(3):666-672. DOI: 10.1111/j.1420-9101.2010.02195.x |
| **carabidae** | Insecta | Animal | 0.0416781 | 0.0431 | 17 | Marino_Coleoptera_proxy | order_proxy | 10813253 | 13.0 | Marino A, Karagyan G, Hidalgo O, Pellicer J, Barker MS, Leitch IJ. 2024. Genome-wide dN/dS supports an N-centric view of genome size and chromosome evolution. eLife 13:RP98386. DOI: 10.7554/eLife.98386 |
| **carnivora** | Mammalia | Animal | 0.0477485 | 0.19 | 41 | Marino_order | direct | 34269856 | 16.0 | Marino A, Karagyan G, Hidalgo O, Pellicer J, Barker MS, Leitch IJ. 2024. Genome-wide dN/dS supports an N-centric view of genome size and chromosome evolution. eLife 13:RP98386. DOI: 10.7554/eLife.98386 |
| **caudata** | Amphibia | Animal | 0.0021749 | | 0 | MISSING | missing | 338909 | 28.0 | |
| **cetacea** | Mammalia | Animal | 0.0081549 | 0.2284 | 7 | Marino_cetacean_families | direct | 111638763 | 28.0 | Marino A, Karagyan G, Hidalgo O, Pellicer J, Barker MS, Leitch IJ. 2024. Genome-wide dN/dS supports an N-centric view of genome size and chromosome evolution. eLife 13:RP98386. DOI: 10.7554/eLife.98386 |
| **characidae** | Actinopterygii | Animal | 0.0134715 | 0.1033 | 21 | Marino_Otophysi_proxy | order_proxy | 7076810 | 29.0 | Marino A, Karagyan G, Hidalgo O, Pellicer J, Barker MS, Leitch IJ. 2024. Genome-wide dN/dS supports an N-centric view of genome size and chromosome evolution. eLife 13:RP98386. DOI: 10.7554/eLife.98386 |
| **chiroptera** | Mammalia | Animal | 0.0694322 | 0.1462 | 19 | Marino_order | direct | 13454520 | 26.0 | Marino A, Karagyan G, Hidalgo O, Pellicer J, Barker MS, Leitch IJ. 2024. Genome-wide dN/dS supports an N-centric view of genome size and chromosome evolution. eLife 13:RP98386. DOI: 10.7554/eLife.98386 |
| **chondrichthyes** | Chondrichthyes | Animal | 0.0312500 | | 0 | MISSING | missing | 18243305 | 27.0 | |
| **chrysomelidae** | Insecta | Animal | 0.0232578 | 0.0421 | 2 | Marino_family | direct | 4987157 | 14.0 | Marino A, Karagyan G, Hidalgo O, Pellicer J, Barker MS, Leitch IJ. 2024. Genome-wide dN/dS supports an N-centric view of genome size and chromosome evolution. eLife 13:RP98386. DOI: 10.7554/eLife.98386 |
| **cichlidae** | Actinopterygii | Animal | 0.0262911 | 0.1809 | 5 | Marino_family | direct | 21129809 | 26.0 | Marino A, Karagyan G, Hidalgo O, Pellicer J, Barker MS, Leitch IJ. 2024. Genome-wide dN/dS supports an N-centric view of genome size and chromosome evolution. eLife 13:RP98386. DOI: 10.7554/eLife.98386 |
| **coccinellidae** | Insecta | Animal | 0.0062698 | 0.0431 | 3 | Marino_family | direct | | | Marino A, Karagyan G, Hidalgo O, Pellicer J, Barker MS, Leitch IJ. 2024. Genome-wide dN/dS supports an N-centric view of genome size and chromosome evolution. eLife 13:RP98386. DOI: 10.7554/eLife.98386 |
| **cricetidae** | Mammalia | Animal | 0.2725445 | 0.1534 | 11 | Marino_family | direct | 1314348 | 12.0 | Marino A, Karagyan G, Hidalgo O, Pellicer J, Barker MS, Leitch IJ. 2024. Genome-wide dN/dS supports an N-centric |

| Clade | Higher Classification | Kingdom | Median Dysploidy Rate | Median dNdS | dNdS N Species | dNdS Source | dNdS Quality | Median Range km2 | Range N Species | dNdS_Reference |
|---|---|---|---|---|---|---|---|---|---|---|
| | | | | | | | | | | view of genome size and chromosome evolution. eLife 13:RP98386. DOI: 10.7554/eLife.98386 |
| **curculionidae** | Insecta | Animal | 0.0071242 | 0.0416 | 2 | Marino_family | direct | 60394120 | 2.0 | Marino A, Karagyan G, Hidalgo O, Pellicer J, Barker MS, Leitch IJ. 2024. Genome-wide dN/dS supports an N-centric view of genome size and chromosome evolution. eLife 13:RP98386. DOI: 10.7554/eLife.98386 |
| **cyprinidae** | Actinopterygii | Animal | 0.0047136 | 0.1289 | 2 | Marino_family | direct | 1401833 | 13.0 | Marino A, Karagyan G, Hidalgo O, Pellicer J, Barker MS, Leitch IJ. 2024. Genome-wide dN/dS supports an N-centric view of genome size and chromosome evolution. eLife 13:RP98386. DOI: 10.7554/eLife.98386 |
| **drosophilidae** | Insecta | Animal | 0.0197546 | 0.0444 | 46 | Marino_family | direct | 40524430 | 5.0 | Marino A, Karagyan G, Hidalgo O, Pellicer J, Barker MS, Leitch IJ. 2024. Genome-wide dN/dS supports an N-centric view of genome size and chromosome evolution. eLife 13:RP98386. DOI: 10.7554/eLife.98386 |
| **dytiscidae** | Insecta | Animal | 0.0142444 | 0.0431 | 17 | Marino_Coleoptera_proxy | order_proxy | 9999422 | 14.0 | Marino A, Karagyan G, Hidalgo O, Pellicer J, Barker MS, Leitch IJ. 2024. Genome-wide dN/dS supports an N-centric view of genome size and chromosome evolution. eLife 13:RP98386. DOI: 10.7554/eLife.98386 |
| **fabaceae** | Angiosperm | Plant | 0.0308889 | 0.121 | 42 | DeLaTorre2017 | direct | 16471919 | 27.0 | De La Torre AR, Li Z, Van de Peer Y, Ingvarsson PK. 2017. Contrasting rates of molecular evolution and patterns of selection among gymnosperms and flowering plants. Molecular Biology and Evolution 34(6):1363-1377. DOI: 10.1093/molbev/msx069 |
| **fungi** | Fungi | Fungi | 0.0066463 | | 0 | MISSING | missing | 294164103 | 12.0 | |
| **galliformes** | Reptilia | Animal | 0.0967372 | 0.1398 | 9 | Marino_order | direct | 8159848 | 30.0 | Marino A, Karagyan G, Hidalgo O, Pellicer J, Barker MS, Leitch IJ. 2024. Genome-wide dN/dS supports an N-centric view of genome size and chromosome evolution. eLife 13:RP98386. DOI: 10.7554/eLife.98386 |
| **gekkonidae** | Reptilia | Animal | 0.0395387 | | 0 | MISSING | missing | 3370143 | 27.0 | |
| **gobiidae** | Actinopterygii | Animal | 0.0109226 | 0.0881 | 3 | Marino_family | direct | 7199706 | 24.0 | Marino A, Karagyan G, Hidalgo O, Pellicer J, Barker MS, Leitch IJ. 2024. Genome-wide dN/dS supports an N-centric view of genome size and chromosome evolution. eLife 13:RP98386. DOI: 10.7554/eLife.98386 |
| **gymnospermae** | Gymnosperms | Plant | 0.0085692 | 0.314 | 42 | Buschiazzo2012 | direct | 35295486 | 27.0 | Buschiazzo E, Ritland C, Bohlmann J, Ritland K. 2012. Slow but not low: genomic comparisons reveal slower evolutionary rate and higher dN/dS in conifers compared to angiosperms. BMC Evolutionary Biology 12:8. DOI: 10.1186/1471-2148-12-8 |
| **hemiptera** | Insecta | Animal | 0.0108603 | 0.0498 | 16 | Marino_order | direct | 12178187 | 17.0 | Marino A, Karagyan G, Hidalgo O, Pellicer J, Barker MS, Leitch IJ. 2024. Genome-wide dN/dS supports an N-centric |

| Clade | Higher Classification | Kingdom | Median Dysploidy Rate | Median dNdS | dNdS N Species | dNdS Source | dNdS Quality | Median Range km2 | Range N Species | dNdS_Reference |
|---|---|---|---|---|---|---|---|---|---|---|
| | | | | | | | | | | view of genome size and chromosome evolution. eLife 13:RP98386. DOI: 10.7554/eLife.98386 |
| **hydrophilidae** | Insecta | Animal | 0.0042701 | 0.0431 | 17 | Marino_Coleoptera_proxy | order_proxy | 5196294 | 17.0 | Marino A, Karagyan G, Hidalgo O, Pellicer J, Barker MS, Leitch IJ. 2024. Genome-wide dN/dS supports an N-centric view of genome size and chromosome evolution. eLife 13:RP98386. DOI: 10.7554/eLife.98386 |
| **hymenoptera** | Insecta | Animal | 0.0375153 | 0.0911 | 43 | Marino_order | direct | 19798909 | 8.0 | Marino A, Karagyan G, Hidalgo O, Pellicer J, Barker MS, Leitch IJ. 2024. Genome-wide dN/dS supports an N-centric view of genome size and chromosome evolution. eLife 13:RP98386. DOI: 10.7554/eLife.98386 |
| **iguania** | Reptilia | Animal | 0.0249282 | | 0 | MISSING | missing | 1120187 | 28.0 | |
| **lepidoptera** | Insecta | Animal | 0.1304819 | 0.0444 | 39 | Marino_order | direct | 9942894 | 19.0 | Marino A, Karagyan G, Hidalgo O, Pellicer J, Barker MS, Leitch IJ. 2024. Genome-wide dN/dS supports an N-centric view of genome size and chromosome evolution. eLife 13:RP98386. DOI: 10.7554/eLife.98386 |
| **liliaceae** | Angiosperm | Plant | 0.0266970 | 0.11 | 0 | Angiosperm_average_proxy | group_proxy | 234163 | 24.0 | De La Torre AR, Li Z, Van de Peer Y, Ingvarsson PK. 2017. Contrasting rates of molecular evolution and patterns of selection among gymnosperms and flowering plants. Molecular Biology and Evolution 34(6):1363-1377. DOI: 10.1093/molbev/msx069 |
| **magnoliaceae** | Angiosperm | Plant | 0.0008477 | 0.11 | 0 | Angiosperm_average_proxy | group_proxy | 55092155 | 12.0 | De La Torre AR, Li Z, Van de Peer Y, Ingvarsson PK. 2017. Contrasting rates of molecular evolution and patterns of selection among gymnosperms and flowering plants. Molecular Biology and Evolution 34(6):1363-1377. DOI: 10.1093/molbev/msx069 |
| **marsupialia** | Mammalia | Animal | 0.0094438 | 0.1142 | 3 | Marino_marsupial_orders | direct | 2528431 | 29.0 | Marino A, Karagyan G, Hidalgo O, Pellicer J, Barker MS, Leitch IJ. 2024. Genome-wide dN/dS supports an N-centric view of genome size and chromosome evolution. eLife 13:RP98386. DOI: 10.7554/eLife.98386 |
| **muridae** | Mammalia | Animal | 0.2300528 | 0.1256 | 4 | Marino_family | direct | 12921671 | 20.0 | Marino A, Karagyan G, Hidalgo O, Pellicer J, Barker MS, Leitch IJ. 2024. Genome-wide dN/dS supports an N-centric view of genome size and chromosome evolution. eLife 13:RP98386. DOI: 10.7554/eLife.98386 |
| **nothobranchiidae** | Actinopterygii | Animal | 0.0697547 | 0.1204 | 2 | Marino_family | direct | 236318 | 14.0 | Marino A, Karagyan G, Hidalgo O, Pellicer J, Barker MS, Leitch IJ. 2024. Genome-wide dN/dS supports an N-centric view of genome size and chromosome evolution. eLife 13:RP98386. DOI: 10.7554/eLife.98386 |
| **odonata** | Insecta | Animal | 0.0029479 | 0.0909 | 2 | Marino_order | direct | 4859271 | 20.0 | Marino A, Karagyan G, Hidalgo O, Pellicer J, Barker MS, Leitch IJ. 2024. Genome-wide dN/dS supports an N-centric view of genome size and chromosome evolution. eLife 13:RP98386. DOI: 10.7554/eLife.98386 |

| Clade | Higher Classification | Kingdom | Median Dysploidy Rate | Median dNdS | dNdS N Species | dNdS Source | dNdS Quality | Median Range km2 | Range N Species | dNdS_Reference |
|---|---|---|---|---|---|---|---|---|---|---|
| **orchidaceae** | Angiosperm | Plant | 0.7155285 | 0.11 | 0 | Angiosperm_average_proxy | group_proxy | 7059869 | 13.0 | De La Torre AR, Li Z, Van de Peer Y, Ingvarsson PK. 2017. Contrasting rates of molecular evolution and patterns of selection among gymnosperms and flowering plants. Molecular Biology and Evolution 34(6):1363-1377. DOI: 10.1093/molbev/msx069 |
| **orthoptera** | Insecta | Animal | 0.0184776 | 0.0444 | 86 | Marino_Insecta_proxy | class_proxy | 3759408 | 8.0 | Marino A, Karagyan G, Hidalgo O, Pellicer J, Barker MS, Leitch IJ. 2024. Genome-wide dN/dS supports an N-centric view of genome size and chromosome evolution. eLife 13:RP98386. DOI: 10.7554/eLife.98386 |
| **passeriformes** | Reptilia | Animal | 0.0897500 | 0.152 | 138 | Marino_order | direct | 27608488 | 26.0 | Marino A, Karagyan G, Hidalgo O, Pellicer J, Barker MS, Leitch IJ. 2024. Genome-wide dN/dS supports an N-centric view of genome size and chromosome evolution. eLife 13:RP98386. DOI: 10.7554/eLife.98386 |
| **passifloraceae** | Angiosperm | Plant | 0.0082359 | 0.11 | 0 | Angiosperm_average_proxy | group_proxy | 75301309 | 25.0 | De La Torre AR, Li Z, Van de Peer Y, Ingvarsson PK. 2017. Contrasting rates of molecular evolution and patterns of selection among gymnosperms and flowering plants. Molecular Biology and Evolution 34(6):1363-1377. DOI: 10.1093/molbev/msx069 |
| **phasmatodea** | Insecta | Animal | 0.0151691 | 0.0444 | 86 | Marino_Insecta_proxy | class_proxy | 1155384 | 11.0 | Marino A, Karagyan G, Hidalgo O, Pellicer J, Barker MS, Leitch IJ. 2024. Genome-wide dN/dS supports an N-centric view of genome size and chromosome evolution. eLife 13:RP98386. DOI: 10.7554/eLife.98386 |
| **primates** | Mammalia | Animal | 0.0990239 | 0.2162 | 25 | Marino_order | direct | 62693686 | 14.0 | Marino A, Karagyan G, Hidalgo O, Pellicer J, Barker MS, Leitch IJ. 2024. Genome-wide dN/dS supports an N-centric view of genome size and chromosome evolution. eLife 13:RP98386. DOI: 10.7554/eLife.98386 |
| **pteridophyta** | Pteridophytes | Plant | 0.0141757 | 0.11 | 0 | Pteridophyte_lit_proxy | group_proxy | 11907040 | 18.0 | Barker MS, Wolf PG. 2010. Unfurling fern biology in the genomics age. BioScience 60(3):177-185. DOI: 10.1525/bio.2010.60.3.4 |
| **rubiaceae** | Angiosperm | Plant | 0.0053351 | 0.11 | 0 | Angiosperm_average_proxy | group_proxy | 7025084 | 7.0 | De La Torre AR, Li Z, Van de Peer Y, Ingvarsson PK. 2017. Contrasting rates of molecular evolution and patterns of selection among gymnosperms and flowering plants. Molecular Biology and Evolution 34(6):1363-1377. DOI: 10.1093/molbev/msx069 |
| **scarabidae** | Insecta | Animal | 0.0073315 | 0.0542 | 4 | Marino_scarabaeidae | direct | 10268851 | 20.0 | Marino A, Karagyan G, Hidalgo O, Pellicer J, Barker MS, Leitch IJ. 2024. Genome-wide dN/dS supports an N-centric view of genome size and chromosome evolution. eLife 13:RP98386. DOI: 10.7554/eLife.98386 |
| **scincoidea** | Reptilia | Animal | 0.0075668 | | 0 | MISSING | missing | 881114 | 27.0 | |
| **scorpiones** | Arachnida | Animal | 0.0211364 | | 0 | MISSING | missing | 2649082 | 12.0 | |

| Clade | Higher Classification | Kingdom | Median Dysploidy Rate | Median dNdS | dNdS N Species | dNdS Source | dNdS Quality | Median Range km2 | Range N Species | dNdS_Reference |
|---|---|---|---|---|---|---|---|---|---|---|
| **serpentes** | Reptilia | Animal | 0.0239865 | | 0 | MISSING | missing | 4976397 | 28.0 | |
| **siluriformes** | Actinopterygii | Animal | 0.1176281 | 0.1297 | 5 | Marino_order | direct | 2389470 | 25.0 | Marino A, Karagyan G, Hidalgo O, Pellicer J, Barker MS, Leitch IJ. 2024. Genome-wide dN/dS supports an N-centric view of genome size and chromosome evolution. eLife 13:RP98386. DOI: 10.7554/eLife.98386 |
| **solanaceae** | Angiosperm | Plant | 0.0078591 | 0.11 | 0 | Angiosperm_average_proxy | group_proxy | 16335592 | 14.0 | De La Torre AR, Li Z, Van de Peer Y, Ingvarsson PK. 2017. Contrasting rates of molecular evolution and patterns of selection among gymnosperms and flowering plants. Molecular Biology and Evolution 34(6):1363-1377. DOI: 10.1093/molbev/msx069 |
| **tenebrionidae** | Insecta | Animal | 0.0071796 | 0.0431 | 17 | Marino_Coleoptera_proxy | order_proxy | 436135 | 3.0 | Marino A, Karagyan G, Hidalgo O, Pellicer J, Barker MS, Leitch IJ. 2024. Genome-wide dN/dS supports an N-centric view of genome size and chromosome evolution. eLife 13:RP98386. DOI: 10.7554/eLife.98386 |
| **testudines** | Reptilia | Animal | 0.0103661 | | 0 | MISSING | missing | 6636177 | 22.0 | |

We tested the relationship between each Ne proxy and dysploidy rate using phylogenetic generalized least squares (PGLS) implemented in the R package caper (*42*). Dysploidy rates and both predictor variables were log10-transformed prior to analysis. A 56-tip ultrametric phylogeny was constructed from published divergence time estimates (*43, 44*); for the focal clades. Pagel's lambda was estimated by maximum likelihood, with bounds set to [0.001, 1.0].

Among animals only (N = 26 clades with direct or order-level dN/dS estimates), dN/dS showed a positive but marginally non-significant relationship with dysploidy rate (slope = 0.871, p = 0.059, R2 = 0.147, lambda = 0.0; Fig. S65). This relationship became significant when Lepidoptera--an outlier with an exceptionally low dN/dS relative to its dysploidy rate--was excluded (slope = 1.19, p = 0.015, R2 = 0.242). Across all clades with any dN/dS estimate (N = 29, including plant proxies), the relationship was weaker (slope = 0.606, p = 0.149). Geographic range size showed no significant relationship with dysploidy rate (N = 55, p = 0.293; Fig. S66). Leave-one-out sensitivity analysis confirmed that no single animal clade drove the dN/dS result (Fig. S67), and lambda profile likelihood plots indicated negligible phylogenetic signal in the residuals (Fig. S68). Variance inflation factors for the two predictors were low (VIF = 1.02), confirming that dN/dS and range size capture largely independent axes of variation. Full PGLS results and sensitivity analyses are reported in Figures S65-S68 and Table S5.

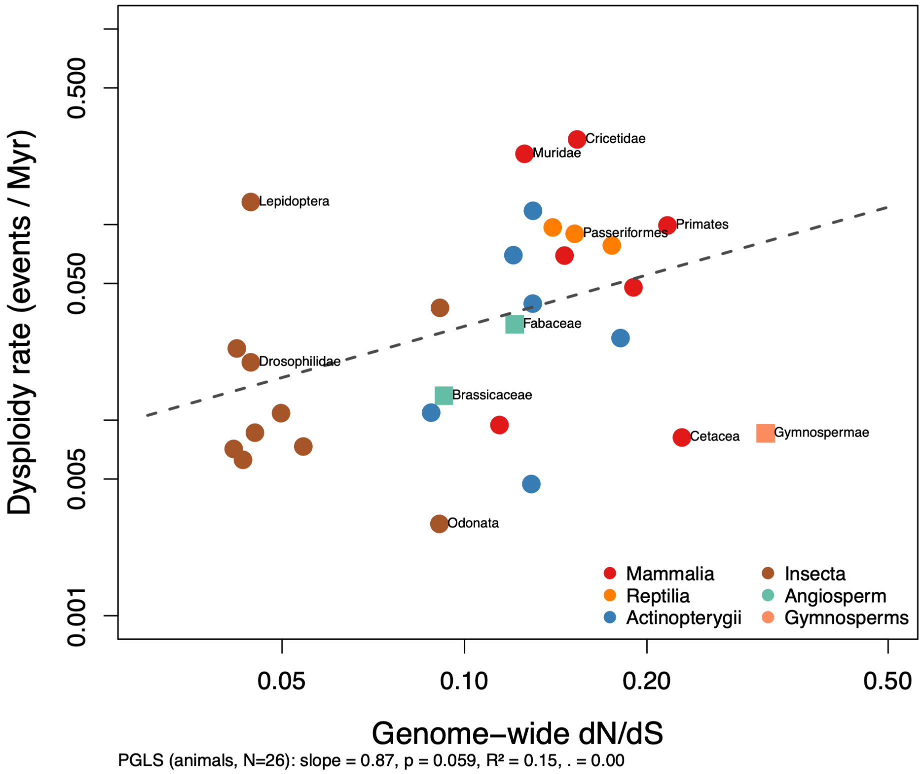


**Figure S65. Genome-wide dN/dS versus dysploidy rate.** Relationship between log10-transformed genome-wide dN/dS and log10-transformed dysploidy rate for animal clades with direct taxonomic matches to the Marino et al. (2024) dataset. The blue line shows the PGLS regression (slope = 0.871, p = 0.059). Higher dN/dS values (indicating smaller effective population sizes) are associated with faster rates of chromosome number evolution. Point labels identify individual clades.

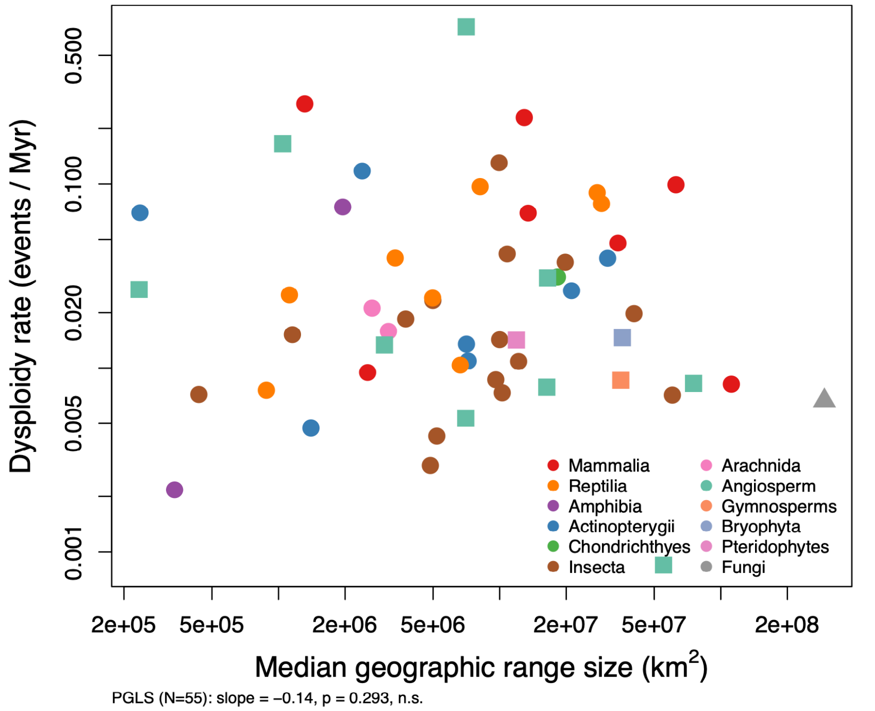


**Figure S66. Geographic range size versus dysploidy rate.** Relationship between log10-transformed median geographic range size (km2) and log10-transformed dysploidy rate across 55 eukaryotic clades. The PGLS regression line is shown ($p = 0.293$, n.s.). Points are colored by kingdom. Range size does not predict dysploidy rate across the sampled clades.

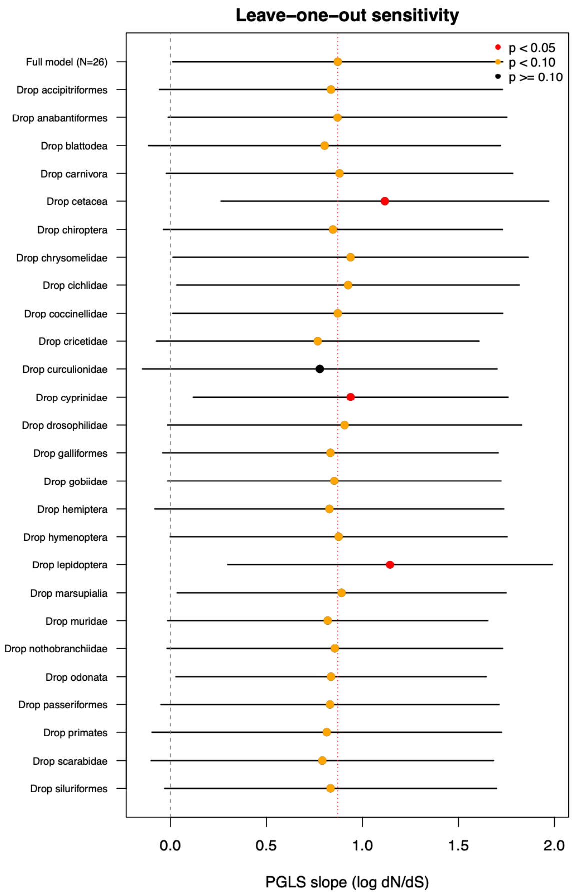


**Figure S67. Leave-one-out sensitivity analysis for dN/dS versus dysploidy rate.** Forest plot showing PGLS slope estimates and 95% confidence intervals when each animal clade is sequentially excluded from the analysis. No single clade exclusion qualitatively changes the result, indicating that the marginal relationship is not driven by any individual outlier.

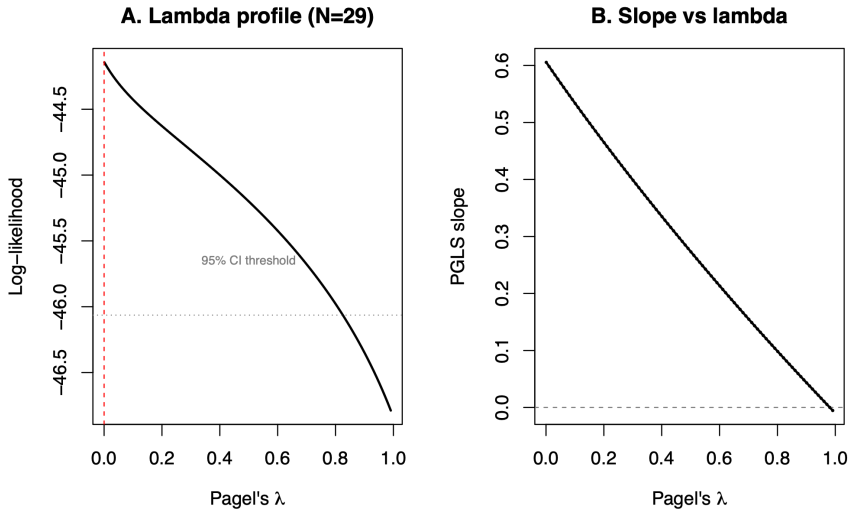


**Figure S68. Lambda profile likelihood for PGLS residuals.** Profile likelihood surface for Pagel’s lambda estimated from the PGLS residuals of log10(dysploidy rate) on log10(dN/dS) across 26 animal clades. The maximum likelihood estimate of lambda = 0.0 indicates negligible phylogenetic signal in the residuals, supporting the use of a star phylogeny error structure and suggesting that the relationship between dN/dS and dysploidy rate is not confounded by shared ancestry among clades.

#### Variance Decomposition: Within- vs. Between-Group Variation in Dysploidy Rates

A central claim of this study is that the tempo of chromosomal evolution is not partitioned by major phylogenetic divisions. To formally test this, we performed a hierarchical variance decomposition of $\log_{10}$-transformed median dysploidy rates across the 55 focal clades (excluding Magnoliaceae), partitioning variance at two taxonomic levels: kingdom (Animal, Plant, Fungi; 3 groups) and higher taxonomic classification (Mammalia, Insecta, Reptilia, Angiospermae, etc.; 12 groups).

At each level, we estimated variance components using three complementary approaches: (1) classical one-way ANOVA with unbalanced variance component estimation, (2) restricted maximum likelihood (REML) via a random-intercept model implemented in lme4, and (3) a permutation test (9,999 permutations) to assess the significance of the observed F-statistic without distributional assumptions. For the kingdom-level contrast, we additionally tested the effect of kingdom membership using phylogenetic generalized least squares (PGLS) with Pagel's lambda estimated by maximum likelihood (caper), fitting a binary plant-versus-animal predictor on the 54-clade ultrametric phylogeny (excluding Fungi, n = 1). Finally, we fit a nested variance components model (Higher Classification within Kingdom) to simultaneously estimate the proportion of variance attributable to kingdom, higher taxonomy, and residual (among-clade) variation.

**Kingdom level.** Kingdom membership explained none of the observed variance in dysploidy rates. The geometric mean dysploidy rate for animals (0.023 events/Myr, n = 43) and plants (0.023 events/Myr, n = 11) were nearly identical, differing by only 1.0-fold, whereas rates within each kingdom spanned 125-fold (animals: caudata 0.002 to Cricetidae 0.27 events/Myr) and 134-fold (plants: rubiaceae 0.005 to Orchidaceae 0.72 events/Myr). The ANOVA was non-significant ($F_{2,52}$ = 0.48, p = 0.62), the REML variance component for kingdom was estimated at zero (ICC = 0.000), and the permutation test confirmed this result (p = 0.64). The PGLS analysis likewise found no effect of kingdom on dysploidy rate ($R^2$ = 0.003, p = 0.75, $\lambda$ = 0.19).

**Higher taxonomic classification level.** At the finer level of higher taxonomy, group identity explained a small fraction of the variance. The ANOVA was non-significant ($F_{11,43}$ = 0.90, p = 0.54; permutation p = 0.53), but the REML model estimated a non-zero Higher Classification variance component corresponding to an ICC of 0.062, indicating that approximately 6.2% of the variance in dysploidy rates is attributable to differences among higher taxonomic groups, with the remaining 93.8% residing among clades within groups. Notably, Mammalia exhibited the highest group mean (−1.26 on the $\log_{10}$ scale), but the within-group standard deviation (0.61) exceeded the difference between the Mammalia mean and the grand mean (0.42), indicating substantial overlap with other groups.

**Nested model.** The nested variance components model partitioning variance into kingdom, higher classification (within kingdom), and residual components confirmed that 93.8% of variance is residual (among clades within higher taxonomic groups), 6.2% is attributable to higher classification, and 0.0% to kingdom (Fig. S69).

These results provide formal statistical support for the claim that the tempo of karyotype evolution is not determined by deep phylogenetic affiliation. The vast majority of variance in dysploidy rates is distributed among clades within taxonomic groups at every level of the hierarchy examined, consistent with the interpretation that lineage-specific biological traits and ecological context (rather than shared evolutionary history) govern the pace of chromosomal change.

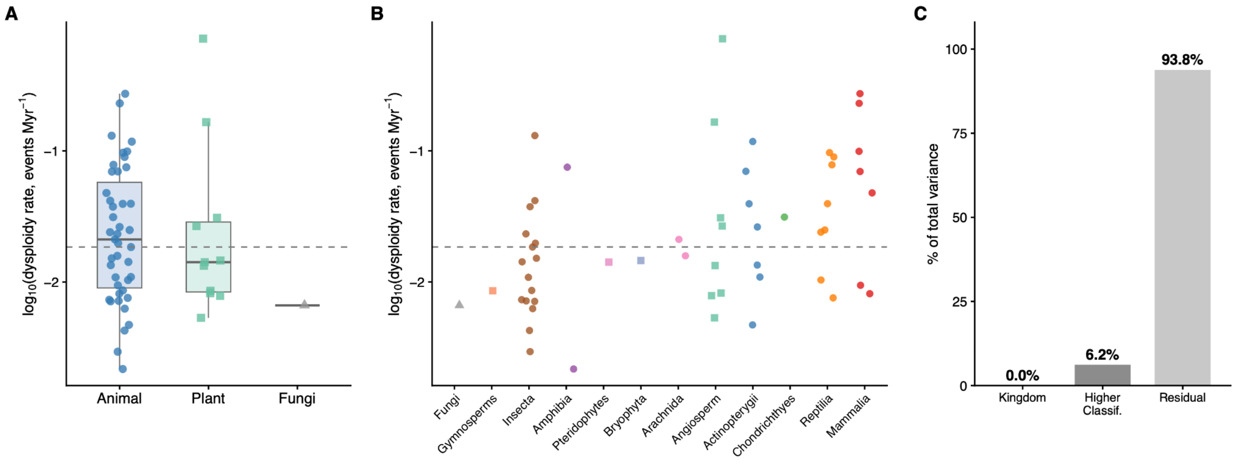


**Figure S69. Variance decomposition of dysploidy rates across taxonomic levels.** (A) Distribution of $\log_{10}$-transformed median dysploidy rates by kingdom (Animal, Plant, Fungi). Points represent individual clades; boxes show interquartile ranges. The dashed line indicates the global median. (B) Distribution of $\log_{10}$-transformed median dysploidy rates by higher taxonomic classification, ordered by group median. Colors indicate kingdom membership (blue = Animal, green = Plant, purple = Fungi). (C) Proportion of total variance in dysploidy rates attributable to kingdom (0.0%), higher taxonomic classification (6.2%), and residual within-group variation (93.8%), estimated from a nested REML variance components model. Magnoliaceae excluded from all analyses (N = 55 clades).

#### Online Database

To make our data more broadly available and easier to access all data included in this paper has been included in an online database that allows users to subset data by clade and download any portion they would like to as CSV file that can be used in comparative software to replicate the analyses included in this manuscript (https://coleoguy.github.io/cures-karyotype-database.html).
